# A transcriptional atlas of early Arabidopsis seed development suggests mechanisms for inter-tissue coordination

**DOI:** 10.1101/2025.04.29.651360

**Authors:** Caroline A. Martin, Kylee R. Cogdill, Alesandra L. Pusey, Mary Gehring

## Abstract

Successful seed development is essential for flowering plant reproduction and requires the coordination of three genetically distinct tissues: the embryo and endosperm, which are the products of fertilization, and the maternal seed coat. Our understanding of the transcriptional programs underlying tissue-specific functions and inter-tissue coordination in seeds remains incomplete. To address this, we performed single nucleus RNA-sequencing on *Arabidopsis thaliana* seeds at 3, 5, and 7 days after pollination. We characterize all major seed cell or nuclei types, further refine transcriptional states in the endosperm, and map signatures of selection on cell type-specific genes. Among other findings, our analyses reveal the compartmentalization of genes involved in brassinosteroid-responsive transcription factor activation, abundant endosperm expression of genes that encode short, secreted peptides (SSPs), and expression enrichment of rapidly evolving genes in endosperm and seed coat subtypes, illuminating the cell type and species specificity of seed genes.

## Introduction

The three genetically distinct tissues of the seed are developmentally coordinated to ensure propagation of the next generation^1^. This process begins when the egg cell and central cell of the female gametophyte are fertilized by sperm to generate the embryo and endosperm, respectively. After fertilization the growth, development, and differentiation of the embryo and the more rapidly-developing endosperm are accompanied by growth and development of the ovule integuments, which become the seed coat^2,3,4,5^. The close synchronization of developmental transitions in the seed suggests widespread signaling between tissues, even though inter-tissue signals must cross cell walls and membranes as the embryo, endosperm, and seed coat are isolated symplastic fields^6,7^. Mechanisms underlying coordination within and between tissues are beginning to be elucidated^8^. For example, seed coat development requires endosperm-derived auxin, and embryo morphogenesis relies on auxin from the early seed coat^9,10,11^. The complete mechanisms for these interactions are unclear, and they are only two components of an extensive molecular dialogue.

The embryo-nourishing endosperm is a dynamic tissue that has been implicated in several axes of inter-tissue signaling^12,13,14^. It begins as a coenocyte, then undergoes cellularization before embryo invasion and consumption during the final stages of seed development^15^. Arabidopsis endosperm was previously transcriptionally characterized throughout seed development by laser-capture microdissection followed by microarray analysis, and was transcriptionally characterized at the single-nucleus level at 4 days after pollination (DAP)^16,17^. Endosperm nuclei that are at key tissue interfaces show the highest transcriptional distinction, namely the embryo-proximal micropylar endosperm (MCE) and the chalazal endosperm (CZE), which sits at the maternal-offspring interface and acts as a gateway for maternal resources into the embryo sac^16,17,18^. How endosperm compartments locally coordinate processes at tissue interfaces is incompletely understood. However, some short, secreted peptides (SSPs) from the micropylar endosperm are well-characterized inter-tissue signals. For example, the embryo-derived TWS1 SSP is processed in the endosperm by the ALE1 subtilase, and the mature peptide directs cuticle development in the embryo^19,20^. Given the symplastic isolation of seed compartments, SSP signaling could be a frequent conduit for inter-tissue signals.

Throughout seed development, maternal resources are deposited into, and distributed from, the chalazal seed coat (CZSC), a specialized region of the seed coat at the terminus of the vasculature^21,5^. Transporter- and channel-dense cells import nutrients and hormones and unload them into integument symplastic domains^22^. Recent evidence suggests that maternally synthesized auxin and abscisic acid are imported from the funiculus and control seed size and dormancy, respectively^23,24^. The CZSC is a morphologically complex region and the degree of functional specialization among CZSC cell types is unknown.

We present the first timepoint-resolved snRNA-seq atlas of early Arabidopsis seed development. Among other findings, our analyses revealed transcriptionally defined CZSC subtypes with complementary functions, and concentration of brassinosteroid biosynthesis in a micropylar seed coat subtype. In the endosperm, we report genes that underlie transcriptional polarity within the CZE cyst, many of them SSPs, which is consistent with a general enrichment of SSP expression in the CZE and MCE. Finally, we show that that the PEN and CZE express the majority of seed-expressed genes that appear to be under positive selection. Our atlas is available for exploration at https://seedatlas.wi.mit.edu/.

## Results

### A transcriptional atlas of early Arabidopsis seed development

To capture the most dynamic period of endosperm development, we isolated and sequenced RNA from individual nuclei from 3, 5, and 7 DAP Col-0 seeds using the 10x Genomics platform (Fig. 1A, Methods). At 3 DAP, the embryo is at the globular stage and the endosperm is coenocytic. At 5 DAP, the endosperm begins to cellularize at the micropylar pole, and at 7 DAP cellularization is complete and the embryo expands rapidly (Fig. 1A)^15^. For each time point we collected two biological replicates, which showed high transcriptional correlation within timepoints (Extended Data 1). After raw snRNA-seq data filtering and correction (Methods) we identified an optimal clustering resolution based on both the number of clusters and cluster neighborhood purity (Supplementary Fig. 1). We assigned clusters to cell types by analyzing the expression of known published marker genes, differential expression (DE) and GO term enrichment analysis, and by hybridization chain reaction RNA-fluorescent *in situ* hybridization (HCR RNA-FISH) of cluster marker genes (Supplementary Tables 1-4, Methods). We further subclustered embryo and endosperm data to identify previously-characterized cell types in the embryo and reveal putative novel nuclei types in the endosperm (Extended Data 2 and Supplementary Fig. 2-4). At the highest level of resolution, which we refer to as level 3 annotation (L3), we identified 34, 33, and 25 clusters at 3, 5, and 7 DAP, respectively (Fig. 1B-D, Extended Data 3, Supplementary Table 3). The L3 clusters were then assigned level 2 (L2) and level 1 (L1) annotations, which were harmonized across timepoints (Fig. 1B, C). Once each timepoint was annotated separately, the datasets were integrated into a final atlas dataset (Fig. 1B-D). In total, our atlas contains 54,210 profiles (24,024 at 3 DAP, 16,039 at 5 DAP, 14,147 at 7 DAP) and is approximately 10.5% embryo, 23.4% endosperm, 64.3% seed coat, and 1.8% unfertilized ovule and funiculus (Fig. 1E).

**Figure 1:**
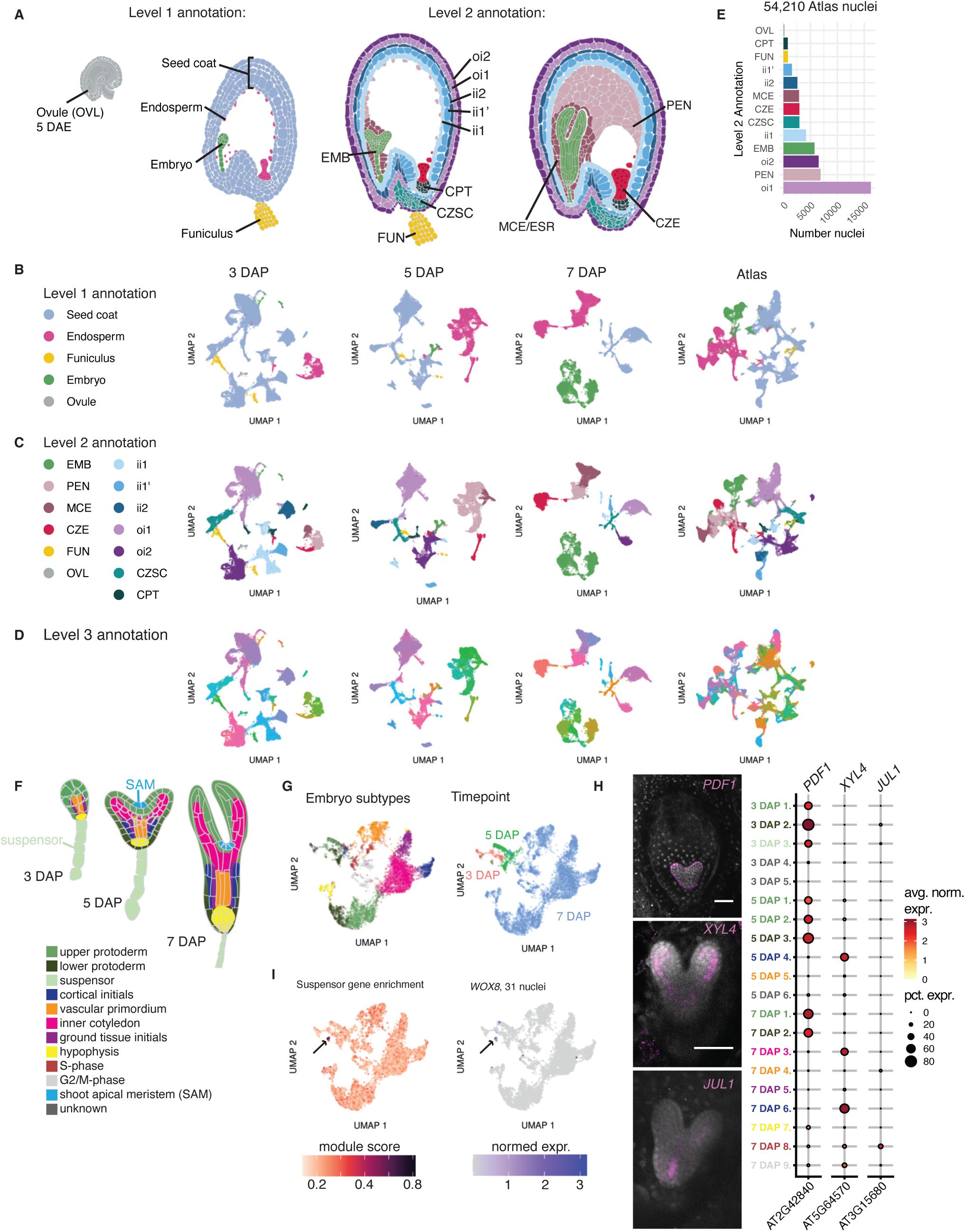
A transcriptional atlas of early Arabidopsis seed development. **A**, Seed developmental stages profiled in this study and Level 1 (L1) and Level 2 (L2) annotations. Outer integument 2 (oi2), outer integument 1 (oi1), inner integument 2 (ii2), inner integument 1’ (ii1’), inner integument 1 (ii1), embryo (EMB), chalazal proliferating tissue (CPT), chalazal seed coat (CZSC), funiculus (FUN), peripheral endosperm (PEN), micropylar endosperm (MCE), embryo surrounding region (ESR), chalazal endosperm (CZE), ovule (OVL). **B-D**, Level 1-3 annotations across all datasets. See Supplementary Table 3 for complete descriptions of all L3 annotations. **E**, Cell type proportions for L2 annotations in the full atlas dataset. **F**, Morphological stages of the embryo. **G**, Left: merged 3-7 DAP embryo datasets colored by annotations from F and right: timepoint. **H**, Left: HCR validation of the protoderm (*PDF1+*), inner cotyledon (*XYL4+*), and vascular primordium (*JUL1+*) in the embryo. Scale bar = 50 µm. *PDF1* and *JUL1* images are representative of signal observed in at least six seeds in two independent experiments, and the image for *XYL4* represents signal observed in four seeds in one experiment. Right: Expression of genes corresponding to HCR probes in the integrated embryo dataset, split by L3 annotation. **I**, Detection of the suspensor, a rare nucleus type in the merged embryo dataset. Left: module score analysis for 29 suspensor marker genes curated in Kao et al. 2021. Right: *WOX8* is a suspensor marker gene enriched in the suspensor cluster and detected in 31 nuclei in the entire dataset. Black arrows point to the putative suspensor population.

To assess atlas completeness, we determined whether we had captured rare embryonic cell types (Fig. 1F). The majority of embryo nuclei were isolated from seeds at 7 DAP (5,106), with 178 nuclei from 3 DAP and 399 nuclei from 5 DAP (Fig. 1G, Supplementary Table 3). The shoot apical meristem (SAM) and suspensor represent rare embryonic cell types, which respectively specifically express *CUC1* and *WOX8* (Fig. 1F)^25,26^. A sub-clustering analysis revealed 5 clusters in the 3 DAP embryo and 6 in the 5 DAP embryo (Extended Data 2, Methods). Using the embryo subtype markers curated in Kao et al. 2021, we annotated embryo subclusters with known subtypes, if possible^27^. We identified upper and lower protoderm populations at each timepoint and a 3 DAP-specific *WOX8+* suspensor population, among other subclusters (Fig. 1I and Extended Data 2). 7 DAP embryo subtypes were resolved at the *de novo* clustering resolution and clusters corresponding to the inner cotyledon, vascular primordium, ground tissue initials, cortical initials, and hypophysis were identified (Fig. 1G)^27–32^. Correspondence between clusters and cell types was validated by HCR RNA-FISH (Fig. 1H, Supplementary Fig. 5, Supplementary Table 1-2). *CUC1* was detected in 12 protodermal nuclei at 5 and 7 DAP, and these nuclei are enriched with SAM-specific gene expression (Extended Data 2). However, other embryo clusters showed similar levels of SAM-specific gene expression, and *WUS*, a known marker for a subpopulation of the SAM expressed early in development, is not detected in any embryo nuclei (Extended Data 2)^28,33^. This indicates that ∼54k nuclei from whole seeds enable the detection of some rare cell populations in the seed, such as the suspensor, but characterized SAM cell types appear to be absent. Overall, the clusters we defined include representatives of most previously anatomically and morphologically defined seed cell types (Fig. 1, Extended Data 3, Supplementary Table 3).

### A high-resolution census of the developing seed coat

The seed coat is the predominant seed tissue type at early stages (Fig. 1B, E), comprised of the five-layered testa, which in past studies has been referred to as the “general seed coat” (GSC), and the chalazal seed coat (CZSC), or the cell types near maternal vascular terminals (Fig. 2A)^17,34^. Each layer of the GSC is one cell thick, which has made comprehensive transcriptional profiling of these cell types difficult, although many layer-specific genes have been identified^35–40^. Using published markers, we identified all five layers of the seed coat in our L2 annotations across all timepoints, except for 7 DAP, for which there is a single ii1’/ii2 (inner integument 1’/inner integument 2) cluster (Extended Data 4, Supplementary Table 1, Supplementary Table 3).

**Figure 2:**
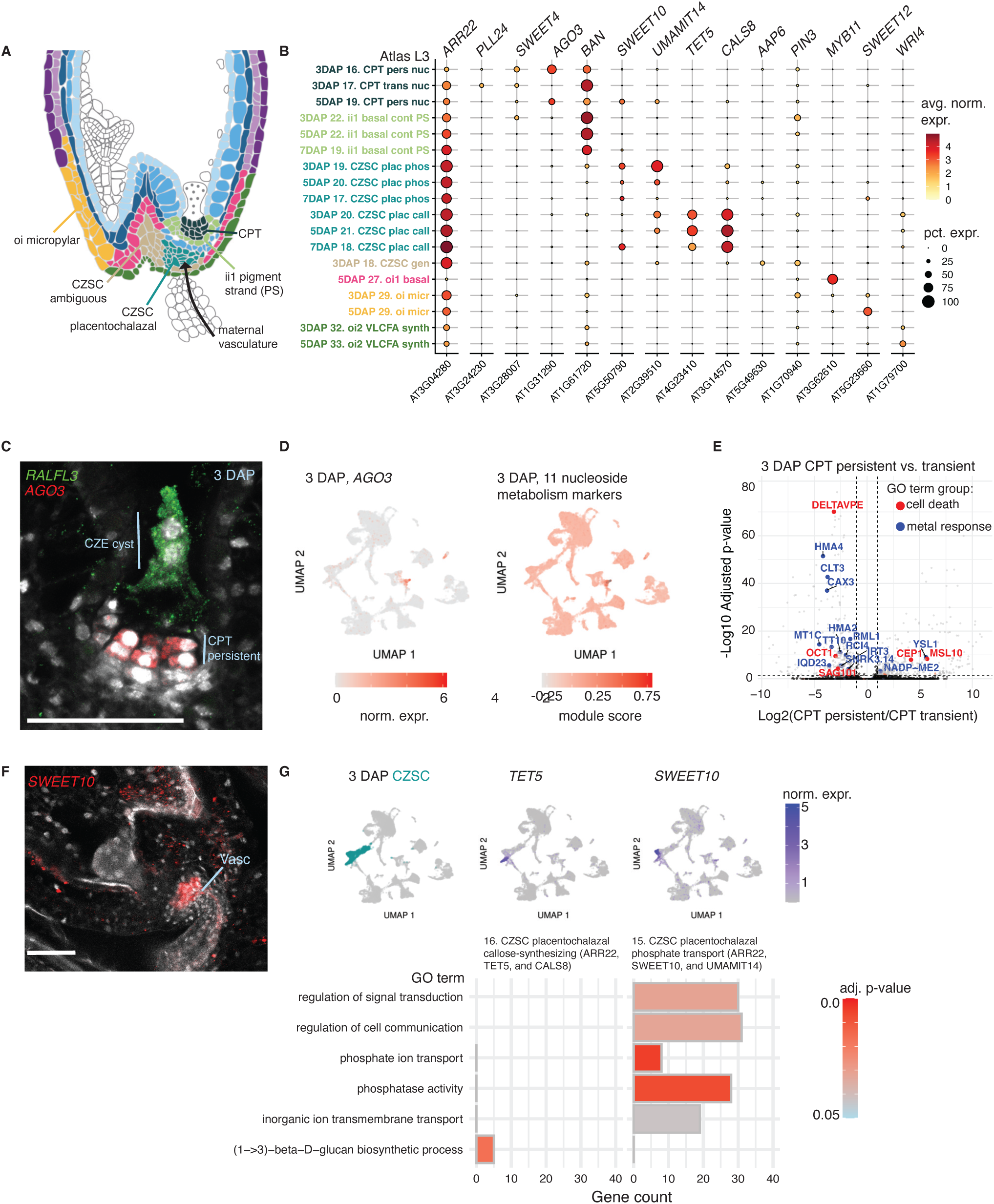
Specialized cell types of the lower seed coat. **A**, A model for the distribution of lower seed coat nuclei types identified in this study at 5 DAP. Xylem and phloem terminate in the CZSC. **B**, Expression patterns of published markers (Supplementary Table 1) and HCR-validated markers (*SWEET10, AGO3*) across all L3 annotations for the lower seed coat. See Supplementary Table 3 to match abbreviated L3 names to their full descriptions. **C**, *AGO3* labels a subpopulation of the CPT at 3 DAP by HCR, while *RALFL3* labels the CZE. Scale bar = 50 µm. The *AGO3* and *RALFL3* image is representative of signal observed in at least three independent experiments for each probe. **D**, *AGO3* is highly specific to CPT (left), and eleven nucleoside metabolism markers follow the same expression pattern (right). **E**, Differential expression between the 3 DAP CPT persistent and transient nuclei, adjusted p-values calculated through a two-sided Wilcoxon Rank Sum test with Bonferroni correction. **F**, *SWEET10* is localized to the placentochalazal region by HCR, and appears to be surrounding maternal vascular (Vasc) terminals. Scale bar = 50 µm. **G**, Top: *TET5* and *SWEET10* are the best markers for the L3 subtypes of the 3 DAP placentochalazal CZSC. Bottom: a subset of GO terms exclusively associated with DE genes (log2FC > 1, adjusted p-value ≤ 0.05, based on a hypogeometric test with Benjamini-Hochberg adjustment) for each placentochalazal cluster.

Whereas the upper seed coat contains the five principal layers, the lower seed coat is a densely heterogeneous region where maternal vasculature, the CZSC, and layers of the general seed coat meet (Fig. 2A). It serves roles in hormonal signaling and nutrient uptake in the developing seed^23,41–43^. To identify lower seed coat subtypes, we used the published marker *ARR22,* which is enriched in the CZSC but is also detected in adjacent lower seed coat tissues^44^. We identified 18 *ARR22*+ clusters across all timepoints (Fig. 2B), including subtypes of the GSC, CZSC, and the chalazal proliferating tissue (CPT), a small population derived from the nucellus and that lies beneath the chalazal endosperm cyst^45^. After fertilization, the CPT is partially degraded, characterized by “persistent” and “transient” CPT subtypes, and then fully degraded between 5-7 DAP^46,47^. We used the markers *SWEET4* and *PLL24*^48^ to identify the persistent and transient populations, respectively. *AGO3* promoter activity has been reported in the chalazal integument of seeds, and we defined its expression in the putative persistent CPT by HCR RNA-FISH (Fig. 2B-C)^49^. A GO term analysis using the top persistent CPT DE genes (log_2_FC > 1, adj. pval <0.05) indicated that it is a hotspot for nucleoside catabolism in the seed, consistent with ongoing programmed cell death (PCD) in this region (Extended Data 4)^46^. Eleven nucleoside catabolism genes showed high specificity for the persistent CPT (Fig. 2D, Extended Data 4). DE analysis between the 3 DAP persistent and transient CPT clusters implicated genes associated with cell death and metal response (Fig. 2E). Notably, *DELTAVPE*, a contributor to cell death in the inner integument^39^, is DE in the transient CPT, while *MSL10*, an ion channel that positively regulates programmed cell death in a mechanically sensitive manner^50^, is DE in the persistent population, suggesting differing cell death triggers in the transient and persistent CPT. This represents the first transcriptomic characterization of the CPT in Arabidopsis.

### Cell types in the placentochalazal region of the chalazal seed coat have complementary functions

The CZSC is appreciated for being the primary site of active nutrient transfer into the seed from maternal vascular terminals, facilitated by SWEET, UMAMIT, and PHO1 transporters ^41,42,51,52^. The cells closest to the maternal vasculature comprise the placentochalazal region and specifically express the gene encoding the UMAMIT14 amino acid transporter^51^ (Fig. 2A, B). Based on *UMAMIT14* and high *ARR22* expression, we identified six putative CZSC clusters across all timepoints, which were grouped into three subtypes based on the expression of *SWEET10*, *TET5*, and *AAP6* (Fig. 2B, F, Supplementary Table 1). Of these, we found two putative placentochalazal subtypes at 3 and 5 DAP CZSC, which were labeled specifically by *SWEET10* and *TET5* and shared *UMAMIT14* expression (Fig. 2B, G). We identified DE genes in these subtypes compared to all other clusters at 3 DAP (average log_2_FC > 1, adj. p-value >0.05) (Fig. 2G, Extended Data 5). *TET5*+ cluster DE genes are associated with the GO term “(1->3)-beta-D-glucan (callose) metabolic process”. Callose deposits are known to regulate plasmodesmata activity and serve an insulating role between cell types of the developing ovule^53^. A callose-rich “phloem end” has recently been described in the CZSC^54^. Module score analysis (Methods) using all genes associated with the GO term “callose biosynthesis” revealed high, significant enrichment in *TET5*+ clusters across all timepoints (Extended Data 5). Several putative callose biosynthesis genes are specifically expressed in this population, the strongest being *CALS8* (Fig. 2B, Extended Data 5).

In contrast, top exclusive GO terms for the *SWEET10*+ cluster DE genes include “phosphate ion transport”, and “phosphatase activity”, supported by the differential expression of *PHO1;H1*; and *TPPE, TPPB,* and *TPPG*, respectively (Fig. 2G, Extended Data 5). *PHO1;H1* is a phosphate exporter with demonstrated activity in the *Arabidopsis* CZSC^42^. Additionally, trehalose phosphatase expression suggests that these cells participate in trehalose-6-phosphate signaling, which directs sugar utilization^55–58^. The invertase *cwINV4* sugar transporter gene is also specifically expressed in this cluster (Extended Data 5). These results indicate that the *SWEET10+* cluster is likely the primary site of nutrient transfer in the placentochalazal CZSC, and that the *TET5*+ and *SWEET10*+ populations have complementary functions influencing the permeability of maternal vascular terminals and surrounding cells.

### Brassinosteroid biosynthesis, homeostasis, and response genes show concentrated expression in the micropylar region of the seed

Brassinosteroids (BR) are hormones that act widely in plant physiology and are known to direct organ formation and cell expansion in reproductive tissues^59–65^. In the seed they promote endosperm proliferation by reducing the physical resistance of the seed coat through cell wall weakening^65^. Although maternal seed coat-derived BR are hypothesized to interact directly with the endosperm, a recent study indicates that BR controls endosperm proliferation through cell autonomous effects in the seed coat^65^. To identify sites of BR biosynthesis in the developing seed, we performed a module score analysis for the GO term gene sets “BR biosynthesis” and “BR homeostasis.” Across all timepoints, the ii2 (inner integument 2) seed coat layer showed the greatest enrichment for BR biosynthesis, followed by the oi1 (outer integument 1) cluster (Fig. 3A,B). Additionally, partitioning by timepoint and L3 annotation revealed that a putative micropylar oi subtype (Fig. 3A-C) drives the L2 oi1 enrichment for BR biosynthesis gene expression and exhibits the highest enrichment for genes associated with BR homeostasis atlas-wide. We annotated an oi micropylar cluster based on the specific expression of *SWEET12* (Fig. 2B), which has previously been localized to the micropylar end of the seed coat, and because the cluster shows high transcriptional similarity to oi clusters, with a slight bias toward oi1 (Supplementary Fig. 6)^52^. An inspection of the genes underlying the micropylar oi enrichment scores revealed that the enzymes that catalyze the last two steps of BR biosynthesis, *BR6OX1* and *BR6OX2*, are upregulated in this subtype (Fig. 3D). We propose that the micropylar oi is a key site for brassinosteroid production at early to intermediate stages of seed development.

**Figure 3:**
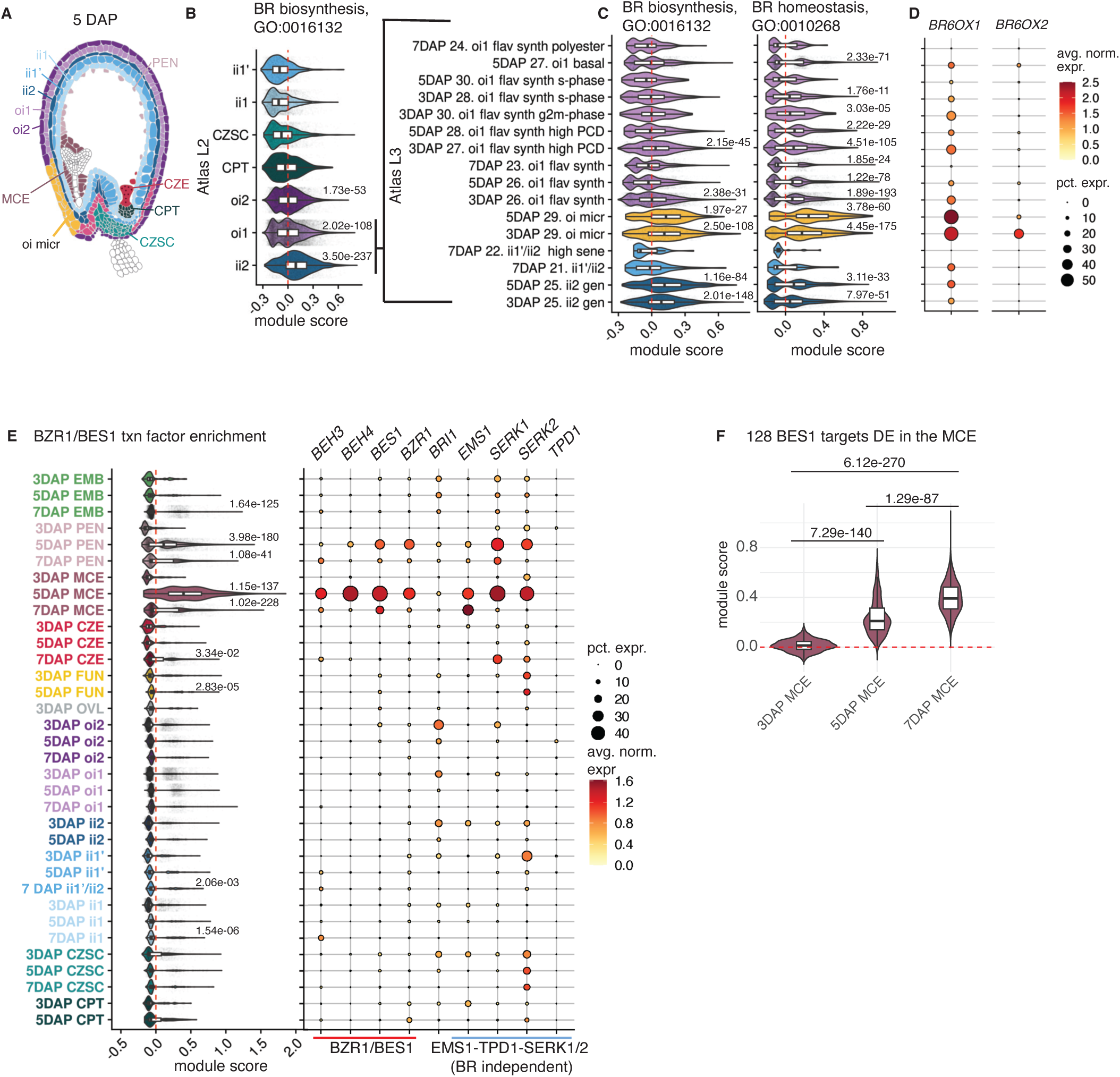
Genes that underlie BZR/BES1 transcription factor activity are expressed in the micropylar region of the seed. **A**, A subset of 5 DAP L2 annotations with the L3 oi1 micropylar region highlighted. **B**, Module score analysis for all genes associated with brassinosteroid (BR) biosynthesis (GO: 0016132) in the L2 seed coat across all timepoints. **C**, Module score analysis for BR biosynthesis and homeostasis (GO:0010268) in a subset of L3 seed coat clusters. See Supplementary Table 3 to match abbreviated L3 names to their full descriptions. **D**, Gene expression pattern for *BR6OX1* and *BR6OX2*. **E**, Module score analysis for all six BZR/BES1 transcription factors detected in the atlas (*BES1, BEH1, BEH2, BEH3, BEH4, BZR1*). *BES1, BEH3, BZR1*, and *BEH4* show the strongest expression and are enriched in 5 DAP MCE. Components of the BR-independent BES1 activation pathway (*EMS1, SERK1/2*) show overlapping expression with *BES1*, although *TPD1* shows low, nonspecific expression throughout seed development. **F**, Module score analysis for 128 BES1 targets (identified in O’Malley et al. 2016 and DE in the atlas MCE (log2FC > 1, adjusted p-value ≤ 0.05)), showing an increase in BES1 target genes through development. In B, C, and E, adjusted p-values are to the right above clusters with significantly high average positive module scores in a cluster-vs-all other nuclei comparison (sample size = 54,210, treating individual nuclei as biological replicates). In F, adjusted p-values were generated from pairwise comparisons (sample sizes: 3 DAP MCE = 524, 5 DAP MCE = 426, and 7 DAP MCE = 1929, treating individual nuclei as biological replicates). All p-values are derived from a two-sided Wilcoxon Rank-Sum test with Bonferroni correction. See Supplementary Table 7 for the module scores, p-values, and pairwise group sizes for all comparisons. For all boxplots, the center corresponds to the median, the upper and lower hinges correspond to the 25th and 75th percentiles, and the whiskers extend to the highest and lowest values that are within 1.5 times the interquartile range.

BZR1/BES1 transcription factors are activated by BR via the BRI receptor-like kinase, although BR-independent activation of *BES1* by the SSP TPD1 through EMS1-SERK1/2 signaling has also been observed^66^. Once activated, they promote the expression of genes involved in cell elongation, light-regulated development, and cell wall remodeling^66–71^. The activity of some BZR1/BES1 family members has been characterized in the endosperm: constitutively active *BES1* or *BZR1* causes reduced proliferation of endosperm nuclei, whereas quintuple mutants of BZR1/BES1 family members (*bzr1;bes1;beh1;beh3;beh4)* show no endosperm phenotype, suggesting redundant mechanisms for BZR1/BES1-family contributions to endosperm development^65^. To map BZR1/BES1 activity, we performed a module score analysis for the six BZR1/BES1 genes detected in the atlas, revealing high and specific expression in the 5 DAP MCE, driven by *BES1*, *BEH3*, *BZR1,* and *BEH4* (Fig. 3E). The transmembrane BR receptor *BRI1* shows broad expression throughout the seed coat but is depleted in the 5 DAP MCE (Fig. 3E). To determine whether BZR1/BES1 may be activated in a BR-independent manner in the MCE, we mapped the expression of *EMS1*, *TPD1*, and *SERK1/2*. Although *EMS1* and *SERK1/2* show overlapping spatiotemporal expression patterns with MCE BZR1/BES1 transcription factors, *TPD1* is nearly undetectable atlas-wide (Fig. 3E).

To test whether the expression of *BES1* targets show increased expression after *BES1* upregulation in the MCE, we performed a module score analysis of all *BES1* targets that are DE in the MCE (log_2_FC > 1, adj. pval <0.05), which indicated significant upregulation from 3 to 7 DAP (Fig. 3F)^72^. The co-occurrence of BR biosynthesis and response genes in proximal micropylar cell types, the absence of *TPD1*, and the timing of BES1 target upregulation supports the hypothesis that BRs are transported from the micropylar region of outer integument 1 to the endosperm. However, it is plausible that *EMS1* and *SERK1/2* may be activated by an alternative TPD1-like SSP, such as TPD1-like 1 (TDL1), which is the seed SSP that shows the highest sequence similarity to TPD1 of all Arabidopsis SSPs and is expressed in the early embryo (Supplementary Fig. 7). In that case, BZR1/BES1 activation might also occur cell non-autonomously, but in a BR-independent manner.

### The micropylar-to-Embryo Surrounding Region endosperm shift is characterized by UMAMIT and nitrate transporter gene expression

Based on a correlation analysis of L2 endosperm clusters throughout development, a dramatic transcriptional shift occurs in the endosperm between 5-7 DAP. We observed that the 7 DAP endosperm is the most transcriptionally divergent endosperm cluster (Extended Data 6). Within 7 DAP endosperm clusters, the *GLIP6+* MCE (referred to as the “embryo surrounding region” (ESR) at this stage), is the most distinct, driven by the upregulation of *UMAMIT*s and nitrate transporters (Extended Data 6). The ESR is known to shape embryo viability by controlling the formation of two barriers that support successful germination: the embryonic cuticle and sheath, which respectively prevent embryo dehydration and endosperm adherence^19,73^. These processes are coordinated with programmed cell death through the transcription factor *ZOU/RGE1*^73–76^. At 7 DAP, we observed two clusters within the *ZOU+* population: one expressing *KRS,* a signaling peptide that directs embryo sheath formation, and another specifically expressing *NAC074* and enriched for *NAC087,* which directs programmed cell death in the endosperm (Extended Data 6)^73,76^. *ZOU* induces *KRS* expression^74^ and likely controls PCD in partially parallel pathways with NAC transcription factors^76^, and these genes appear to be distributed on a continuum of MCE nuclei states at 7 DAP. Other genes follow this pattern, such as the nitrate transporters, which are enriched in the *KRS+* cluster (Extended Data 6).

### The developmental basis for transcriptional polarity in the chalazal endosperm

The chalazal endosperm (CZE) is composed of an unusual population of nuclei that does not cellularize^77^. The CZE is important for maternal resource allocation in the developing seed due to its position at the maternal-filial interface, and it exhibits the strongest parent-of-origin allelic expression bias for imprinted genes of all endosperm subtypes^16,78^. In particular, paternally expressed genes (PEGs) are upregulated in the CZE (Supplementary Fig. 8)^16^. A corresponding enrichment of epigenetic and transcriptional regulator expression in the CZE has been described^16,79^. This atlas contains 2,941 CZE nuclei, enabling higher-powered differential expression analyses. We inspected expression patterns of 460 chromatin-associated genes that showed variable endosperm subtype specificity in our previous snRNA-seq study of 4 DAP endosperm and found that most of these genes are enriched in early endosperm subtypes. Furthermore, some of the epigenetic regulators previously described are depleted in the endosperm and enriched in seed coat layers (Supplementary Fig. 8). To identify additional epigenetic regulators that vary between seed cell and nuclei types, we inspected DE genes associated with chromatin (GO:0000785), transcriptional regulation of gene expression (GO:0010468), and epigenetic regulation of gene expression (GO:0040029) and found 736 additional endosperm-variable genes. This updated gene set generally follows the enrichment pattern exhibited by the gene list from our previous study (Supplementary Fig. 8, Supplementary Table 5)^16^.

A subset of CZE nuclei form the multi-nucleate cyst, a coenocytic sac isolated by the central vacuole, and another subset form the nodules, which lie above the cyst (Fig. 4A)^80^. Developmental time course and live imaging studies of Arabidopsis endosperm indicate that the cyst grows in part through fusion with proximal nuclei from the nodule^3,80,81^. The cyst and nodule were transcriptionally characterized in our previous snRNA-seq study of 4 DAP endosperm, which also described a “nodule-like” chalazal population that was not morphologically defined^16^. We first identified CZE nuclei populations using marker gene sets for the chalazal cyst, nodule, and nodule-like populations defined from 4 DAP, which include AT2G44240 (cyst), *CYCD4;2* (nodule), and *MEA* (nodule-like) (Fig. 4D)^16^. These markers were not detected at 7 DAP, so *NPF4.5* was used to define the 7 DAP CZE (Fig. 4D, Supplementary Fig. 5, Supplementary Table 1)^17^.

**Figure 4:**
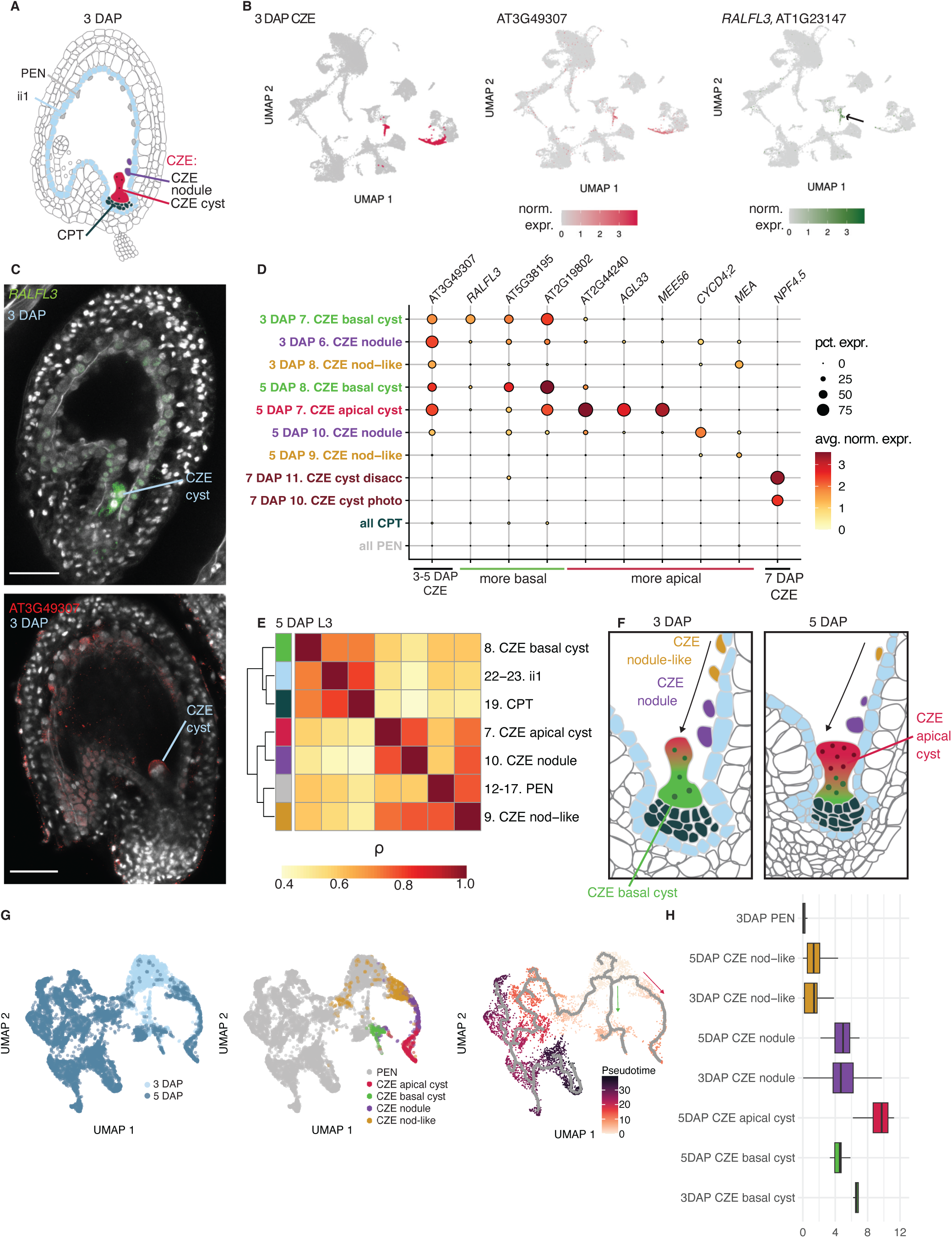
The developmental basis for transcriptional polarity in the chalazal endosperm. **A**, L2 annotations of chalazal-proximal tissues, including the CZE, CPT, PEN, and ii1. The chalazal nodule and cyst are morphologically characterized compartments of the CZE. **B**, Left: the CZE highlighted in the 3 DAP dataset. Middle, right: AT3G49307, a CZE marker, and *RALFL3* (arrow), a marker for a subtype of the CZE at 3 DAP. **C**, HCR validation of *RALFL3* (top) and AT3G49307 (bottom) expression in 3 DAP seeds. Scale bar = 50 µm. Images are representative of signal observed in at least four independent experiments for each probe. **D**, Expression patterns of CZE subtype markers from Picard et al. 2021 (*MEA, CYCD4;2*, AT2G44240) and this study in all L3 clusters for CZE, PEN, and CZE-proximal tissues at all timepoints. See Supplementary Table 3 to match abbreviated L3 names to their full descriptions. **E**, Clustered heatmap of the Spearman correlation coefficients for aggregated expression of the 5 DAP L3 clusters for CZE, PEN, and CZE-proximal maternal subtypes. **F**, A model for CZE cyst growth through fusion with proximal nuclei based on live imaging studies and our transcriptional characterization. **G**, Left: timepoint annotations for the integrated 3-5 DAP CZE and PEN dataset used for pseudotime analysis. Pseudotime is a proxy for progression along a developmental trajectory from the 3 DAP PEN. Middle: L3 annotations for CZE subtypes and L2 annotation for the PEN in the 3-5 DAP integrated datasets. Right: integrated 3-5 DAP datasets colored by 3 DAP PEN-anchored pseudotime, with the monocle3 principal graph indicating developmental branches from the 3 DAP PEN. Green arrow indicates the putative “basal” trajectory from the early PEN, while the red denotes the “apical” branch. **H**, Pseudotime distributions for CZE and 3 DAP PEN nuclei types. The nodule-like and nodule populations represent transitional states between the PEN and CZE. For all boxplots, the center corresponds to the median, the upper and lower hinges correspond to the 25th and 75th percentiles, and the whiskers extend to the highest and lowest values that are within 1.5 times the interquartile range.

Differential expression analysis of the 3 DAP L2 clusters revealed that the SSP gene *RAPID ALKALINIZATION FACTOR-LIKE 3* (*RALFL3*) was a highly specific marker for a subpopulation of CZE nuclei (Fig. 4B, Supplementary Fig. 2-3). *RALFL3* HCR RNA-FISH indicated transcript accumulation either throughout or at the base of the chalazal cyst (Fig. 2C, Fig. 4C, Extended Data 7). We compared the *RALFL3* HCR signal with that of AT3G49307, an SSP gene more broadly expressed in the CZE, and found the highest AT3G49307 signal in the apical region of the cyst and nodules throughout seed development (Fig. 4C, Extended Data. 7). We propose that, rather than being a structure of uniform gene expression, there is a transcriptional apical-basal axis within the chalazal cyst, with AT3G49307 and *RALFL3* labeling the apical and basal regions, respectively. At 3 DAP, the *RALFL3+* basal state predominates in the cyst, but at 5 DAP the apical/basal distinction is pronounced and two cyst states are detectable as subclusters (Fig. 4D). A correlation analysis of aggregated gene expression for cell types within and adjacent to the CZE showed high transcriptional correlation between the basal cyst and proximal maternal tissues, the ii1 and CPT (Fig. 4A, E).

Based on high *RALFL3* HCR signal in CZE cysts when ∼3 nuclei are visible and the transcriptional similarity of *RALFL3*+ basal cyst clusters to both CZE subtypes and proximal maternal tissues, we hypothesize that the *RALFL3*+ basal cyst nuclei represent the “founder” nuclei that migrate to the chalazal region at early timepoints, to which subsequent nuclei fuse to generate the mature chalazal cyst (Fig. 4F, Extended Data 8). To characterize the developmental landscape of the CZE, we performed trajectory inference and pseudotime analysis on the 3 and 5 DAP PEN and CZE, anchored in the 3 DAP PEN (Methods). Pseudotime values are a proxy for progression on a developmental trajectory from the 3 DAP PEN. This analysis revealed two branches, one connecting the 3 DAP PEN with the 3-5 DAP basal cyst clusters and another connecting the 3 DAP PEN with the rest of the 3-5 DAP CZE subtypes. This suggests that the basal cyst has an independent developmental trajectory from the rest of the CZE (Fig. 4G). Pseudotime analysis positioned the nodule-like population closest to the early PEN, and this population might represent the initial commitment to the CZE-bound state (Fig. 4H). However, the majority of the 3 and 5 DAP nodule- and nodule-like populations are both positioned on the non-basal cyst branch, suggesting that the transcriptional underpinnings of the PEN-to-CZE transition are distinct for basal cyst nuclei compared to the rest of the CZE (Fig. 4G, Extended Data 8). Pseudotime within nodule and nodule-like subtypes are similar across 3 and 5 DAP, suggesting that although nuclei migrate through these transition states on their way to the CZE cyst, the states themselves are stable (Fig. 4H).

To identify DE transcription factors that might promote the PEN-to-CZE transition, we performed graph autocorrelation analysis using the 3-5 DAP PEN and CZE dataset (Methods). Focusing on the non-basal cyst branch, we identified 56 transcription factors that significantly vary in pseudotime and are DE (log_2_FC > 1, adj. pval <0.05) in the nodule-like and nodule populations, such as *HDG8* and *GRF2* (Extended Data 8, Supplementary Table 6).

### Discrete families of short, secreted peptides are enriched in endosperm subtypes

We observed that many endosperm subtype marker genes are SSPs, such as *RALFL3* AT3G49307, and *KRS* (Fig. 4, Extended Data 6, Extended Data 7). SSPs are abundant in plant genomes and play roles in reproduction, development, and innate immunity^82–87^. They are characterized by an N-terminal secretory signal sequence and are less than 250 amino acids long (Fig. 5A)^88–90^. Arabidopsis seeds devote more of their transcriptomes to genes that encode SSPs than any other tissue, but little is known about the functions of seed-specific SSPs (Extended Data 9)^91^. This is in part due to their absence from existing stage-resolved transcriptional atlases of seed development. For example, a previous atlas is based on ATH1 microarray data, which had probes for only ∼50% of SSP genes with conserved SSP motifs (Extended Data 9)^17,88^.

**Figure 5:**
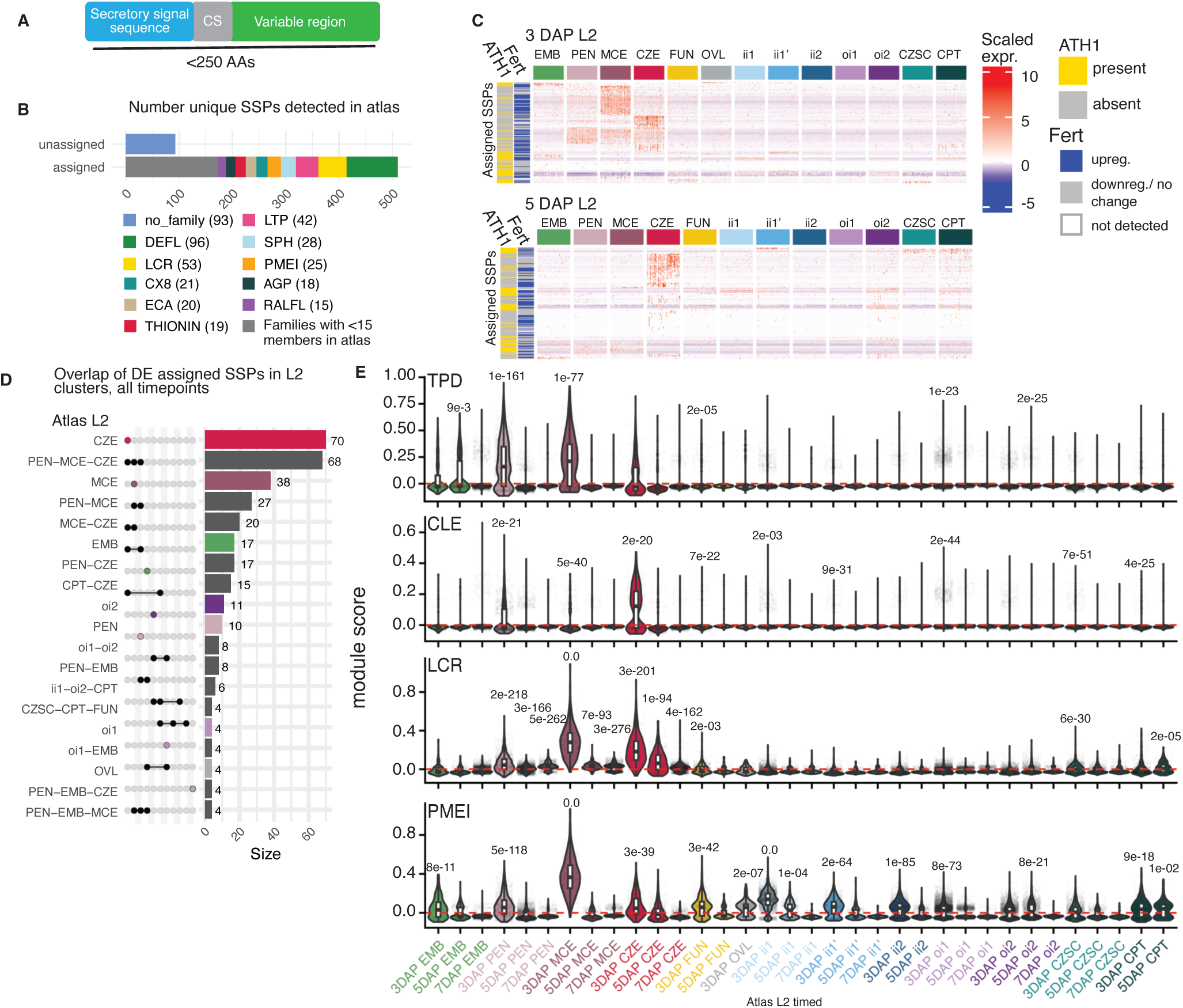
Short, secreted peptides are enriched in subtypes of the endosperm. **A**, Conventional structure of short, secreted peptides (SSPs), and the criteria used for SSP detection. Cleavage site (CS). **B**, Number of SSPs detected in the atlas assigned to characterized families based on a motif analysis (Methods), and the number of each peptide family represented among the assigned SSPs. The grey area in the “assigned” bar consists of all peptides in families with less than 15 SSPs detected in the atlas. **C**, Clustered heatmaps of scaled gene expression for variable SSPs detected in the atlas. The row annotations “ATH1” and “Fert” indicates presence or absence from the ATH1 microarray, and whether the SSP was upregulated after fertilization (adjusted p-value ≤ 0.05, log2FC > 0.5, limma t-test with Benjamini-Hochberg correction) in the bulk expression data from Figueiredo et al. 2016. **D**, Upset plot indicating the proportion of unique SSPs DE (adjusted p-value < 0.05, log2FC >1, two-sided Wilcoxon Rank-Sum test with Bonferroni correction) at any time in development for the L2 clusters. **E**, The results of module score analysis for four SSP families containing members with signaling (TPD, CLE, and LCRs) as well as inhibitory roles (PMEI). Adjusted p-values are centered above clusters with significantly high average positive module scores in a cluster-vs-all other nuclei comparison (Wilcoxon Rank-Sum test with Bonferroni correc-tion, sample size = 54,210, treating individual nuclei as biological replicates). See Supplementary Table 7 for the module scores, p-values, and pairwise group sizes for all comparisons. For all boxplots, the center corresponds to the median, the upper and lower hinges correspond to the 25th and 75th percentiles, and the whiskers extend to the highest and lowest values that are within 1.5 times the interquartile range.

To characterize the cell- and nuclei-type specificity of seed SSPs and SSP families, we performed DE analysis using genes previously annotated with SSP motifs^88^. This analysis revealed that SSPs containing defensin-like (DEFL), low molecular weight cysteine-rich (LCR), and lipid transfer (LTP) motifs are the three predominant SSP families in the atlas (Fig. 5B, Supplementary Table 10, Methods)^88^. Although most have not been functionally characterized, members of these families have been implicated in defense and signaling; some play roles in pollen-pistil interactions^85^. We found that the 3 DAP MCE, 3 DAP CZE, and 5 DAP CZE are hubs of SSP expression, both by expression level and number of unique SSPs expressed (Fig. 5C, D). The SSPs enriched in the MCE and CZE have motifs found in families with characterized roles in cell-cell signaling (TPD, CLE, LCR), as well as inhibitory roles (PMEI)^92–95^. Furthermore, many exhibit upregulation after fertilization (Fig. 5C, Methods). SSP enrichment and diversity in the CZE and MCE are compelling from a signaling point of view because these regions are important interfaces: the CZE is a gateway for maternal resources into the seed and the MCE is the most embryo-proximal seed tissue (Fig. 1A).

### Rapidly evolving single-copy orthologs are compartmentalized in the endosperm

Previous studies have suggested that seed genes show higher rates of rapid evolution than other tissue-specific genes, with those specifically expressed at maternal-offspring interfaces showing the highest evolutionary rates^96^. One explanation for this is that genes involved in maternal resource allocation are expected to rapidly evolve due to intrafamilial conflict. A previous study of seed tissue-specific gene evolution analyzed signatures of selection for gene sets but did not resolve individual rapidly evolving genes. To identify individual rapidly evolving seed genes, their protein domains under selection, and their expression patterns in seed cell and nuclei types, we used codon-substitution site models of positive, negative, and neutral selection implemented in the codeml program in the PAML package to calculate the likelihood of positive selection for all single-copy orthologs (SCOs) shared by Arabidopsis, *Arabidopsis lyrata*, *Arabidopsis arenosa*, and *Capsella grandiflora* (Methods)^97^. This analysis produces a likelihood ratio test statistic (LRT) for each SCO, which indicates the goodness-of-fit of its phylogeny to a positive (M2a) or nearly neutral (M1a) model of selection. We generated LRTs for 7,187 SCOs and found that 141 seed genes have statistically high M2a/M1a LRTs (“M2a/M1a-sig”), 103 of which are DE among seed clusters.

An inspection of the 103 DE M2a/M1a-sig SCOs revealed that endosperm subtypes differentially express the highest number of M2a/M1a-sig SCOs (Fig. 6A, Supplementary Table 8). However, a module score analysis of M2a/M1a-sig SCOs showed that the ii1’ and ii2 seed coat layers have the highest expression enrichment, indicating that these subtypes highly express a small number of M2a/M1a-sig SCOs (Fig. 6A, Extended Data 10). Indeed, 17 genes DE in the ii1’ and ii2 seed coat layers are M2a/M1a-sig SCOs, but the M2a/M1a-sig expression enrichment appears to be largely driven by *DELTAVPE,* which has a rapidly evolving site in a C-terminal legumain prodomain (Pfam ID: PF20985) (Extended Data 10). Intersecting all 359 high-confidence rapidly evolving sites (p(dN/dS) > 0.95, Bayes Empirical Bayes) within M2a/M1a-sig SCOs with predicted protein domain coordinates revealed a functionally heterogeneous protein domain list, including 50 and 25 unique PANTHER and Pfam domains, respectively (Supplementary Table 8). A plurality of rapidly evolving sites were detected in extracellular domains or signal peptides of secreted proteins (147/359) (Fig. 6A). Intrinsically disordered regions (IDRs) were the second most prevalent selected domain (63/359) (Extended Data 10). There were no statistically significant shared GO terms among all DE M2a/M1a-sig SCOs, but genes implicated in protein degradation, secretion, and transcriptional regulation recurred in the M2a/M1a-sig SCOs list (Supplementary Table 8). For example, *AFA1* and *HON1* encode an F-box protein (a putative E3 ubiquitin ligase adaptor) and histone H1.1, which contains a winged-helix DNA-binding domain, respectively, and both are differentially expressed in the endosperm and CPT (Fig. 6B-C). *XYN4*, the most endosperm-specific M2a/M1a-sig SCO, unusually does not have secreted regions or IDRs with selected sites (Fig. 6B-C). Taken together, this analysis supports the finding that endosperm subtypes are enriched for rapidly evolving genes and identifies sites in secreted extracellular domains and IDRs that are the targets of positive selection in seeds.

**Figure 6:**
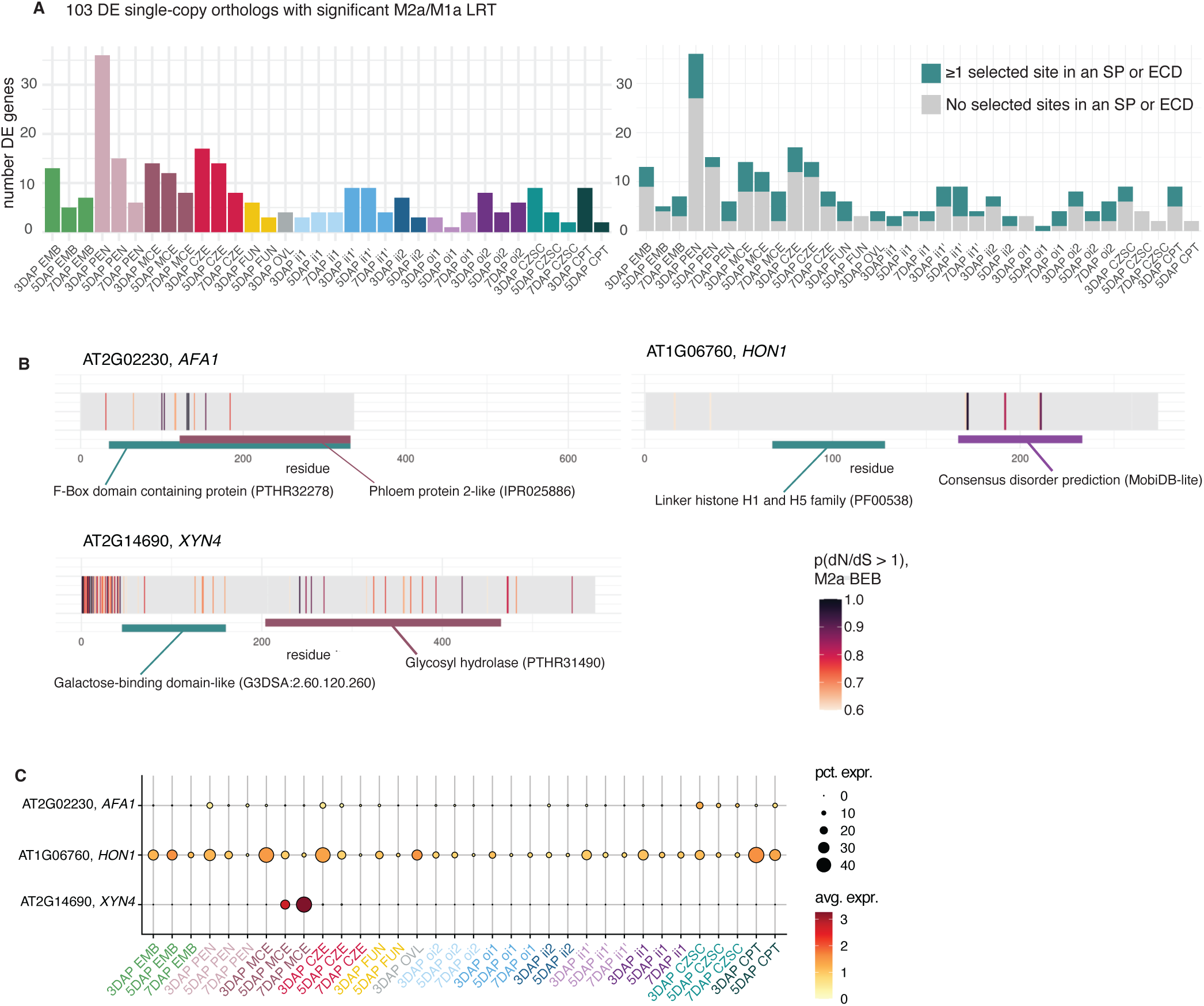
Rapidly evolving genes differentially expressed in seed tissues. **A**, Left: the number of differentially expressed genes (adjusted p-value ≤ 0.05, log2FC > 1, two-sided Wilcoxon Rank-Sum test with Bonferroni correction) with a significant M2a/M1a LRTs for timed atlas L2 clusters. Right: the same genes as left, but labeled if at least one selected site falls in an extracellular domain (ECD) or signal peptide (SP), as predicted by Phobius. **B**, The residues likely under positive selection in three select genes differentially expressed in the endosperm (adjusted p-value ≤ 0.05, log2FC > 1, two-sided Wilcoxon Rank-Sum test with Bonferroni correction). Coding sequences were translated and individual residues colored by the Bayes Emprical Bayes (BEB) posterior probability of having a dN/dS > 1 under the M2a model. Informative protein domains near or containing selected sites are highlighted. Pfam, PANTHER, and InterPro identifiers from InterProScan in parentheses. Consensus disorder regions are predicted by MobiDB-lite via InterProScan. Domain coordinates were taken from InterProScan for the longest isoform. **C**, Expression patterns for the genes shown in B in timed L2 clusters.

## Discussion

We present a comprehensive atlas of early seed development that illuminates aspects of cell-cell signaling, functional compartmentalization, and gene diversification in transcriptionally distinct cell or nuclei types. This adds to a growing compendium of single cell or nuclei transcriptional atlases for various stages of development in Arabidopsis and other plant species^98–103^. We revealed additional insights into nutrient transport within the seed as well as sites of callose and brassinosteroid biosynthesis. Our high coverage of endosperm nuclei allowed for the identification of a rare nuclei population in the CZE, which may clarify the origins of the CZE cyst. Furthermore, this atlas will enable the identification of promoters with high spatial and temporal specificity and will serve as a community resource for seed research.

Our study has also revealed additional complexity within the chalazal endosperm. We propose that *RALFL3*+ CZE nuclei may be the “founder” CZE population to which subsequent nuclei fuse to create the cyst. A time-lapse live-cell imaging study of Arabidopsis coenocytic endosperm development described two nuclei that migrate to the CZE, divide once, and persist at the chalazal pole^104^, and we hypothesize that these are the *RALFL3+* founders. We further posit that nodule nuclei fusion to the early CZE generates an apical/basal gradient of gene expression in the CZE at early timepoints. Notably, cytological features following this polarity have been observed in the CZE cyst: mitochondria and thylakoid stacks are enriched in the basal and apical regions, respectively^105^. The transcriptional similarity of the basal cyst with the CPT and ii1 suggests congruence or coordination with maternal tissue transcriptomes. Interestingly, “tentacle-like processes” embedded in maternal tissues have been described at the base of the Arabidopsis cyst^105^ and in other species the chalazal endosperm takes on haustorial properties, which could facilitate such coordination^16^. How the *RALFL3*+ nuclei are established at the chalazal pole, and the extent to which their DE SSPs contribute to this process, remains to be studied.

Prior to this work, we had only partial understanding of the extent of SSP expression and diversity in seed cell types. Although the MCE is a well-appreciated source of signaling SSPs, most of the MCE/CZE-specific SSPs were not assayed in existing transcriptional seed atlases, which may have led to an underestimation of the contribution of SSPs to seed development. Some of the MCE/CZE SSPs that are also expressed in the ovule could function in fertilization, but many are more highly expressed after fertilization, suggesting alternative functions. Many of the seed SSPs are annotated as defensins. This class of cysteine-rich peptides is typically thought of as acting as anti-microbial peptides in seeds, particularly against fungi^106^. However, few seed-expressed defensins have been specifically evaluated for anti-microbial activity. It is possible that defensins, and other seed SSPs, act as ligands that are perceived by receptor-like kinases or receptor-like proteins to activate a variety of signaling pathways. The atlas will further allow the evaluation of the expression of potential receptors in various seed compartments. The enrichment for SSP expression at the embryo-endosperm and maternal-offspring interfaces is consistent with cell non-autonomous functions, although this remains to be demonstrated for individual peptides.

Our study supports the finding that the endosperm is enriched for rapidly evolving genes and highlights signatures of rapid evolution in seed coat layers. The majority of positively-selected sites fall in secreted extracellular domains and IDRs. The structural flexibility of IDRs allows them to engage in diverse interactions, and as a result, they are often implicated in cell signaling and gene expression regulation. Thus, our findings suggest that protein-protein interactions involved in signal transduction within and outside of the cell might be sites of rapid evolution in the seed. However, it is unclear to what extent the rapid evolution observed in these IDRs is due to their putative functions or lack of structural constraint^107^. A limitation of our approach is that we restricted our analysis to SCOs to prevent evolutionary analyses on false orthologs, thus omitting genes that are members of expanded families. We provide the list of M2a/M1a-sig SCOs and the coordinates of their rapidly evolving protein domains to guide future hypotheses about the functions of rapidly evolving seed genes. Functional studies are needed to discern whether these genes underlie the roles and diverse morphologies of Brassica seeds.

Taken together, we provide a transcriptional atlas with a high-resolution annotation that will serve as the basis for future studies of early seed development. To this end, we have created an online resource for exploring the data at https://seedatlas.wi.mit.edu/.

## Acknowledgements

We thank Souraya Khouider and Elizabeth Hemenway for assistance with timed seed isolation for snRNA-seq. We thank Troy Whitfield, Sumeet Gupta, and Alex Dionisio for guidance on module score statistical analyses, snRNA-seq analysis, and seed atlas app development, respectively. Additionally, we thank Jennifer Love, Stephen Mraz III, and Amanda Chilaka at the WIBR Genome Technology Core for all snRNA-seq library preparation and sequencing. We also thank Patrick Autissier and Aditya Rathee for performing FANS procedures at the WIBR flow core facility. This research was supported by The Manton Foundation, The Dr. Vincent J. Ryan Orphan Plant Project, and a National Science Foundation Graduate Research Fellowship to CAM. MG is an Investigator of the Howard Hughes Medical Institute.

## Author Contributions Statement

CAM and MG conceived this study. CAM performed the snRNA-seq and HCR experiments and imaging with help from ALP and KRC. CAM performed all data analyses and CAM and MG interpreted results. The manuscript was prepared by CAM and MG.

## Online Methods

### Generation of single nucleus transcriptomes

#### Plant material

All plants (*Arabidopsis thaliana* (L.) Heynh., Col-0) used in HCR and snRNA-seq experiments were grown at 22°C in a glasshouse on soil under long-day conditions (16 h: 8 h, light: dark). For timed crosses, we emasculated flower buds and after two days pollinated stigma with mature papillae.

#### Tissue preparation and nuclei extraction for snRNA-seq

Two biological replicates (different plants hand-pollinated and processed for snRNA-seq on different days), each containing seeds isolated from 10-15 siliques (500-800 seeds), were collected for each timepoint, producing six replicates total. Seeds were dissected into 150 μL cold extraction buffer on ice (1x Partec CyStain UV Precise P nuclei extraction buffer (Sysmex #05-5002), 4% BSA, 1mM DTT, 1:100 protease inhibitor cocktail for plants (Sigma #P9599), and 1:30 Protector RNase inhibitor (Fisher Scientific #NC1877809)). Seeds were mechanically dissociated in 1.5 mL tubes by grinding with an Axygen pestle (Corning #PES-15-B-SI) for 10 turns. The nuclei suspension was filtered through a 30 μm cell strainer (Fisher Scientific # NC9682496), prewet with Partec CyStain UV Precise P staining buffer (Sysmex #05-5002), into a 5 mL tube for fluorescence activated nuclei sorting (FANS). The strainer was rinsed with Partec CyStain UV Precise P staining buffer into the 5 mL tube to bring the final suspension volume to 1 mL. 1 μL of 1mg/mL DAPI (ThermoFisher Scientific #62248) was added to increase nuclear signal. Nuclei were purified and concentrated by FANS on a BD FACS ARIA Cell Sorter using a 70 μm nozzle chip. We gated on 2C, 3C, 4C, 6C, 8C, and 16C peaks (Supplementary Fig. 9). Nuclei were sorted into 30-50 μL collection buffer (PBS-4% BSA) in a 1.5 mL tube and concentration was assessed using the ARIA nuclei count and by manual counting on a Neubauer Improved hemocytometer. Each biological replicate was processed individually using one Chromium Next GEM Single Cell 3’ Kit (v3.1) reaction with a target nuclei recovery of 10k, except for one 3 DAP biological replicate, which was split across two reactions. Libraries were sequenced on a NovaSeqS2 with 50×50 paired- end reads. In total, 7 libraries were generated in this study.

### Computational analysis of single-nucleus transcriptomes

#### Raw data pre-processing, integration and clustering

Raw sequencing data were processed using Cell Ranger v7.1.0 (10X Genomics). The *Arabidopsis thaliana* TAIR10 genome sequence (Athaliana_447_TAIR10.fa) and the Araport11 annotation (Araport11_GFF3_genes_transposons.filtered.201606.gtf)^108^ were used as inputs to “cellranger mkref” v6.0.02. “cellranger count” was used with default parameters and STAR 2.7.1a.

All scripts for the following analysis steps are deposited in the Gehring Lab GitHub: https://github.com/Gehring-Lab/seed_atlas_2025.

The “filtered_feature_bc_matrix” for each library was individually preprocessed with the “per_library_QCs.R” script, which implements Seurat v5.0.0 for object generation and SoupX v1.6.2 for background RNA correction^109,110^. Genes detected in less than 10 nuclei were removed, and profiles with less than 250 genes filtered out. Following the recommendation from Heumos et al. 2023, we further identified and removed “outlier” nuclei, defined as those with a gene/nucleus or a transcript/nucleus that differs by 5 median absolute deviations from the rest of the sample^111^. We then performed an initial clustering analysis and removed clusters that had significantly lower genes/nucleus than the mean. After low-quality profile removal, we ran scDblFinder v1.12.0 on each library in random mode to generate a per-nucleus doublet score, and removed putative doublets based on the recommended threshold generated by scDblFinder, outlined in the “score_doublets.R” script^112^.

The preprocessed libraries were then merged and further quality controlled, outlined in the “merge_libraries_and_batch_effects_harmony.R” script, which is executed in two rounds. In the first round, libraries are merged by timepoint and an initial clustering analysis is performed to generate cluster-level QCs. Additionally, the JackStraw and ScoreJackStraw functions in Seurat are used to detect statistically significant principal components (PCs) for downstream analysis^110^. Then, the clusters are visually assessed for batch effects. In the second round, we perform a final dimensionality reduction involving log normalization followed by a principal component analysis using 3000 highly variable genes determined by the FindVariableFeatures function of Seurat, the number of PCs determined by JackStraw analysis, k-nearest neighbor graph construction, and UMAP projection. Timepoints that exhibited clustering by biological replicate were integrated with Harmony 1.2.0, which was required for 5 DAP (Supplementary Fig. 9)^113^. At this step, cell cycle genes from Picard et al. 2021 and Menges et al. 2005 were used to categorize nuclei into G0, G1, G1/S, S, G2, and G2/M-phases, if possible, using a modified version of the CellCycleScoring function in Seurat (Extended Data 1, Supplementary Table 11)^16,114^.

To identify an appropriate initial clustering resolution for each timepoint, we performed a parameter sweep of the Seurat FindClusters function, varying the resolution parameter from 1 to 2 in increments of 0.1 (see the “clustering.R” script). For each resolution, an average cluster purity and silhouette score was calculated using the neighborPurity and approxSilhouette bluster 1.8.0 functions, respectively^115^. For each timepoint, we selected a resolution that preceded the greatest drop in neighborhood purity on a resolution vs. average neighborhood purity plot. This clustering, referred to as the “*de novo”* clustering, served as the basis for the level 3 annotation (L3), the highest annotation resolution (Fig. 1D, Supplementary Fig. 1, Supplementary Table 3).

#### Cluster annotation

Using both published markers (Supplementary Table 1) and markers identified in this study with HCR-RNA FISH validation (Supplementary Table 2), we classified all clusters, giving each L3 a full name that includes a number, descriptor, and most informative marker gene(s) (Supplementary Table 3 and Extended Data 3, see “manual_annotation.R” script). *RALFL3*+ nuclei, localized to the CZE cyst by HCR RNA-FISH, were manually annotated based on *RALFL3* expression in the 3-5 DAP datasets. We performed an endosperm-only clustering analysis to isolate nodule and nodule-like clusters at 3 and 5 DAP using markers identified in Picard et al. 2021^16^. Using the Seurat FindClusters function, the nodule and nodule-like populations separated at resolutions 1.6 and 2.4 in 3 and 5 DAP, respectively. We appended the basal cyst, nodule nodule-like annotations to the full 3-5 DAP datasets (Supplementary Fig. 2-3). We also performed a subclustering analysis on the 3 and 5 DAP embryo to identify characterized embryo subpopulations. We selected a clustering resolution for the Seurat

FindClusters function that separated upper and lower protoderm populations in each embryo dataset, which was 1.2 and 1.5 for 3 and 5 DAP, respectively (Extended Data 2). Suspensor nuclei were annotated in the 3 DAP embryo based on enrichment for suspensor markers curated in Kao et al. 2021: nuclei in the 90^th^ percentile of suspensor gene enrichment were classified with as suspensor nuclei (see the “subclustering_embryo_reviews.R” script)^27^. We also re-annotated putative G2/M oi nuclei that initially clustered with the 3 DAP embryo in a separate subclustering analysis (Supplementary Fig. 4). To do this, we re-scaled the 3 DAP expression data with G2/M- and S-phase enrichment scores regressed out (for example, using Seurat::ScaleData(dap3_seurat_object, features = VariableFeatures(dap3_seurat_object), vars.to.regress = c(“S.Score”, “G2M.Score”))), and performed k-means clustering with the same optimal resolution identified in the non-regressed object. We found that the embryo subclustered into two populations in the cell-cycle-regressed object: one which expressed the embryo marker *PDF1,* and one that expressed the oi1 marker *MYB11*. Both of these clusters had high G2/M scores. We subsequently re-annotated the *MYB11+* embryo subcluster as G2/M oi1 in the non-cell-cycle regressed dataset (Supplementary Fig. 4). In all timepoint datasets, clusters with negligible differential expression differences in the oi1 were merged into one cluster. We annotated timepoints separately and integrated them into a final atlas dataset using robust principal component analysis (RPCA), see “atlas_merging_rpca.R” script^110^.

#### GO term and gene module analysis with statistical testing

Based on GO term analysis for all DE genes for each atlas cluster across all annotation levels using clusterProfiler 4.7.1.002 (see “clusterprofiler_intermediate.R” script), we identified gene sets (“modules”) for follow-up enrichment analysis^116^. We retrieved gene lists for GO terms from uniprot (https://www.uniprot.org/) using GO term IDs filtered by the *Arabidopsis thaliana* taxon ID (taxonomy_id:3702). Gene lists were used as input to the Seurat AddModuleScore function with default settings (number of bins, “nbin” = 24; number of control genes per bin, “nctrl” = 100), which implements the gene set enrichment approach described in Tirosh 2016^110,117^. The resulting “module score” is the difference in average expression between the gene set of interest and a randomly-generated gene set with matched expression level variability. All gene lists are deposited with the scripts used for generating scores, which include “signalling_transport_gene_enrichment.R”, “peptide_enrichment.R”, and “protein_catabolism_enrichment.R”. The same approach was used for curated gene lists of SSPs, BZR/BES1 transcription factors, and embryo subtype marker genes.

To statistically test module scores by cluster, we used the Wilcoxon Rank-Sum test in focal cluster vs. all other nuclei comparisons. After performing this test for all clusters for a given module score, we adjusted p-values using Bonferroni correction. See Supplementary Table 7 for the results of all statistical analyses of module scores performed in this study.

#### Correlation analysis of pseudobulked clusters

To quantify similarity between cell types, we pseudobulked clusters using the Seurat function AggregateExpression using default arguments (normalization.method = “LogNormalize”, scale.factor = 10000) and the top 3000 variable genes for the snRNA-seq datasets of interest^110^. We used the pseudoexpression matrices as input to generate correlation matricies using Spearman correlation. To characterize gene expression correlation between biological replicates, we used the same approach, but pseudobulked all genes detected and used Pearson correlation.

#### Pseudotime analysis

To define the transcriptional landscape of CZE development, we performed pseudotime analysis on integrated 3-5 DAP PEN and CZE nuclei. We merged 3 and 5 DAP PEN and CZE data, then performed dimensionality reduction analysis on the subset, regressing out cell cycle genes during data scaling and centering. Datasets were then integrated across timepoints using Harmony. To identify only one trajectory, we assigned a single partition to all nuclei in each dataset, and manually specified the 3 DAP PEN as the root for trajectory and pseudotime inference. Nuclei were ordered and assigned a pseudotime value using the “learn_graph” and “order_cells” functions in monocle3 1.3.7^118^. To identify genes that vary in pseudotime, we implemented graph autocorrelation analysis using the graph_test function in monocle3 1.3.7^118^. See “level_2_pseudotime_merged_timepoint.R” and “harmony_chalazal_endosperm_trajectory.R” scripts for details^118^.

#### Identifying differentially expressed genes

Differential expression analysis was performed on each timepoint and the integrated atlas object using the Seurat FindAllMarkers function with default arguments, generating cluster vs. all other nuclei results for each gene and p-values from a Wilcoxon Rank Sum test with Bonferroni correction, see “differential_expression.R” script for details. To determine whether a gene is upregulated after fertilization (Fig. 5C), we re-analyzed the published expression data from four day after emasculation unfertilized ovules and 2 DAP seeds (“GSE85751_RMA_matrix.txt”) from Figueiredo et al. 2016^11^, using limma^119^ with Benjamini-Hochberg correction to calculate the significance of differentially expressed genes. Genes with a Log_2_FC greater than 0.5 with an adjusted p-value of less than or equal to 0.05 in the fertilized condition were classified as upregulated.

### Marker validation using HCR RNA-FISH

To localize clusters and marker genes *in situ*, we used HCR-RNA FISH based on a modified version of the whole mount HCR protocol outlined in Huang et al. 2023 and the general HCR-RNA FISH protocol developed by Molecular Instruments: https://www.molecularinstruments.com/hcr-gold-rnafish^120^. All HCR probes and fluorescent hairpins were synthesized by Molecular Instruments (Supplementary Table 2). Hand-pollinated or unstaged siliques were collected and fixed with 4% paraformaldehyde (PFA) in 1X PBS through vacuum infiltration. After overnight fixation at 4 °C, samples were washed in 1x PBS twice before embedding in 3% agarose for vibratome sectioning. Whole siliques were sectioned longitudinally to 60-150 μm thickness and stored in PBS on ice. To generate negative controls in parallel with each probe set, we stored sections in two tubes (one tube for the experimental and one for the negative control) during slicing, alternating tube placement every other section. Sections were subject to a second 4% PFA fixation for 30 minutes at room temperature before two washes in PBS, then transferred to 100% methanol and stored at -20 °C. To permeabilize tissue before probe hybridization, samples were subjected to alternating ethanol and methanol incubations, following Huang et al. 2023, with a clearing step using 50% Histoclear (National Diagnostics #HS-200)/50% ethanol halfway through the permeabilization washes^120^. Samples were rehydrated with an ethanol/PBS-Tween-20 series (25%, 50%, 75%, 100%). Then, samples were incubated in preheated Probe Hybridization Buffer (Molecular Instruments) for 30 minutes. Sections in the experimental tube were then exchanged into preheated Probe Hybridization Buffer (Molecular Instruments) with probes, and sections in the negative control tube were exchanged into preheated Probe Hybridization Buffer without probes. For targets with < 5 probe sets, we used 64 μL of 1 μM probe in 400 μL Probe Hybridization Buffer, and for all other probes we used 12-24 μL probe 1 μM in 400 μL. Probes were hybridized to samples in 1.5mL tubes overnight in a 37 °C water bath. Probe washes closely followed the Molecular Instruments HCR protocol: we performed four 15-minute buffer exchanges with Probe Wash Buffer (Molecular Instruments) preheated to 37 °C in a water bath, followed by two quick washes with 5X SSC-Tween-20. Samples in both positive and negative control tubes were incubated in Amplification Buffer (Molecular Instruments) for 30 minutes at room temperature before exchange to Amplification Buffer containing snap-cooled hairpins (8 μL of 3 μM stock per hairpin per 400 μL reaction volume). We used hairpins containing B2 and B3 adapter sequences with Alexa-488 or Alexa-647 dyes (Molecular Instruments). After an overnight incubation in the dark, excess probes were washed off with four 5x SSC-Tween-20 exchanges. Samples were stored for up to two weeks at 4 °C before imaging.

To prepare samples for confocal microscopy, we counterstained nuclei with 1μg/mL DAPI and mounted silique sections on thin 2% agar pads suspended in water on glass slides. We imaged Z-stacks of samples on a Zeiss LSM 710 confocal and generated maximum intensity projection images using FIJI.

Whereas our negative controls indicate that the presented HCR results were not derived from spurious signals, it is possible that our HCR probes non-specifically bind to other RNAs or tissue components. This issue would be resolved if sense probes designed to mRNA targets were used as an additional negative control.

### SSP detection and annotation

To detect all SSPs in the Arabidopsis genome, we filtered all Araport11 protein sequences for those less than 250 residues and used this as input to SignalP6 to identify those with N-terminal secretory signal sequences, see “signalp6_command.sh” for details. We transferred the SSP annotations described in Ghorbani et al. 2015 to the SSPs detected by SignalP6^88,121^. For all SSPs detected by SignalP6 expressed in the atlas that did not have an annotation from Ghorbani et al 2015, we used the Araport11 annotation categorize these sequences into SSP families (Supplementary Table 10).

#### Maximum likelihood estimation of positive selection at sites in *A. thaliana* single-copy orthologs

To implement site models of codon evolution using codeml/PAML 4.9^97^, we closely followed the procedure outlined in the Bioinformatics Workbook^122^, based on the analyses in Petersen et al. 2007^123^. We identified orthologs between the translated coding sequences (CDS) of *Arabidopsis thaliana (*Araport11, Phytozome genome ID: 447), *Arabidopsis lyrata (*v2.1, Phytozome genome ID: 384), *Arabidopsis arenosa* (AARE701a), and *Capsella grandiflora* (v1.1, Phytozome genome ID: 266) using OrthoFinder 2.5.4^124^. Arabidopsis single-copy orthologs that were found in all four species were used for subsequent analysis. Protein sequences from gene trees generated by Orthofinder were aligned with clustalo^125^. The protein alignments and CDSs from each gene tree were used as input to the pal2nal.pl script^126^ to produce codon alignments, omitting gaps in all sequence alignments (pal2nal.pl -nogap). Codon alignments and pruned gene trees were used as input to codeml, with arguments for performing maximum likelihood estimation of site models of codon evolution (runmode = 0, seqtype = 1, CodonFreq = 2, model = 0, NSsites = 0 1 2 7 8, fix_kappa = 0, kappa = 2, fix_omega = 0, omega = 0.4, cleandata = 1). We calculated both M2a/M1a and M8/M7 LRTs (2*(lnL_alt_ – lnL_null_)), but proceeded with M2a/M1a for higher stringency. See “orthofinder_to_codeml.sh” script for more details. LRTs were significance tested using the chi square test function pchisq in R. All p-values were adjusted within M2a/M1a comparisons using Benjamini-Hochberg correction. The Bayes Empirical Bayes results from M2a/M1a analyses for individual sites were extracted from codeml outputs and evaluated only for genes with significant M2a/M1a LRTs.

To identify protein domains that overlap with positively-selected sites in DE M2a/M1a genes, we performed an InterProScan (https://www.ebi.ac.uk/interpro/) for all translated DE M2a/M1a CDSs. We downloaded the resulting GFF and matched selected sites to protein domains using a custom R script, see “analyzing_codeml.R” for details.

## Data Availability

All sequencing data is available in NCBI GEO GSE295007. A browser for the data is available at https://seedatlas.wi.mit.edu/

## Code Availability

Scripts for analyses are deposited in GitHub at https://github.com/Gehring-Lab/seed_atlas_2025.

---

**Extended Data 1:**
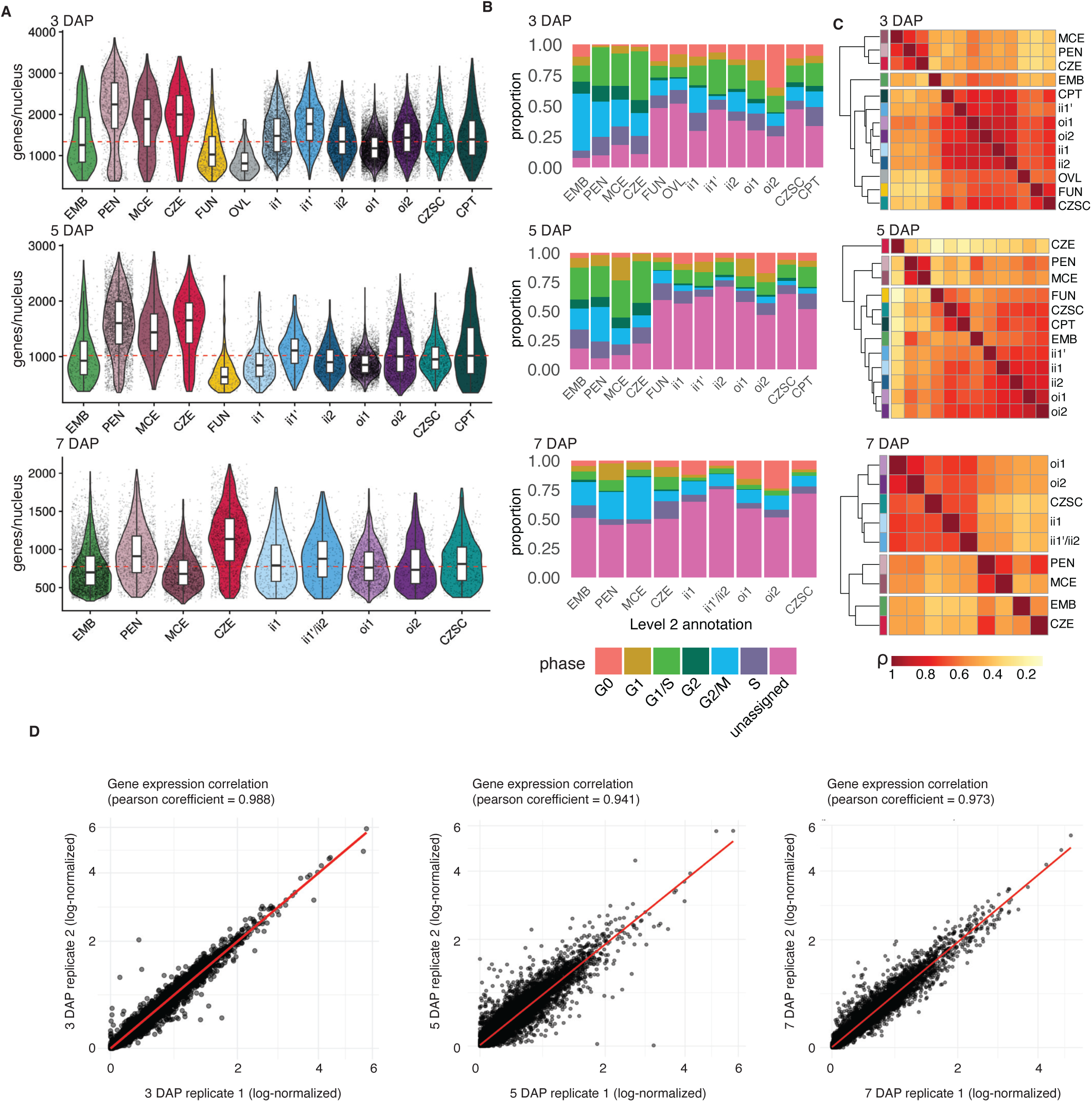
General cluster characteristics and similarity. **A**, Genes per nucleus distributions for level 2 (L2) annotations across all timepoints. Red dotted line drawn at the median. **B**, Cell cycle stage proportionality for L2 clusters across all timepoints based on cell cycle phase marker genes defined in Picard et al. 2021 and Menges et al. 2005. **C**, Clustered heatmaps of the Spearman correlations for the aggregated expression of 3000 highly variable genes across all L2 annotations and timepoints. **D**, Pearson correlation of aggregated gene expression between two biological replicates for each timepoint (3 DAP: 21755 genes, 5 DAP: 21227 genes, 7 DAP: 19705 genes). Axes are pseudo log-transformed.

**Extended Data 2:**
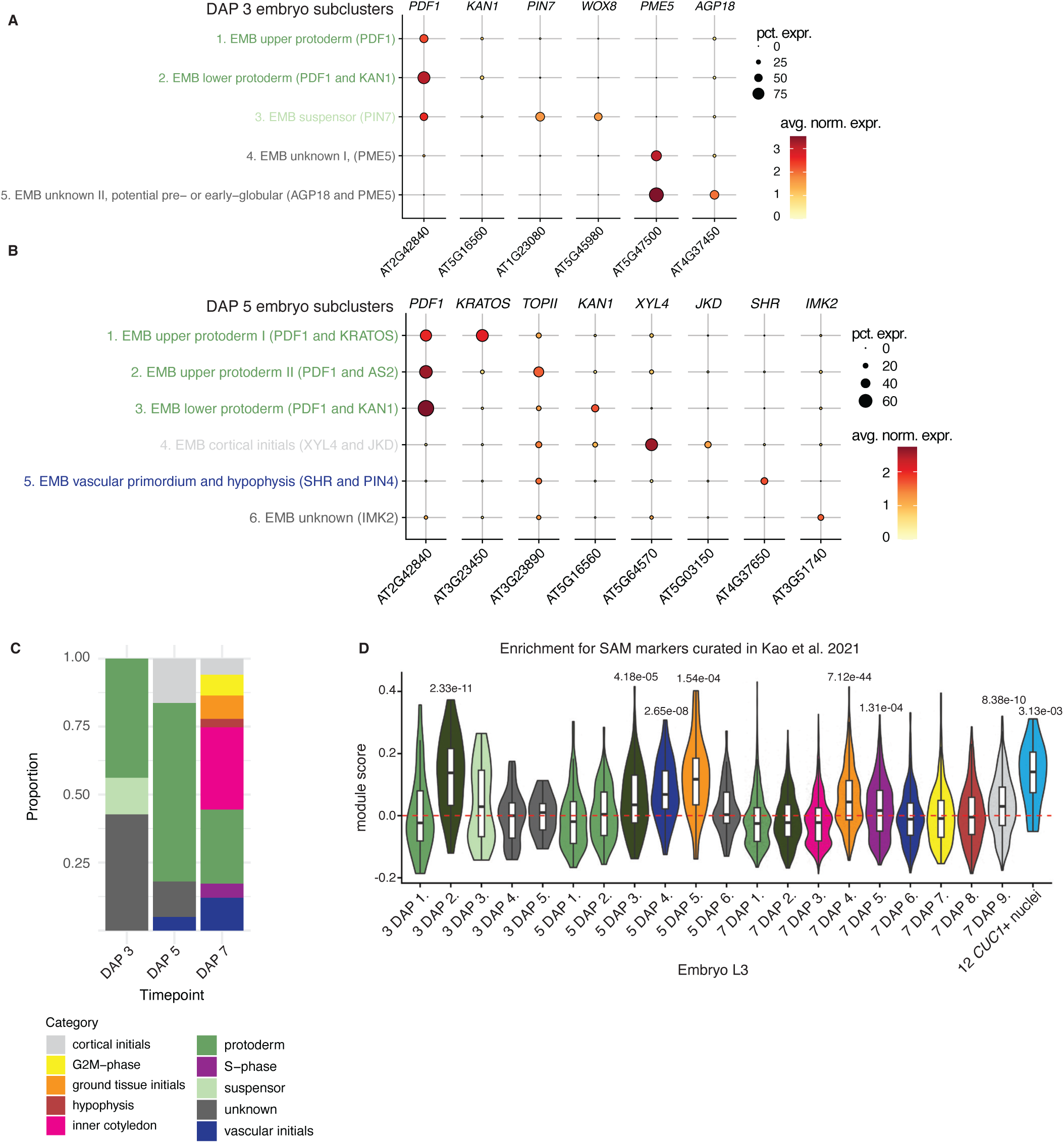
Embryo subclustering analysis. **A-B**, The most informative markers for clusters identified in the 3 DAP and 5 DAP embryo, which include markers with characterized embryonic expression patterns (*PDF1, KAN1, AGP18, SHR, PIN4, XYL4, JKD*, and *PIN7*), and those that are specific to embryo subclusters in this dataset (*PME5, KRATOS, IMK2*, and *TOPII*). **C**, Proportions of embryo subcluster categories at each timepoint. **D**, Module score analysis for 40 strong shoot apical meristem (SAM) markers curated in Kao et al. 2021 across all embryo L3 clusters and CUC1+ nuclei. p-values are centered above clusters with significantly high positive module scores in a cluster-vs-all other nuclei comparison. *p*-values are derived from a two-sided Wilcoxon Rank-Sum test with Bonferroni correction. See Supplementary Table 7 for the module scores and p-values for all clusters.

**Extended Data 3:**
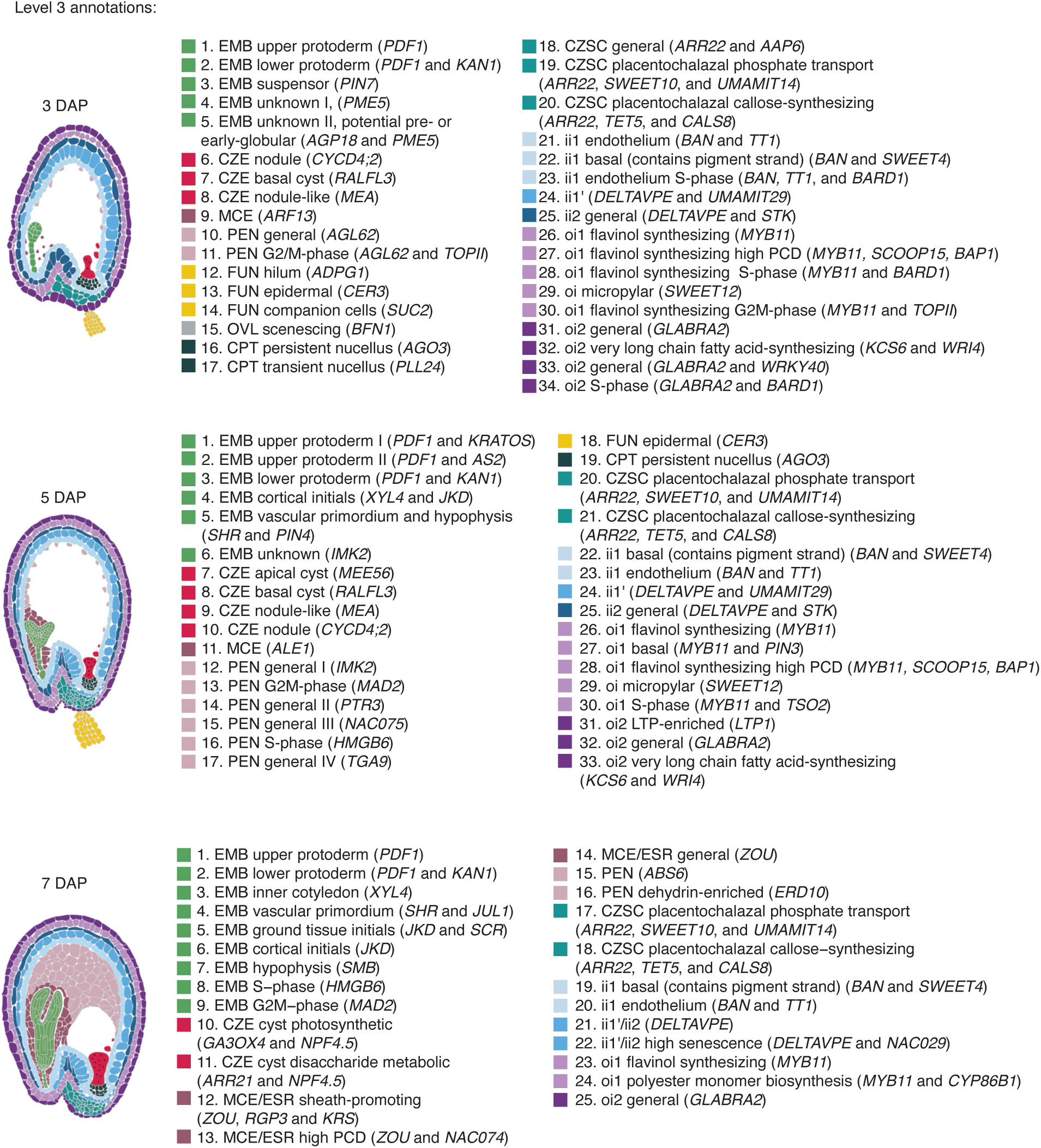
Level 3 annotation key. Full names for level 3 (L3) annotations for all timepoints, with defining marker genes in parenthe-ses. EMB, embryo; CZE, chalazal endosperm; MCE, micropylar endosperm; PEN, peripheral endosperm; FUN, funiculus; OVL, ovule 5 days after emasculation; CPT, chalazal proliferating tissue; CZSC, chalazal seed coat; ii1, inner integument 1; ii1’, inner integument 1’; ii2; inner integument 2; oi1, outer integument 1; oi2, outer integument 2; ESR, embryo-surrounding region; LTP, lipid transfer protein; PCD,programmed cell death.

**Extended Data 4:**
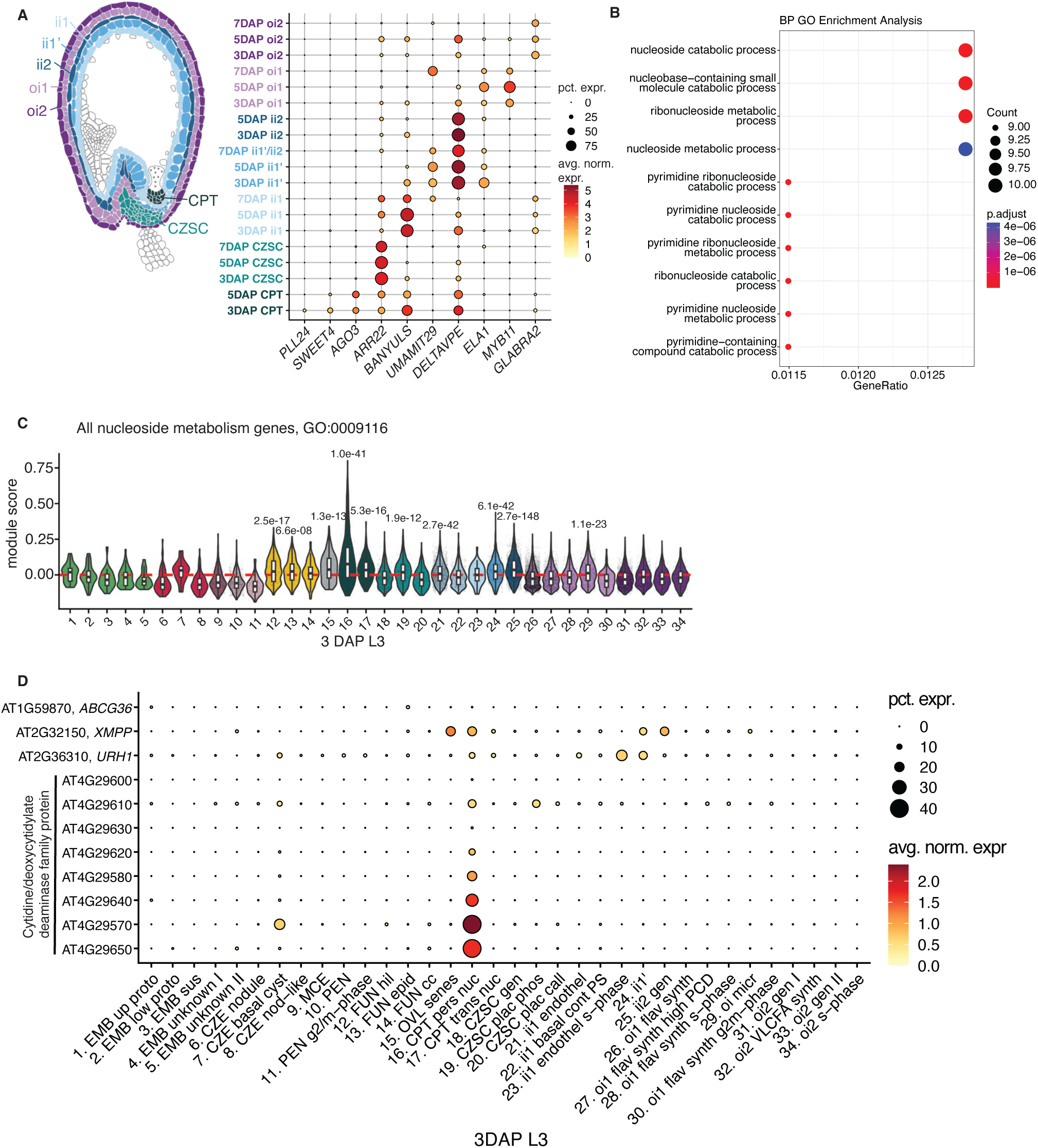
Informative markers for seed coat layers and nucleoside metabolism enrichment in the chalazal proliferating tissue. **A**, The most informative markers for L2 seed coat layers. All markers except MYB11 have previously characterized layer-specific expression patterns. *MYB11*, a regulator of flavanol biosynthesis, is the top marker for oi1 in this dataset. **B**, The most significant GO terms for the 3 DAP CPT, based on differentially expressed genes in a cluster vs. all 3 DAP clusters comparison (log2FC > 1, adj. p-value <0.05). **C**, Module score results for all nucleo-side metabolism genes in 3 DAP clusters. p-values are centered above clusters with significantly high average positive module scores in a cluster-vs-all other nuclei comparison. p-values are derived from a two-sided Wilcoxon Rank-Sum test with Bonferroni correction. **D**, The expression patterns of the top genes that underlie nucleoside metabolism enrichment in the 3 DAP persistent CPT.

**Extended Data 5:**
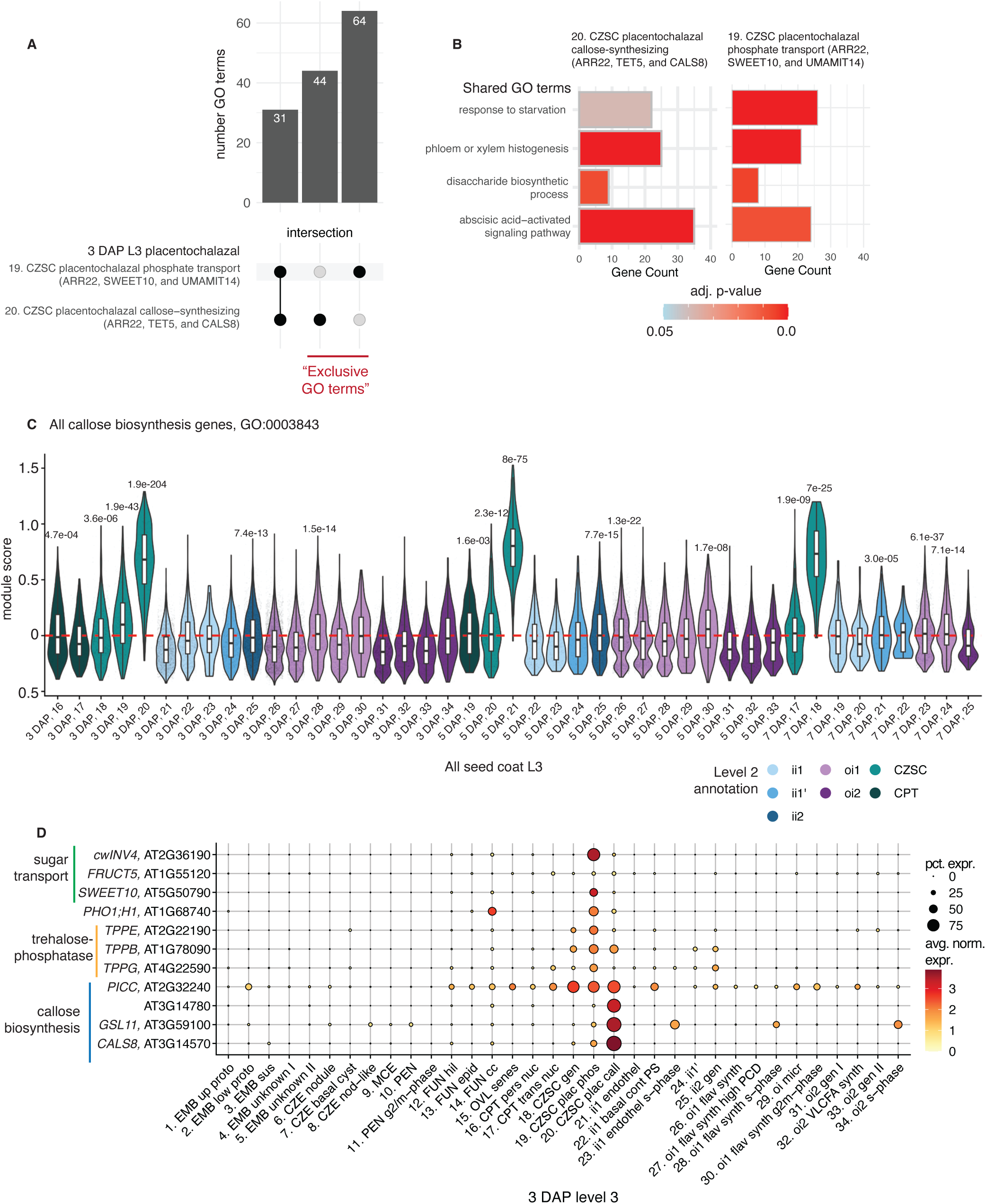
Cell types with complementary functions in the 3 DAP CZSC. **A**, GO term overlap for genes DE (log2FC > 1, adj. p-value <0.05) in two putative placentochalazal clusters. Exclusive GO terms are those associated with only one of the clusters. **B**, A subset of shared GO terms for the two putative placentochalazal clusters at 3 DAP. **C**, Module score analysis for all genes associated with the GO term “callose biosynthesis” shown across all L3 seed coat clusters in the atlas (Methods). p-values are centered above clusters with significantly high average positive module scores in a cluster-vs-all other nuclei comparison. p-values are derived from a two-sided Wilcoxon Rank-Sum test with Bonferroni correction, see Supplementary Table 7 for the module scores and p-values for all clusters. **D**, Expression patterns of genes that underlie exclusive GO terms for the two putative CZSC clusters at 3 DAP. See Supplementary Table 3 to match abbreviated L3 names to their full descriptions

**Extended Data 6:**
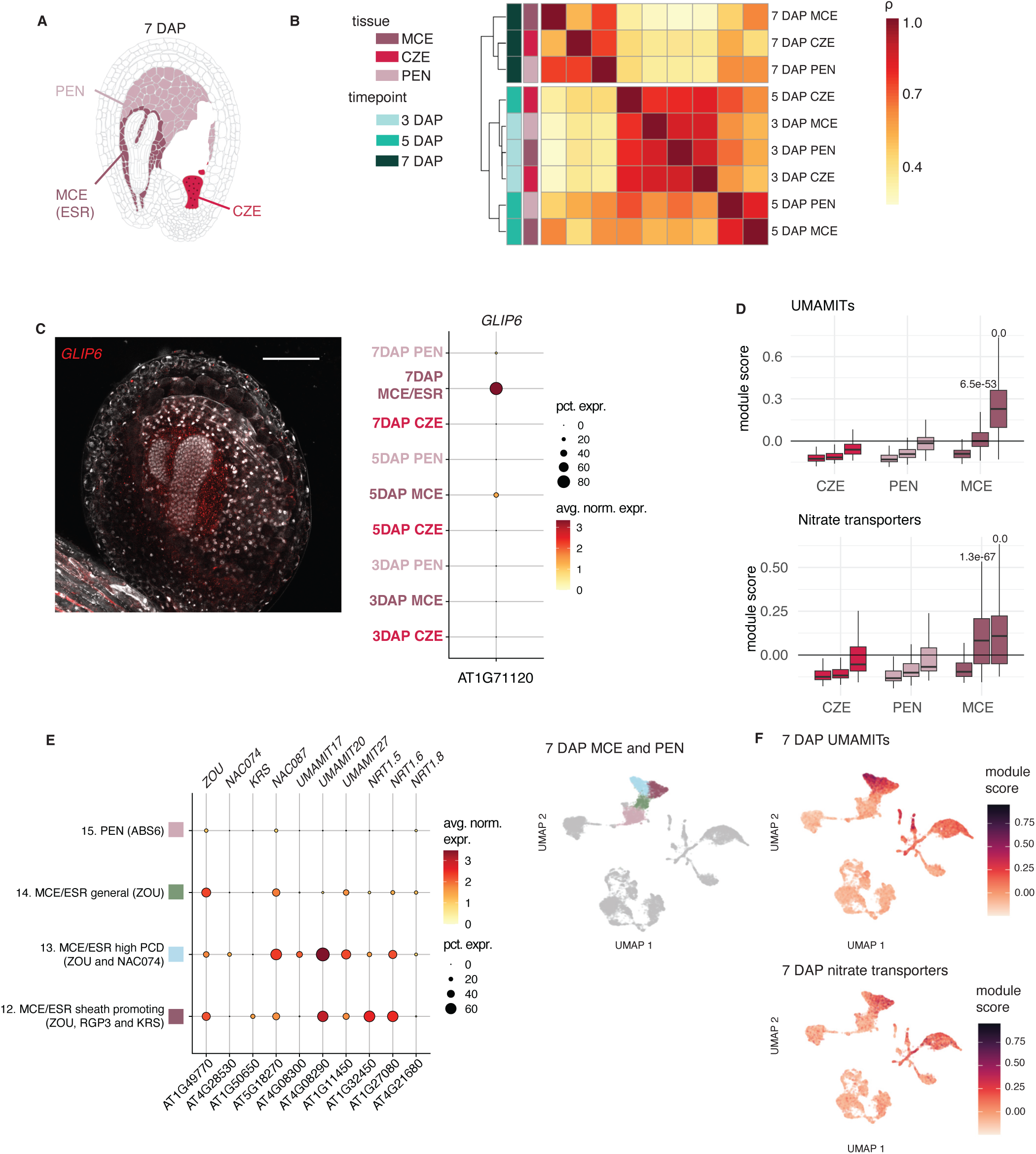
The 7 DAP MCE is the most distinctive endosperm subtype. **A**, L2 endosperm subtypes at 7 DAP. **B**, A clustered heatmap of the Spearman correlation coefficient (ρ) of aggregated expression for L2 endosperm clusters split by time. **C**, *GLIP6* is a highly specific marker for the ESR at 7 DAP, validated by HCR.Scale bar = 100 µm. **D**, Module score analysis for all UMAMITs (top) and nitrate transporters (bottom) detected in the atlas for all L2 endosperm clusters. p-values are centered above clusters with significantly high average positive module scores in a cluster-vs-all other nuclei comparison. p-values are derived from a two-sided Wilcoxon Rank-Sum test with Bonferroni correction, see Supplementary Table 7 for the module scores and p-values for all clusters. **E**, 7 DAP L3 MCE clusters show differential expression of *KRS* and two NAC transcription factors. **F**, Nitrate transporters follow the *KRS-NAC074* transcriptional gradient while *UMAMITs* are more broadly expressed.

**Extended Data 7:**
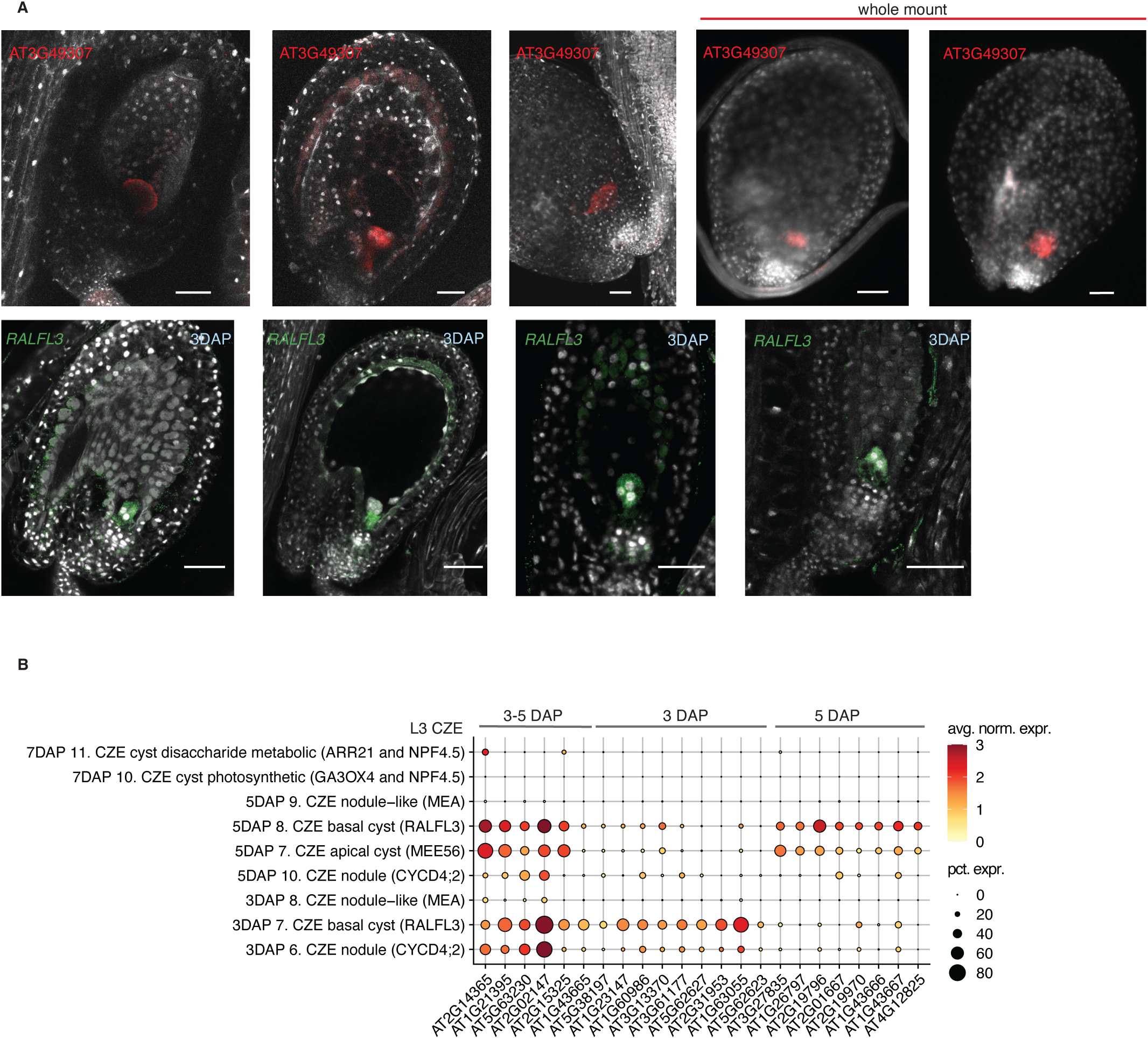
Differentially expressed genes along the apical/basal axis of the CZE. **A**, Top row: HCR for AT3G49307 (red), a gene enriched in the apical CZE during early to intermediate stages of development based on snRNA-seq data. These samples were not staged. The last two images are from whole mount seed preparations (Methods). Bottom row: HCR validation of *RALFL3* (green), a basal cyst marker, assayed on 3 DAP seeds. DAPI staining in white. Scale bar = 50 µm. **B**, Expression patterns for the top basal cyst markers, excluding those in Fig. 4D, through development within L3 CZE subtypes.

**Extended Data 8:**
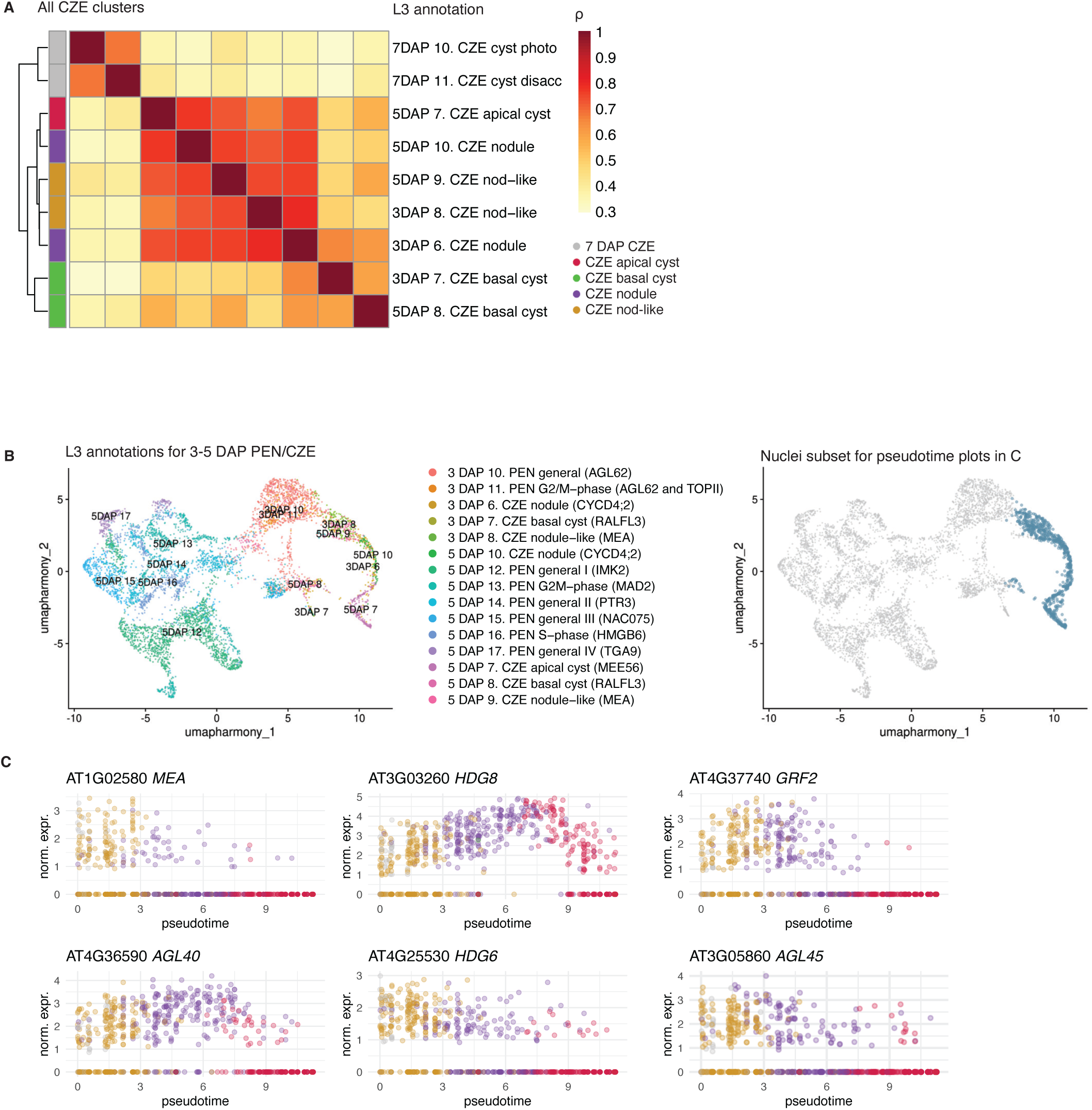
Genes that vary along the CZE nodule-like-to-cyst trajectory. **A**, Spearman correlation coefficients of the pairwise similarity between aggregated expression profiles of L3 CZE subtypes. 7 DAP L3 clusters could not be assigned apical/basal states. See Supplementary Table 3 to match abbreviated L3 names to their full descriptions. **B**, Left: L3 annotations for 3-5 DAP PEN and CZE clusters used in pseudotime analysis. Right: only blue-highlighted nuclei were used in the pseudotime expression plots in C to show gene expression variation in pseudotime along the non-basal cyst PEN to CZE trajectory (see Fig. 4). **C**, Normalized expression of a subset of genes that significantly vary on 3-5 DAP PEN/CZE pseudotime (see Fig. 4) is plotted in pseudotime-ordered nuclei. Color key in A extends to the labeling in C.

**Extended Data 9:**
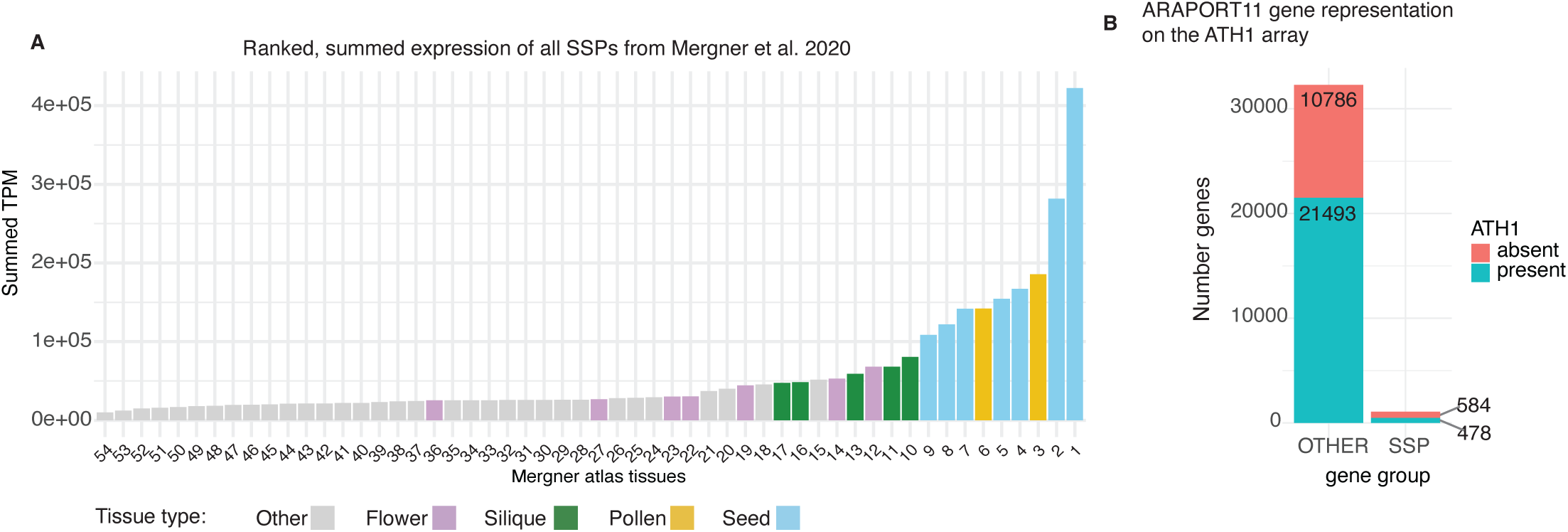
SSPs are highly expressed in seed tissues and were absent from existing transcriptional atlases of seed development. **A**, Ranked, summed TPM of all SSP genes used in this study using the bulk RNA-sequencing data of Mergner et al. 2020. See Supplementary Table 9 for sample names. **B**, ARAPORT11 gene annotation coverage of the Affymetrix ATH1 array.

**Extended Data 10:**
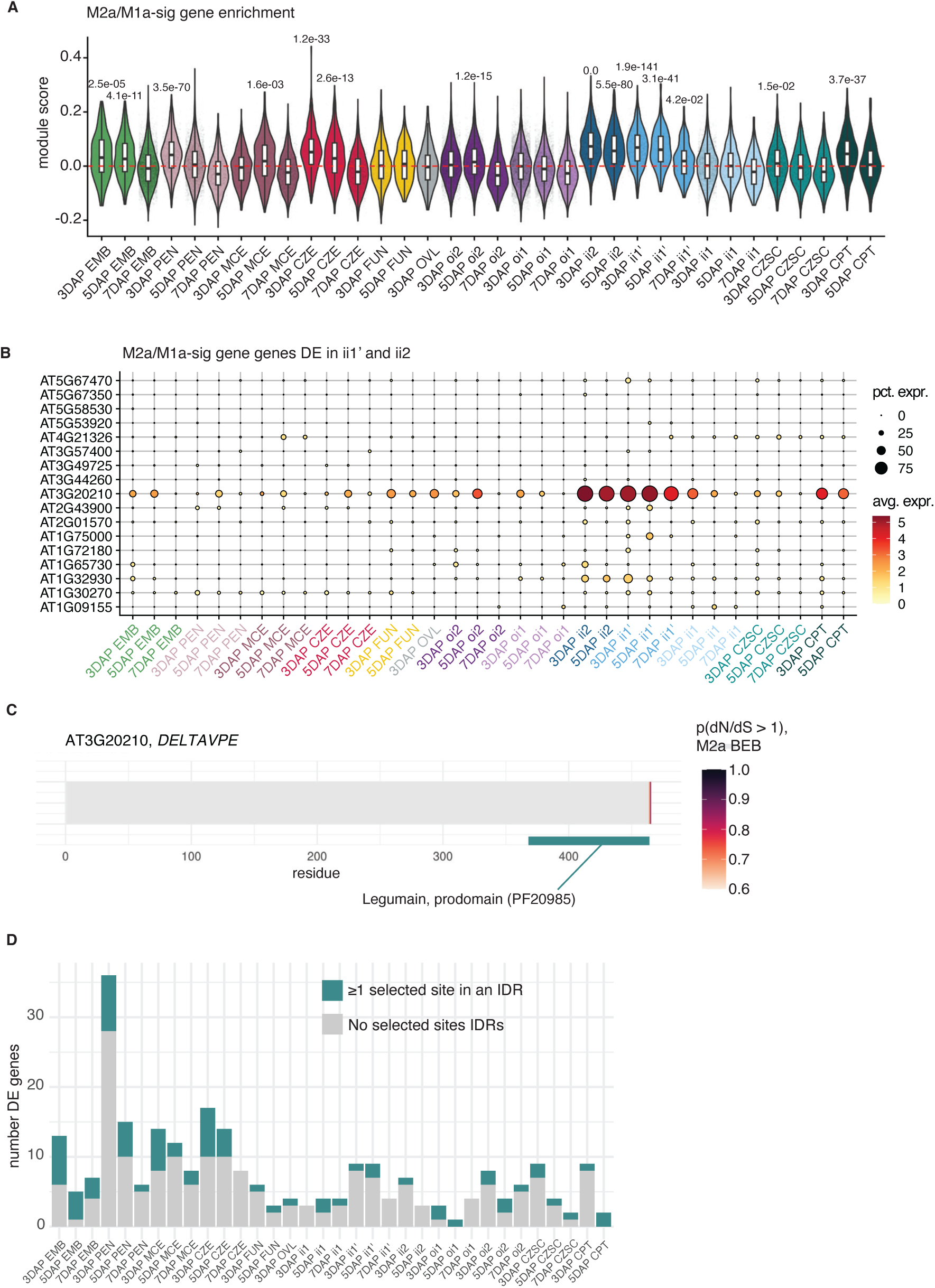
Rapidly evolving single-copy ortholog gene expression enrichment in endosperm and seed coat subtypes. **A**, Module score analysis for single-copy orthologs (SCOs) with significant M2a/M1a likelihood ratio tests(”M2a/M1a-sig”) across L2 timed atlas clusters. p-values are derived from a two-sided Wilcoxon Rank-Sum test with Bonferroni correction in focal cluster vs. all other comparisons. Only p-values for clusters with positive, significant module scores are shown. See Supplementary Table 7 for the module scores and p-values for all clusters. **B**, The expression patterns for M2a/M1a-sig SCOs differentially expressed in ii1’ and ii2 subtypes (p-value ≤ 0.05, log2FC > 1). **C**, The residues likely under positive selection in the DELTAVPE protein sequence. Individual residues colored by the Bayes Emprical Bayes (BEB) posterior probability of having a dN/dS > 1 under the M2a model. Informative protein domains near or containing selected sites are highlighted, Pfam identifier fromInterProScan in parentheses. **D**, The number of DE M2a/M1a-sig genes (p-value ≤ 0.05, log2FC > 1) per L2 timed cluster. Individual genes are colored blue-green if at least one selected site falls in an an intrinsically disordered region (IDR) as predicted by MobiDB-lite.

## References

(1) Nowack, M. K.; Ungru, A.; Bjerkan, K. N.; Grini, P. E.; Schnittger, A. Reproductive Cross-Talk: Seed Development in Flowering Plants. Biochem. Soc. Trans. 2010, 38 (2), 604–612. 10.1042/BST0380604.

(2) Berger, F. Endosperm: The Crossroad of Seed Development. Curr. Opin. Plant Biol. 2003, 6 (1), 42–50. 10.1016/S1369526602000043.

(3) Boisnard-Lorig, C.; Colon-Carmona, A.; Bauch, M.; Hodge, S.; Doerner, P.; Bancharel, E.; Dumas, C.; Haseloff, J.; Berger, F. Dynamic Analyses of the Expression of the HISTONE::YFP Fusion Protein in Arabidopsis Show That Syncytial Endosperm Is Divided in Mitotic Domains. Plant Cell 2001, 13 (3), 495–510.

(4) Becker, M. G.; Hsu, S.-W.; Harada, J. J.; Belmonte, M. F. Genomic Dissection of the Seed. Front. Plant Sci. 2014, 5. 10.3389/fpls.2014.00464.

(5) Khan, D.; Millar, J. L.; Girard, I. J.; Belmonte, M. F. Transcriptional Circuitry Underlying Seed Coat Development in Arabidopsis. Plant Sci. Int. J. Exp. Plant Biol. 2014, 219–220, 51–60. 10.1016/j.plantsci.2014.01.004.

(6) Ingram, G. C. Family Life at Close Quarters: Communication and Constraint in Angiosperm Seed Development. Protoplasma 2010, 247 (3–4), 195–214. 10.1007/s00709-010-0184-y.

(7) Stadler, R.; Lauterbach, C.; Sauer, N. Cell-to-Cell Movement of Green Fluorescent Protein Reveals Post-Phloem Transport in the Outer Integument and Identifies Symplastic Domains in Arabidopsis Seeds and Embryos. Plant Physiol. 2005, 139 (2), 701–712. 10.1104/pp.105.065607.

(8) Doll, N. M.; Ingram, G. C. Embryo-Endosperm Interactions. Annu. Rev. Plant Biol. 2022, 73, 293–321. 10.1146/annurev-arplant-102820-091838.

(9) Robert, H. S.; Park, C.; Gutièrrez, C. L.; Wójcikowska, B.; Pěnčík, A.; Novák, O.; Chen, J.; Grunewald, W.; Dresselhaus, T.; Friml, J.; Laux, T. Maternal Auxin Supply Contributes to Early Embryo Patterning in Arabidopsis. Nat. Plants 2018, 4 (8), 548–553. 10.1038/s41477-018-0204-z.

(10) Figueiredo, D. D.; Köhler, C. Auxin: A Molecular Trigger of Seed Development. Genes Dev. 2018, 32 (7–8), 479–490. 10.1101/gad.312546.118.

(11) Figueiredo, D. D.; Batista, R. A.; Roszak, P. J.; Hennig, L.; Köhler, C. Auxin Production in the Endosperm Drives Seed Coat Development in Arabidopsis. eLife 2016, 5, e20542. 10.7554/eLife.20542.

(12) Lafon-Placette, C.; Köhler, C. Embryo and Endosperm, Partners in Seed Development. Curr. Opin. Plant Biol. 2014, 17, 64–69. 10.1016/j.pbi.2013.11.008.

(13) Xu, W.; Sato, H.; Bente, H.; Santos-González, J.; Köhler, C. Endosperm Cellularization Failure Induces a Dehydration-Stress Response Leading to Embryo Arrest. Plant Cell 2023, 35 (2), 874–888. 10.1093/plcell/koac337.

(14) Song, J.; Xie, X.; Chen, C.; Shu, J.; Thapa, R. K.; Nguyen, V.; Bian, S.; Kohalmi, S. E.; Marsolais, F.; Zou, J.; Cui, Y. LEAFY COTYLEDON1 Expression in the Endosperm Enables Embryo Maturation in Arabidopsis. Nat. Commun. 2021, 12 (1), 3963. 10.1038/s41467-021-24234-1.

(15) Brown, R. C.; Lemmon, B. E.; Nguyen, H.; Olsen, O.-A. Development of Endosperm in Arabidopsis Thaliana. Sex. Plant Reprod. 1999, 12 (1), 32–42. 10.1007/s004970050169.

(16) Picard, C. L.; Povilus, R. A.; Williams, B. P.; Gehring, M. Transcriptional and Imprinting Complexity in Arabidopsis Seeds at Single-Nucleus Resolution. Nat. Plants 2021, 7 (6), 730–738. 10.1038/s41477-021-00922-0.

(17) Belmonte, M. F.; Kirkbride, R. C.; Stone, S. L.; Pelletier, J. M.; Bui, A. Q.; Yeung, E. C.; Hashimoto, M.; Fei, J.; Harada, C. M.; Munoz, M. D.; Le, B. H.; Drews, G. N.; Brady, S. M.; Goldberg, R. B.; Harada, J. J. Comprehensive Developmental Profiles of Gene Activity in Regions and Subregions of the Arabidopsis Seed. Proc. Natl. Acad. Sci. 2013, 110 (5), E435–E444. 10.1073/pnas.1222061110.

(18) van Ekelenburg, Y. S.; Hornslien, K. S.; Van Hautegem, T.; Fendrych, M.; Van Isterdael, G.; Bjerkan, K. N.; Miller, J. R.; Nowack, M. K.; Grini, P. E. Spatial and Temporal Regulation of Parent-of-Origin Allelic Expression in the Endosperm. Plant Physiol. 2023, 191 (2), 986–1001. 10.1093/plphys/kiac520.

(19) Doll, N. M.; Royek, S.; Fujita, S.; Okuda, S.; Chamot, S.; Stintzi, A.; Widiez, T.; Hothorn, M.; Schaller, A.; Geldner, N.; Ingram, G. A Two-Way Molecular Dialogue between Embryo and Endosperm Is Required for Seed Development. Science 2020, 367 (6476), 431–435. 10.1126/science.aaz4131.

(20) Royek, S.; Bayer, M.; Pfannstiel, J.; Pleiss, J.; Ingram, G.; Stintzi, A.; Schaller, A. Processing of a Plant Peptide Hormone Precursor Facilitated by Posttranslational Tyrosine Sulfation. Proc. Natl. Acad. Sci. 2022, 119 (16), e2201195119. 10.1073/pnas.2201195119.

(21) Millar, J. L.; Khan, D.; Becker, M. G.; Chan, A.; Dufresne, A.; Sumner, M.; Belmonte, M. F. Chalazal Seed Coat Development in *Brassica Napus*. Plant Sci. 2015, 241, 45–54. 10.1016/j.plantsci.2015.09.019.

(22) Pegler, J. L.; Grof, C. P.; Patrick, J. W. Sugar Loading of Crop Seeds – a Partnership of Phloem, Plasmodesmal and Membrane Transport. New Phytol. 2023, 239 (5), 1584–1602. 10.1111/nph.19058.

(23) Liu, H.; Luo, Q.; Tan, C.; Song, J.; Zhang, T.; Men, S. Biosynthesis- and Transport-Mediated Dynamic Auxin Distribution during Seed Development Controls Seed Size in Arabidopsis. *Plant J*. Cell Mol. Biol. 2023, 113 (6), 1259–1277. 10.1111/tpj.16109.

(24) Chen, X.; Bezodis, W.; González-Suárez, P.; Knitlhoffer, V.; Goldson, A.; Lister, A.; Macaulay, I.; Penfield, S. Trans-Generational Adaptation to Maternal Climate through Hormone Transport in Plants. bioRxiv October 5, 2024, p 2024.10.04.616646. 10.1101/2024.10.04.616646.

(25) Hibara, K.; Takada, S.; Tasaka, M. CUC1 Gene Activates the Expression of SAM-Related Genes to Induce Adventitious Shoot Formation. Plant J. 2003, 36 (5), 687–696. 10.1046/j.1365-313X.2003.01911.x.

(26) Ueda, M.; Zhang, Z.; Laux, T. Transcriptional Activation of *Arabidopsis* Axis Patterning Genes *WOX8/9* Links Zygote Polarity to Embryo Development. Dev. Cell 2011, 20 (2), 264–270. 10.1016/j.devcel.2011.01.009.

(27) Kao, P.; Schon, M. A.; Mosiolek, M.; Enugutti, B.; Nodine, M. D. Gene Expression Variation in Arabidopsis Embryos at Single-Nucleus Resolution. Dev. Camb. Engl. 2021, 148 (13), dev199589. 10.1242/dev.199589.

(28) Capron, A.; Chatfield, S.; Provart, N.; Berleth, T. Embryogenesis: Pattern Formation from a Single Cell. Arab. Book 2009, 7, e0126. 10.1199/tab.0126.

(29) Wysocka-Diller, J. W.; Helariutta, Y.; Fukaki, H.; Malamy, J. E.; Benfey, P. N. Molecular Analysis of SCARECROW Function Reveals a Radial Patterning Mechanism Common to Root and Shoot. Development 2000, 127 (3), 595–603. 10.1242/dev.127.3.595.

(30) Welch, D.; Hassan, H.; Blilou, I.; Immink, R.; Heidstra, R.; Scheres, B. Arabidopsis JACKDAW and MAGPIE Zinc Finger Proteins Delimit Asymmetric Cell Division and Stabilize Tissue Boundaries by Restricting SHORT-ROOT Action. Genes Dev. 2007, 21 (17), 2196–2204. 10.1101/gad.440307.

(31) Willemsen, V.; Bauch, M.; Bennett, T.; Campilho, A.; Wolkenfelt, H.; Xu, J.; Haseloff, J.; Scheres, B. The NAC Domain Transcription Factors FEZ and SOMBRERO Control the Orientation of Cell Division Plane in *Arabidopsis* Root Stem Cells. Dev. Cell 2008, 15 (6), 913–922. 10.1016/j.devcel.2008.09.019.

(32) Zhang, T.; Ge, Y.; Cai, G.; Pan, X.; Xu, L. WOX-ARF Modules Initiate Different Types of Roots. Cell Rep. 2023, 42 (8), 112966. 10.1016/j.celrep.2023.112966.

(33) Su, Y. H.; Zhao, X. Y.; Liu, Y. B.; Zhang, C. L.; O’Neill, S. D.; Zhang, X. S. Auxin-Induced WUS Expression Is Essential for Embryonic Stem Cell Renewal during Somatic Embryogenesis in Arabidopsis. *Plant J*. Cell Mol. Biol. 2009, 59 (3), 448–460. 10.1111/j.1365-313X.2009.03880.x.

(34) Beeckman, T.; De Rycke, R.; Viane, R.; Inzé, D. Histological Study of Seed Coat Development in Arabidopsis Thaliana. J. Plant Res. 2000, 113 (2), 139–148. 10.1007/PL00013924.

(35) Francoz, E.; Ranocha, P.; Pernot, C.; Le Ru, A.; Pacquit, V.; Dunand, C.; Burlat, V. Complementarity of Medium-Throughput in Situ RNA Hybridization and Tissue-Specific Transcriptomics: Case Study of Arabidopsis Seed Development Kinetics. Sci. Rep. 2016, 6, 24644. 10.1038/srep24644.

(36) Mizzotti, C.; Ezquer, I.; Paolo, D.; Rueda-Romero, P.; Guerra, R. F.; Battaglia, R.; Rogachev, I.; Aharoni, A.; Kater, M. M.; Caporali, E.; Colombo, L. SEEDSTICK Is a Master Regulator of Development and Metabolism in the Arabidopsis Seed Coat. PLoS Genet. 2014, 10 (12), e1004856. 10.1371/journal.pgen.1004856.

(37) Windsor, J. B.; Symonds, V. V.; Mendenhall, J.; Lloyd, A. M. Arabidopsis Seed Coat Development: Morphological Differentiation of the Outer Integument. *Plant J*. Cell Mol. Biol. 2000, 22 (6), 483–493. 10.1046/j.1365-313x.2000.00756.x.

(38) Creff, A.; Brocard, L.; Ingram, G. A Mechanically Sensitive Cell Layer Regulates the Physical Properties of the Arabidopsis Seed Coat. Nat. Commun. 2015, 6, 6382. 10.1038/ncomms7382.

(39) Nakaune, S.; Yamada, K.; Kondo, M.; Kato, T.; Tabata, S.; Nishimura, M.; Hara-Nishimura, I. A Vacuolar Processing Enzyme, δVPE, Is Involved in Seed Coat Formation at the Early Stage of Seed Development. Plant Cell 2005, 17 (3), 876–887. 10.1105/tpc.104.026872.

(40) Muller, B. Characterization of UmamiTs in Arabidopsis: Amino Acid Transporters Involved in Amino Acid Cycling, Phloem Unloading and the Supply of Symplasmically Isolated Sink Tissues. Doctoral defense, Universität Regensburg, 2016. https://epub.uni-regensburg.de/34834/1/Doktorarbeit%20Benedikt%20M%C3%BCller.pdf.

(41) Sanden, N. C. H.; Kanstrup, C.; Crocoll, C.; Schulz, A.; Nour-Eldin, H. H.; Halkier, B. A.; Xu, D. An UMAMIT-GTR Transporter Cascade Controls Glucosinolate Seed Loading in Arabidopsis. Nat. Plants 2024, 10 (1), 172–179. 10.1038/s41477-023-01598-4.

(42) Vogiatzaki, E.; Baroux, C.; Jung, J.-Y.; Poirier, Y. PHO1 Exports Phosphate from the Chalazal Seed Coat to the Embryo in Developing *Arabidopsis* Seeds. Curr. Biol. 2017, 27 (19), 2893–2900.e3. 10.1016/j.cub.2017.08.026.

(43) Li, M.; Li, Q.; Li, S.; Niu, X.; Xu, H.; Li, P.; Bian, X.; Chen, Z.; Liu, Q.; Zhang, H.; Liu, Y.; Wu, S. SHORT-ROOT Specifically Functions in the Chalazal Region to Modulate Assimilate Partitioning into Seeds. *Plant J*. Cell Mol. Biol. 2024, 120 (5), 2031–2044. 10.1111/tpj.17096.

(44) Horák, J.; Grefen, C.; Berendzen, K. W.; Hahn, A.; Stierhof, Y.-D.; Stadelhofer, B.; Stahl, M.; Koncz, C.; Harter, K. The Arabidopsis Thaliana Response Regulator ARR22 Is a Putative AHP Phospho-Histidine Phosphatase Expressed in the Chalaza of Developing Seeds. BMC Plant Biol. 2008, 8, 77. 10.1186/1471-2229-8-77.

(45) Lu, J.; Magnani, E. Seed Tissue and Nutrient Partitioning, a Case for the Nucellus. Plant Reprod. 2018, 31 (3), 309–317. 10.1007/s00497-018-0338-1.

(46) Xu, W.; Fiume, E.; Coen, O.; Pechoux, C.; Lepiniec, L.; Magnani, E. Endosperm and Nucellus Develop Antagonistically in Arabidopsis Seeds. Plant Cell 2016, 28 (6), 1343–1360. 10.1105/tpc.16.00041.

(47) Lu, J.; Le Hir, R.; Gómez-Páez, D.-M.; Coen, O.; Péchoux, C.; Jasinski, S.; Magnani, E. The Nucellus: Between Cell Elimination and Sugar Transport. Plant Physiol. 2021, 185 (2), 478–490. 10.1093/plphys/kiaa045.

(48) Xu, W.; Gomez-Paez, D.-M.; Choinard, S.; Iannaccone, M.; Maricchiolo, E.; Peaucelle, A.; Voxeur, A.; Haas, K. T.; Pompa, A.; Magnani, E. A Change in the Cell Wall Status Initiates the Elimination of the Nucellus in Arabidopsis. bioRxiv April 29, 2024, p 2024.04.23.590775. 10.1101/2024.04.23.590775.

(49) Jullien, P. E.; Grob, S.; Marchais, A.; Pumplin, N.; Chevalier, C.; Bonnet, D. M. V.; Otto, C.; Schott, G.; Voinnet, O. Functional Characterization of Arabidopsis ARGONAUTE 3 in Reproductive Tissues. *Plant J*. Cell Mol. Biol. 2020, 103 (5), 1796–1809. 10.1111/tpj.14868.

(50) Veley, K. M.; Maksaev, G.; Frick, E. M.; January, E.; Kloepper, S. C.; Haswell, E. S. Arabidopsis MSL10 Has a Regulated Cell Death Signaling Activity That Is Separable from Its Mechanosensitive Ion Channel Activity. Plant Cell 2014, 26 (7), 3115–3131. 10.1105/tpc.114.128082.

(51) Müller, B.; Fastner, A.; Karmann, J.; Mansch, V.; Hoffmann, T.; Schwab, W.; Suter-Grotemeyer, M.; Rentsch, D.; Truernit, E.; Ladwig, F.; Bleckmann, A.; Dresselhaus, T.; Hammes, U. Z. Amino Acid Export in Developing Arabidopsis Seeds Depends on UmamiT Facilitators. Curr. Biol. CB 2015, 25 (23), 3126–3131. 10.1016/j.cub.2015.10.038.

(52) Chen, L.-Q.; Lin, I. W.; Qu, X.-Q.; Sosso, D.; McFarlane, H. E.; Londoño, A.; Samuels, A. L.; Frommer, W. B. A Cascade of Sequentially Expressed Sucrose Transporters in the Seed Coat and Endosperm Provides Nutrition for the Arabidopsis Embryo. Plant Cell 2015, 27 (3), 607–619. 10.1105/tpc.114.134585.

(53) Pinto, S. C.; Leong, W. H.; Tan, H.; McKee, L.; Prevost, A.; Ma, C.; Shirley, N. J.; Petrella, R.; Yang, X.; Koltunow, A. M.; Bulone, V.; Kanaoka, M. M.; Higashyiama, T.; Coimbra, S.; Tucker, M. R. Germline β-1,3-Glucan Deposits Are Required for Female Gametogenesis in Arabidopsis Thaliana. Nat. Commun. 2024, 15 (1), 5875. 10.1038/s41467-024-50143-0.

(54) Liu, X.; Nakajima, K. P.; Adhikari, P. B.; Wu, X.; Zhu, S.; Okada, K.; Kagenishi, T.; Kurotani, K.; Ishida, T.; Nakamura, M.; Sato, Y.; Kawakatsu, Y.; Xie, L.; Huang, C.; He, J.; Yokawa, K.; Sawa, S.; Higashiyama, T.; Bradford, K. J.; Notaguchi, M.; Kasahara, R. D. Fertilization-Dependent Phloem End Gate Regulates Seed Size. Curr. Biol. 2025, 0 (0). 10.1016/j.cub.2025.03.033.

(55) Ponnu, J.; Wahl, V.; Schmid, M. Trehalose-6-Phosphate: Connecting Plant Metabolism and Development. Front. Plant Sci. 2011, 2, 70. 10.3389/fpls.2011.00070.

(56) Lunn, J. E.; Feil, R.; Hendriks, J. H. M.; Gibon, Y.; Morcuende, R.; Osuna, D.; Scheible, W.-R.; Carillo, P.; Hajirezaei, M.-R.; Stitt, M. Sugar-Induced Increases in Trehalose 6-Phosphate Are Correlated with Redox Activation of ADPglucose Pyrophosphorylase and Higher Rates of Starch Synthesis in Arabidopsis Thaliana. Biochem. J. 2006, 397 (1), 139–148. 10.1042/BJ20060083.

(57) Eastmond, P. J. SUGAR-DEPENDENT1 Encodes a Patatin Domain Triacylglycerol Lipase That Initiates Storage Oil Breakdown in Germinating Arabidopsis Seeds. Plant Cell 2006, 18 (3), 665–675. 10.1105/tpc.105.040543.

(58) Gómez, L. D.; Gilday, A.; Feil, R.; Lunn, J. E.; Graham, I. A. AtTPS1-Mediated Trehalose 6-Phosphate Synthesis Is Essential for Embryogenic and Vegetative Growth and Responsiveness to ABA in Germinating Seeds and Stomatal Guard Cells. *Plant J*. Cell Mol. Biol. 2010, 64 (1), 1–13. 10.1111/j.1365-313X.2010.04312.x.

(59) Jiang, W.-B.; Huang, H.-Y.; Hu, Y.-W.; Zhu, S.-W.; Wang, Z.-Y.; Lin, W.-H. Brassinosteroid Regulates Seed Size and Shape in Arabidopsis. Plant Physiol. 2013, 162 (4), 1965–1977. 10.1104/pp.113.217703.

(60) Cai, H.; Liu, L.; Huang, Y.; Zhu, W.; Qi, J.; Xi, X.; Aslam, M.; Dresselhaus, T.; Qin, Y. Brassinosteroid Signaling Regulates Female Germline Specification in Arabidopsis. Curr. Biol. CB 2022, 32 (5), 1102–1114.e5. 10.1016/j.cub.2022.01.022.

(61) Liao, K.; Peng, Y.-J.; Yuan, L.-B.; Dai, Y.-S.; Chen, Q.-F.; Yu, L.-J.; Bai, M.-Y.; Zhang, W.-Q.; Xie, L.-J.; Xiao, S. Brassinosteroids Antagonize Jasmonate-Activated Plant Defense Responses through BRI1-EMS-SUPPRESSOR1 (BES1). Plant Physiol. 2020, 182 (2), 1066–1082. 10.1104/pp.19.01220.

(62) Lima, R. B.; Figueiredo, D. D. Sex on Steroids: How Brassinosteroids Shape Reproductive Development in Flowering Plants. Plant Cell Physiol. 2024, 65 (10), 1581–1600. 10.1093/pcp/pcae050.

(63) Jia, D.; Chen, L.-G.; Yin, G.; Yang, X.; Gao, Z.; Guo, Y.; Sun, Y.; Tang, W. Brassinosteroids Regulate Outer Ovule Integument Growth in Part via the Control of INNER NO OUTER by BRASSINOZOLE-RESISTANT Family Transcription Factors. J. Integr. Plant Biol. 2020, 62 (8), 1093–1111. 10.1111/jipb.12915.

(64) Vogler, F.; Schmalzl, C.; Englhart, M.; Bircheneder, M.; Sprunck, S. Brassinosteroids Promote Arabidopsis Pollen Germination and Growth. Plant Reprod. 2014, 27 (3), 153–167. 10.1007/s00497-014-0247-x.

(65) Lima, R. B.; Pankaj, R.; Ehlert, S. T.; Finger, P.; Fröhlich, A.; Bayle, V.; Landrein, B.; Sampathkumar, A.; Figueiredo, D. D. Seed Coat-Derived Brassinosteroid Signaling Regulates Endosperm Development. Nat. Commun. 2024, 15 (1), 9352. 10.1038/s41467-024-53671-x.

(66) Chen, W.; Lv, M.; Wang, Y.; Wang, P.-A.; Cui, Y.; Li, M.; Wang, R.; Gou, X.; Li, J. BES1 Is Activated by EMS1-TPD1-SERK1/2-Mediated Signaling to Control Tapetum Development in Arabidopsis Thaliana. Nat. Commun. 2019, 10 (1), 4164. 10.1038/s41467-019-12118-4.

(67) Yin, Y.; Wang, Z. Y.; Mora-Garcia, S.; Li, J.; Yoshida, S.; Asami, T.; Chory, J. BES1 Accumulates in the Nucleus in Response to Brassinosteroids to Regulate Gene Expression and Promote Stem Elongation. Cell 2002, 109 (2), 181–191. 10.1016/s0092-8674(02)00721-3.

(68) Zhu, T.; Wei, C.; Yu, Y.; Zhang, Z.; Zhu, J.; Liang, Z.; Song, X.; Fu, W.; Cui, Y.; Wang, Z.-Y.; Li, C. The BAS Chromatin Remodeler Determines Brassinosteroid-Induced Transcriptional Activation and Plant Growth in Arabidopsis. Dev. Cell 2024, 59 (7), 924–939.e6. 10.1016/j.devcel.2024.01.021.

(69) Wang, W.; Lu, X.; Li, L.; Lian, H.; Mao, Z.; Xu, P.; Guo, T.; Xu, F.; Du, S.; Cao, X.; Wang, S.; Shen, H.; Yang, H.-Q. Photoexcited CRYPTOCHROME1 Interacts with Dephosphorylated BES1 to Regulate Brassinosteroid Signaling and Photomorphogenesis in Arabidopsis. Plant Cell 2018, 30 (9), 1989–2005. 10.1105/tpc.17.00994.

(70) Oh, E.; Zhu, J.-Y.; Ryu, H.; Hwang, I.; Wang, Z.-Y. TOPLESS Mediates Brassinosteroid-Induced Transcriptional Repression through Interaction with BZR1. Nat. Commun. 2014, 5, 4140. 10.1038/ncomms5140.

(71) Wang, T.; Li, M.; Yang, J.; Li, M.; Zhang, Z.; Gao, H.; Wang, C.; Tian, H. Brassinosteroid Transcription Factor BES1 Modulates Nitrate Deficiency by Promoting NRT2.1 and NRT2.2 Transcription in Arabidopsis. *Plant J*. Cell Mol. Biol. 2023, 114 (6), 1443–1457. 10.1111/tpj.16203.

(72) O’Malley, R. C.; Huang, S.-S. C.; Song, L.; Lewsey, M. G.; Bartlett, A.; Nery, J. R.; Galli, M.; Gallavotti, A.; Ecker, J. R. Cistrome and Epicistrome Features Shape the Regulatory DNA Landscape. Cell 2016, 165 (5), 1280–1292. 10.1016/j.cell.2016.04.038.

(73) Doll, N. M.; Bovio, S.; Gaiti, A.; Marsollier, A.-C.; Chamot, S.; Moussu, S.; Widiez, T.; Ingram, G. The Endosperm-Derived Embryo Sheath Is an Anti-Adhesive Structure That Facilitates Cotyledon Emergence during Germination in Arabidopsis. Curr. Biol. CB 2020, 30 (5), 909–915.e4. 10.1016/j.cub.2019.12.057.

(74) Moussu, S.; Doll, N. M.; Chamot, S.; Brocard, L.; Creff, A.; Fourquin, C.; Widiez, T.; Nimchuk, Z. L.; Ingram, G. ZHOUPI and KERBEROS Mediate Embryo/Endosperm Separation by Promoting the Formation of an Extracuticular Sheath at the Embryo Surface. Plant Cell 2017, 29 (7), 1642–1656. 10.1105/tpc.17.00016.

(75) Yang, S.; Johnston, N.; Talideh, E.; Mitchell, S.; Jeffree, C.; Goodrich, J.; Ingram, G. The Endosperm-Specific ZHOUPI Gene of Arabidopsis Thaliana Regulates Endosperm Breakdown and Embryonic Epidermal Development. Dev. Camb. Engl. 2008, 135 (21), 3501–3509. 10.1242/dev.026708.

(76) Doll, N. M.; Van Hautegem, T.; Schilling, N.; De Rycke, R.; De Winter, F.; Fendrych, M.; Nowack, M. K. Endosperm Cell Death Promoted by NAC Transcription Factors Facilitates Embryo Invasion in Arabidopsis. Curr. Biol. CB 2023, 33 (17), 3785–3795.e6. 10.1016/j.cub.2023.08.003.

(77) Olsen, O.-A. Nuclear Endosperm Development in Cereals and Arabidopsis Thaliana. Plant Cell 2004, 16 *Suppl* (Suppl), S214–227. 10.1105/tpc.017111.

(78) Povilus, R. A.; Gehring, M. Maternal-Filial Transfer Structures in Endosperm: A Nexus of Nutritional Dynamics and Seed Development. Curr. Opin. Plant Biol. 2022, 65, 102121. 10.1016/j.pbi.2021.102121.

(79) Le, B. H.; Cheng, C.; Bui, A. Q.; Wagmaister, J. A.; Henry, K. F.; Pelletier, J.; Kwong, L.; Belmonte, M.; Kirkbride, R.; Horvath, S.; Drews, G. N.; Fischer, R. L.; Okamuro, J. K.; Harada, J. J.; Goldberg, R. B. Global Analysis of Gene Activity during Arabidopsis Seed Development and Identification of Seed-Specific Transcription Factors. Proc. Natl. Acad. Sci. 2010, 107 (18), 8063–8070. 10.1073/pnas.1003530107.

(80) Baroux, C.; Fransz, P.; Grossniklaus, U. Nuclear Fusions Contribute to Polyploidization of the Gigantic Nuclei in the Chalazal Endosperm of Arabidopsis. Planta 2004, 220 (1), 38–46. 10.1007/s00425-004-1326-2.

(81) Ali, M. F.; Shin, J. M.; Fatema, U.; Kurihara, D.; Berger, F.; Yuan, L.; Kawashima, T. Cellular Dynamics of Coenocytic Endosperm Development in Arabidopsis Thaliana. Nat. Plants 2023, 9 (2), 330–342. 10.1038/s41477-022-01331-7.

(82) Murphy, E.; Smith, S.; De Smet, I. Small Signaling Peptides in Arabidopsis Development: How Cells Communicate Over a Short Distance. Plant Cell 2012, 24 (8), 3198–3217. 10.1105/tpc.112.099010.

(83) Matsubayashi, Y. Posttranslationally Modified Small-Peptide Signals in Plants. Annu. Rev. Plant Biol. 2014, 65 (Volume 65, 2014), 385–413. 10.1146/annurev-arplant-050312-120122.

(84) Higashiyama, T. Peptide Signaling in Pollen–Pistil Interactions. Plant Cell Physiol. 2010, 51 (2), 177–189. 10.1093/pcp/pcq008.

(85) Marshall, E.; Costa, L. M.; Gutierrez-Marcos, J. Cysteine-Rich Peptides (CRPs) Mediate Diverse Aspects of Cell–Cell Communication in Plant Reproduction and Development. J. Exp. Bot. 2011, 62 (5), 1677–1686. 10.1093/jxb/err002.

(86) Olsson, V.; Joos, L.; Zhu, S.; Gevaert, K.; Butenko, M. A.; Smet, I. D. Look Closely, the Beautiful May Be Small: Precursor-Derived Peptides in Plants. Annu. Rev. Plant Biol. 2019, 70 (Volume 70, 2019), 153–186. 10.1146/annurev-arplant-042817-040413.

(87) Takahashi, F.; Hanada, K.; Kondo, T.; Shinozaki, K. Hormone-like Peptides and Small Coding Genes in Plant Stress Signaling and Development. Curr. Opin. Plant Biol. 2019, 51, 88–95. 10.1016/j.pbi.2019.05.011.

(88) Ghorbani, S.; Lin, Y.-C.; Parizot, B.; Fernandez, A.; Njo, M. F.; Van de Peer, Y.; Beeckman, T.; Hilson, P. Expanding the Repertoire of Secretory Peptides Controlling Root Development with Comparative Genome Analysis and Functional Assays. J. Exp. Bot. 2015, 66 (17), 5257–5269. 10.1093/jxb/erv346.

(89) Hellmann, E. MtSSPdb: A New Database for the Small Secreted Peptide Research Community. Plant Physiol. 2020, 183 (1), 31–32. 10.1104/pp.20.00376.

(90) Hu, X.-L.; Lu, H.; Hassan, M. M.; Zhang, J.; Yuan, G.; Abraham, P. E.; Shrestha, H. K.; Villalobos Solis, M. I.; Chen, J.-G.; Tschaplinski, T. J.; Doktycz, M. J.; Tuskan, G. A.; Cheng, Z.-M. (Max); Yang, X. Advances and Perspectives in Discovery and Functional Analysis of Small Secreted Proteins in Plants. Hortic. Res. 2021, 8 (1), 1–14. 10.1038/s41438-021-00570-7.

(91) Mergner, J.; Frejno, M.; List, M.; Papacek, M.; Chen, X.; Chaudhary, A.; Samaras, P.; Richter, S.; Shikata, H.; Messerer, M.; Lang, D.; Altmann, S.; Cyprys, P.; Zolg, D. P.; Mathieson, T.; Bantscheff, M.; Hazarika, R. R.; Schmidt, T.; Dawid, C.; Dunkel, A.; Hofmann, T.; Sprunck, S.; Falter-Braun, P.; Johannes, F.; Mayer, K. F. X.; Jürgens, G.; Wilhelm, M.; Baumbach, J.; Grill, E.; Schneitz, K.; Schwechheimer, C.; Kuster, B. Mass-Spectrometry-Based Draft of the Arabidopsis Proteome. Nature 2020, 579 (7799), 409–414. 10.1038/s41586-020-2094-2.

(92) Takeuchi, H.; Higashiyama, T. A Species-Specific Cluster of Defensin-Like Genes Encodes Diffusible Pollen Tube Attractants in Arabidopsis. PLOS Biol. 2012, 10 (12), e1001449. 10.1371/journal.pbio.1001449.

(93) Coculo, D.; Lionetti, V. The Plant Invertase/Pectin Methylesterase Inhibitor Superfamily. Front. Plant Sci. 2022, 13. 10.3389/fpls.2022.863892.

(94) Huang, J.; Zhang, T.; Linstroth, L.; Tillman, Z.; Otegui, M. S.; Owen, H. A.; Zhao, D. Control of Anther Cell Differentiation by the Small Protein Ligand TPD1 and Its Receptor EMS1 in Arabidopsis. PLOS Genet. 2016, 12 (8), e1006147. 10.1371/journal.pgen.1006147.

(95) Song, X.-F.; Hou, X.-L.; Liu, C.-M. CLE Peptides: Critical Regulators for Stem Cell Maintenance in Plants. Planta 2021, 255 (1), 5. 10.1007/s00425-021-03791-1.

(96) Geist, K. S.; Strassmann, J. E.; Queller, D. C. Family Quarrels in Seeds and Rapid Adaptive Evolution in Arabidopsis. Proc. Natl. Acad. Sci. 2019, 116 (19), 9463–9468. 10.1073/pnas.1817733116.

(97) Yang, Z. PAML 4: Phylogenetic Analysis by Maximum Likelihood. Mol. Biol. Evol. 2007, 24 (8), 1586–1591. 10.1093/molbev/msm088.

(98) Shahan, R.; Hsu, C.-W.; Nolan, T. M.; Cole, B. J.; Taylor, I. W.; Greenstreet, L.; Zhang, S.; Afanassiev, A.; Vlot, A. H. C.; Schiebinger, G.; Benfey, P. N.; Ohler, U. A Single-Cell *Arabidopsis* Root Atlas Reveals Developmental Trajectories in Wild-Type and Cell Identity Mutants. Dev. Cell 2022, 57 (4), 543–560.e9. 10.1016/j.devcel.2022.01.008.

(99) Ke, Y.; Pujol, V.; Staut, J.; Pollaris, L.; Seurinck, R.; Eekhout, T.; Grones, C.; Saura-Sanchez, M.; Van Bel, M.; Vuylsteke, M.; Ariani, A.; Liseron-Monfils, C.; Vandepoele, K.; Saeys, Y.; De Rybel, B. A Single-Cell and Spatial Wheat Root Atlas with Cross-Species Annotations Delineates Conserved Tissue-Specific Marker Genes and Regulators. Cell Rep. 2025, 44 (2), 115240. 10.1016/j.celrep.2025.115240.

(100) Guillotin, B.; Rahni, R.; Passalacqua, M.; Mohammed, M. A.; Xu, X.; Raju, S. K.; Ramírez, C. O.; Jackson, D.; Groen, S. C.; Gillis, J.; Birnbaum, K. D. A Pan-Grass Transcriptome Reveals Patterns of Cellular Divergence in Crops. Nature 2023, 617 (7962), 785–791. 10.1038/s41586-023-06053-0.

(101) Lee, T. A.; Illouz-Eliaz, N.; Nobori, T.; Xu, J.; Jow, B.; Nery, J. R.; Ecker, J. R. A Single-Cell, Spatial Transcriptomic Atlas of the Arabidopsis Life Cycle. Nat. Plants 2025, 11 (9), 1960–1975. 10.1038/s41477-025-02072-z.

(102) Guo, X.; Wang, Y.; Zhao, C.; Tan, C.; Yan, W.; Xiang, S.; Zhang, D.; Zhang, H.; Zhang, M.; Yang, L.; Yan, M.; Xie, P.; Wang, Y.; Li, L.; Fang, D.; Guang, X.; Shao, W.; Wang, F.; Wang, H.; Sahu, S. K.; Liu, M.; Wei, T.; Peng, Y.; Qiu, Y.; Peng, T.; Zhang, Y.; Ni, X.; Xu, Z.; Lu, H.; Li, Z.; Yang, H.; Wang, E.; Lisby, M.; Liu, H.; Guo, H.; Xu, X. An Arabidopsis Single-Nucleus Atlas Decodes Leaf Senescence and Nutrient Allocation. Cell 2025, 0 (0). 10.1016/j.cell.2025.03.024.

(103) Zhang, X.; Luo, Z.; Marand, A. P.; Yan, H.; Jang, H.; Bang, S.; Mendieta, J. P.; Minow, M. A. A.; Schmitz, R. J. A Spatially Resolved Multi-Omic Single-Cell Atlas of Soybean Development. Cell 2025, 188 (2), 550–567.e19. 10.1016/j.cell.2024.10.050.

(104) Ali, M. F.; Shin, J. M.; Fatema, U.; Kurihara, D.; Berger, F.; Yuan, L.; Kawashima, T. Cellular Dynamics of Coenocytic Endosperm Development in Arabidopsis Thaliana. Nat. Plants 2023, 9 (2), 330–342. 10.1038/s41477-022-01331-7.

(105) Nguyen, H.; Brown, R. C.; Lemmon, B. E. The Specialized Chalazal Endosperm inArabidopsis Thaliana andLepidium Virginicum (Brassicaceae). Protoplasma 2000, 212 (1), 99–110. 10.1007/BF01279351.

(106) Vriens, K.; Cammue, B. P. A.; Thevissen, K. Antifungal Plant Defensins: Mechanisms of Action and Production. Molecules 2014, 19 (8), 12280–12303. 10.3390/molecules190812280.

(107) Singleton, M. D.; Eisen, M. B. Evolutionary Analyses of Intrinsically Disordered Regions Reveal Widespread Signals of Conservation. PLoS Comput. Biol. 2024, 20 (4), e1012028. 10.1371/journal.pcbi.1012028.

## Methods-Only References

(108) Cheng, C.-Y.; Krishnakumar, V.; Chan, A. P.; Thibaud-Nissen, F.; Schobel, S.; Town, C. D. Araport11: A Complete Reannotation of the Arabidopsis Thaliana Reference Genome. Plant J. 2017, 89 (4), 789–804. 10.1111/tpj.13415.

(109) Young, M. D.; Behjati, S. SoupX Removes Ambient RNA Contamination from Droplet-Based Single-Cell RNA Sequencing Data. GigaScience 2020, 9 (12), giaa151. 10.1093/gigascience/giaa151.

(110) Hao, Y.; Stuart, T.; Kowalski, M. H.; Choudhary, S.; Hoffman, P.; Hartman, A.; Srivastava, A.; Molla, G.; Madad, S.; Fernandez-Granda, C.; Satija, R. Dictionary Learning for Integrative, Multimodal and Scalable Single-Cell Analysis. Nat. Biotechnol. 2024, 42 (2), 293–304. 10.1038/s41587-023-01767-y.

(111) Heumos, L.; Schaar, A. C.; Lance, C.; Litinetskaya, A.; Drost, F.; Zappia, L.; Lücken, M. D.; Strobl, D. C.; Henao, J.; Curion, F.; Schiller, H. B.; Theis, F. J. Best Practices for Single-Cell Analysis across Modalities. Nat. Rev. Genet. 2023, 1–23. 10.1038/s41576-023-00586-w.

(112) Germain, P.-L.; Lun, A.; Garcia Meixide, C.; Macnair, W.; Robinson, M. D. Doublet Identification in Single-Cell Sequencing Data Using scDblFinder. F1000Research 2022, 10, 979. 10.12688/f1000research.73600.2.

(113) Korsunsky, I.; Millard, N.; Fan, J.; Slowikowski, K.; Zhang, F.; Wei, K.; Baglaenko, Y.; Brenner, M.; Loh, P.; Raychaudhuri, S. Fast, Sensitive and Accurate Integration of Single-Cell Data with Harmony. Nat. Methods 2019, 16 (12), 1289–1296. 10.1038/s41592-019-0619-0.

(114) Menges, M.; de Jager, S. M.; Gruissem, W.; Murray, J. A. H. Global Analysis of the Core Cell Cycle Regulators of Arabidopsis Identifies Novel Genes, Reveals Multiple and Highly Specific Profiles of Expression and Provides a Coherent Model for Plant Cell Cycle Control. Plant J. Cell Mol. Biol. 2005, 41 (4), 546–566. 10.1111/j.1365-313X.2004.02319.x.

(115) bluster. Bioconductor. http://bioconductor.org/packages/bluster/ (accessed 2025-04-29).

(116) Xu, S.; Hu, E.; Cai, Y.; Xie, Z.; Luo, X.; Zhan, L.; Tang, W.; Wang, Q.; Liu, B.; Wang, R.;Xie, W. Using clusterProfiler to Characterise Multi-Omics Data. Nat. Protoc. 2024, 19 (11), 3292–3320.

(117) Tirosh, I.; Izar, B.; Prakadan, S. M.; Wadsworth, M. H.; Treacy, D.; Trombetta, J. J.; Rotem, A.; Rodman, C.; Lian, C.; Murphy, G.; Fallahi-Sichani, M.; Dutton-Regester, K.; Lin, J.-R.; Cohen, O.; Shah, P.; Lu, D.; Genshaft, A. S.; Hughes, T. K.; Ziegler, C. G. K.; Kazer, S. W.; Gaillard, A.; Kolb, K. E.; Villani, A.-C.; Johannessen, C. M.; Andreev, A. Y.; Van Allen, E. M.; Bertagnolli, M.; Sorger, P. K.; Sullivan, R. J.; Flaherty, K. T.; Frederick, D. T.; Jané-Valbuena, J.; Yoon, C. H.; Rozenblatt-Rosen, O.; Shalek, A. K.; Regev, A.; Garraway, L. A. Dissecting the Multicellular Ecosystem of Metastatic Melanoma by Single-Cell RNA-Seq. Science 2016, 352 (6282), 189–196. 10.1126/science.aad0501.

(118) Cao, J.; Spielmann, M.; Qiu, X.; Huang, X.; Ibrahim, D. M.; Hill, A. J.; Zhang, F.; Mundlos, S.; Christiansen, L.; Steemers, F. J.; Trapnell, C.; Shendure, J. The Single-Cell Transcriptional Landscape of Mammalian Organogenesis. Nature 2019, 566 (7745), 496–502. 10.1038/s41586-019-0969-x.

(119) Ritchie, M. E.; Phipson, B.; Wu, D.; Hu, Y.; Law, C. W.; Shi, W.; Smyth, G. K. Limma Powers Differential Expression Analyses for RNA-Sequencing and Microarray Studies. Nucleic Acids Res. 2015, 43 (7), e47. 10.1093/nar/gkv007.

(120) Huang, T.; Guillotin, B.; Rahni, R.; Birnbaum, K. D.; Wagner, D. A Rapid and Sensitive, Multiplex, Whole Mount RNA Fluorescence in Situ Hybridization and Immunohistochemistry Protocol. Plant Methods 2023, 19, 131. 10.1186/s13007-023-01108-9.

(121) Teufel, F.; Almagro Armenteros, J. J.; Johansen, A. R.; Gíslason, M. H.; Pihl, S. I.; Tsirigos, K. D.; Winther, O.; Brunak, S.; von Heijne, G.; Nielsen, H. SignalP 6.0 Predicts All Five Types of Signal Peptides Using Protein Language Models. Nat. Biotechnol. 2022, 40 (7), 1023–1025. 10.1038/s41587-021-01156-3.

(122) Positive, Neutral, Negative Selection with Codeml using Multiple Genome Annotations. Bioinformatics Workbook. https://bioinformaticsworkbook.org/dataAnalysis/ComparativeGenomics/Finding_Positive_Selection_With_Codeml.html (accessed 2025-04-29).

(123) Petersen, L.; Bollback, J. P.; Dimmic, M.; Hubisz, M.; Nielsen, R. Genes under Positive Selection in Escherichia Coli. Genome Res. 2007, 17 (9), 1336–1343. 10.1101/gr.6254707.

(124) Emms, D. M.; Kelly, S. OrthoFinder: Phylogenetic Orthology Inference for Comparative Genomics. Genome Biol. 2019, 20 (1), 238. 10.1186/s13059-019-1832-y.

(125) Sievers, F.; Wilm, A.; Dineen, D.; Gibson, T. J.; Karplus, K.; Li, W.; Lopez, R.; McWilliam, H.; Remmert, M.; Söding, J.; Thompson, J. D.; Higgins, D. G. Fast, Scalable Generation of High-Quality Protein Multiple Sequence Alignments Using Clustal Omega. Mol. Syst. Biol. 2011, 7, 539. 10.1038/msb.2011.75.

(126) Shakya, M.; Ahmed, S. A.; Davenport, K. W.; Flynn, M. C.; Lo, C.-C.; Chain, P. S. G. Standardized Phylogenetic and Molecular Evolutionary Analysis Applied to Species across the Microbial Tree of Life. Sci. Rep. 2020, 10 (1), 1723. 10.1038/s41598-020-58356-1.

