## Supplemental Figures 1-9 for "A transcriptional atlas of early Arabidopsis seed development suggests mechanisms for inter-tissue coordination"

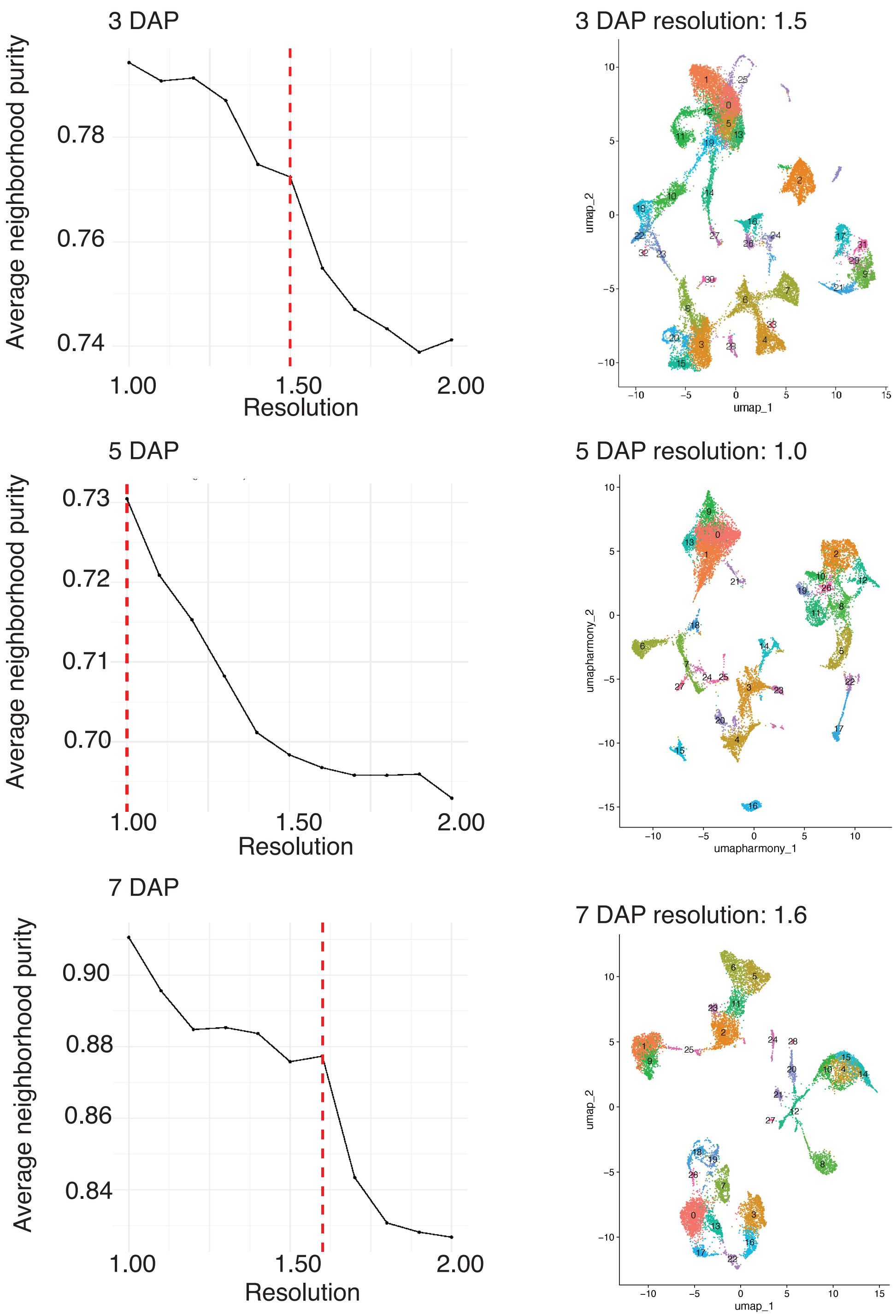


**Supplementary Figure 1. Identifying an initial clustering resolution for level 3 annotations.** Processed snRNA-seq libraries were merged within timepoints and subjected to a parameter sweep of clustering resolutions after initial dimensionality reduction. Each clustering resolution was evaluated by neighborhood purity, which is the percentage of neighbors that belong to the same cluster for each cell based on a weighted calculation. The left column contains the results of this parameter sweep for each timepoint, the red dotted line corresponds to the resolution that was selected for each timepoint. The right column displays the datasets with their *de novo* clusters at the optimal clustering resolution.

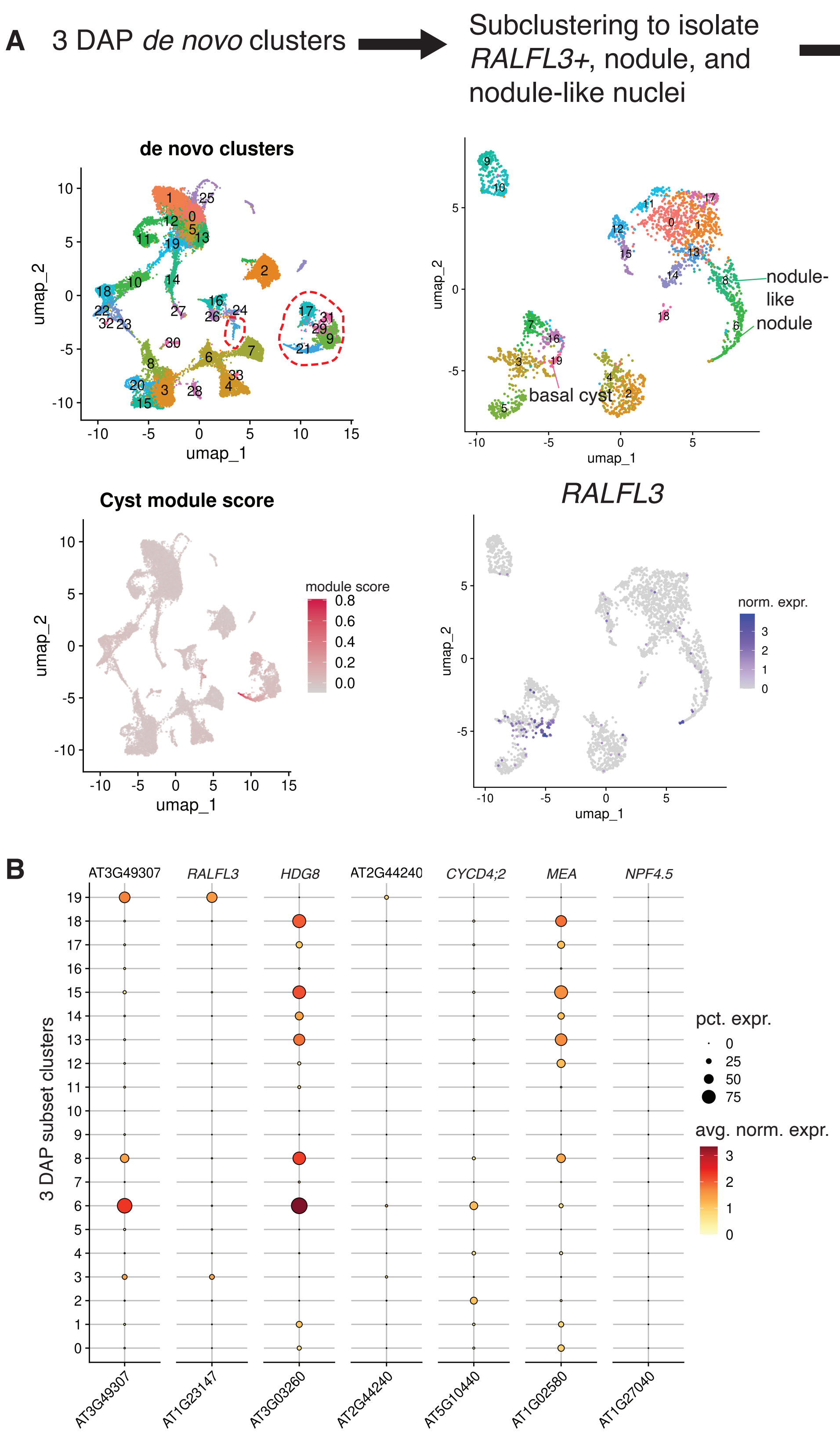


**Supplementary Figure 2. Isolating chalazal endosperm subtypes at 3 DAP through a subclustering analysis.** **A**, Clusters expressing CZE markers from Picard et al. 2021 ("Cyst module") were subset from timepoint datasets and subject to an additional dimensionality reduction and subclustering. Published markers and those validated by HCR were used to annotated the subclusters (**B**), then the nuclei within endosperm clusters were manually labelled in the timepoint object.

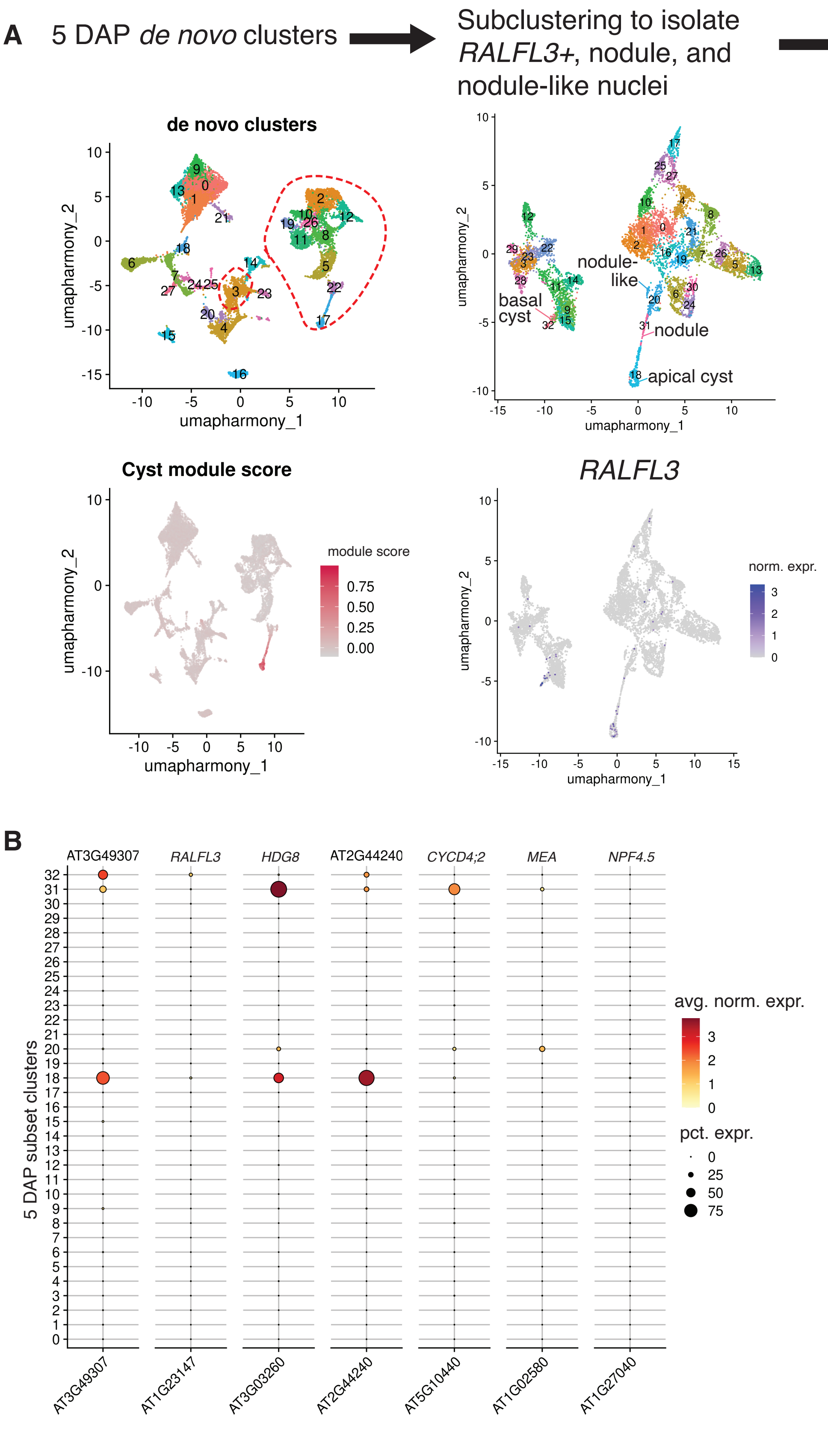


**Supplementary Figure 3. Isolating chalazal endosperm subtypes at 5 DAP through a subclustering analysis.** **A**, **B**, same procedure as described in Supplementary Figure 2, but for 5 DAP CZE nuclei.

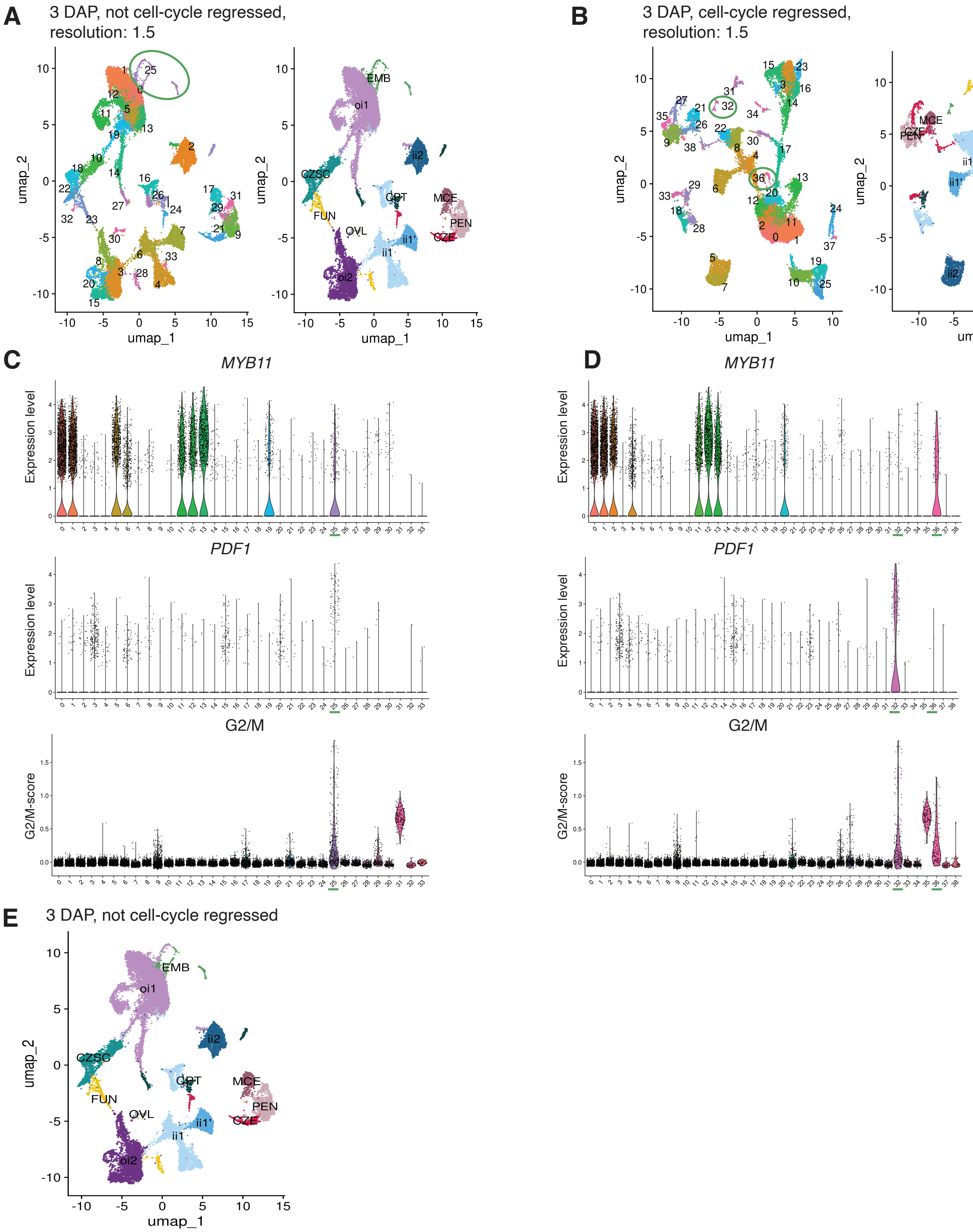


**Supplementary Figure 4. Identifying a contaminating G2/M oi nuclei population in the 3 DAP embryo cluster through a cell cycle regression analysis. A**, *De novo* clusters identified in the 3 DAP dataset (left) and inital L2 annotations (right). The embryo clusters are circled. **B**, Same as in A, but for the 3 DAP dataset after cell cycle regression. The embryo splits into two clusters at the same clustering resolution as A when cell cycle genes are regressed out (32 and 36). **C**, Marker genes used to distinguish oi1 (*MYB11*) and embryo (*PDF1*) nuclei populations (top two panels) and G2/M score in the original 3 DAP dataset, plotted across de novo clusters. The de novo clusters initially annotated as embryo clusters are underlined. One cluster is highly enriched for *MYB11* and *PDF1* expression as well as G2/M score (25). **D**, Same as in C, but for the 3 DAP dataset after cell cycle regression. *MYB11* and *PDF1* are no longer highly expressed in the same cluster. The two clusters originally annotated as embryonic nuclei populations have high G2/M scores. Cluster 32 is likely embryonic, while cluster 36 is a putatively dividing oi1 population. **E**, Updated level 2 annotations after nuclei in cluster 36 of the cell cycle-regressed dataset were re-annotated as oi1.

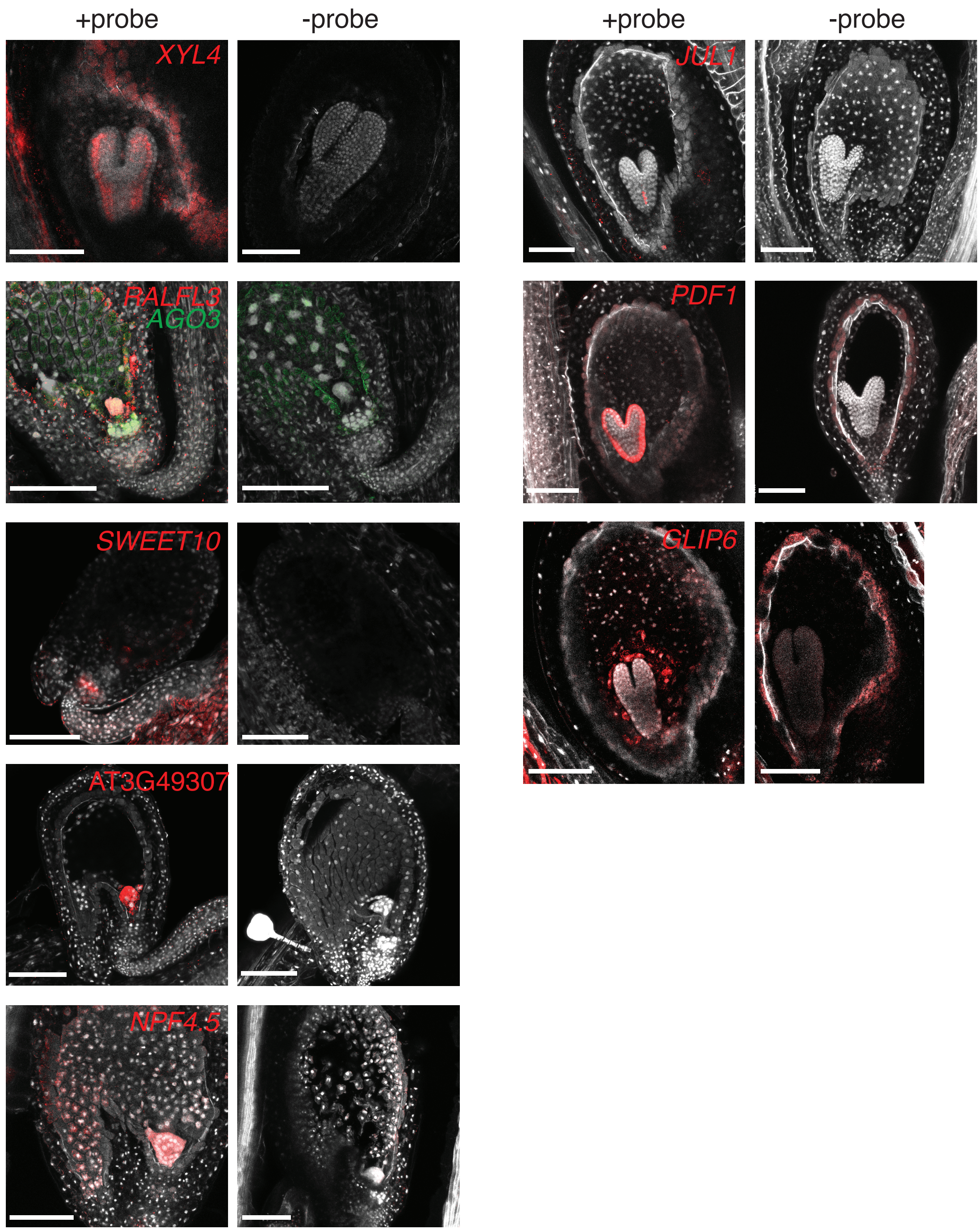


**Supplementary Figure 5. HCR probe validation for markers described in this study.** Negative controls (samples processed through the HCR protocol without mRNA probe application) were generated in parallel with the "+probe" sample shown on the left in each column. Scale bar is 100 $\mu$m. See Supplementary Table 2 for a list of all probes used in this study.

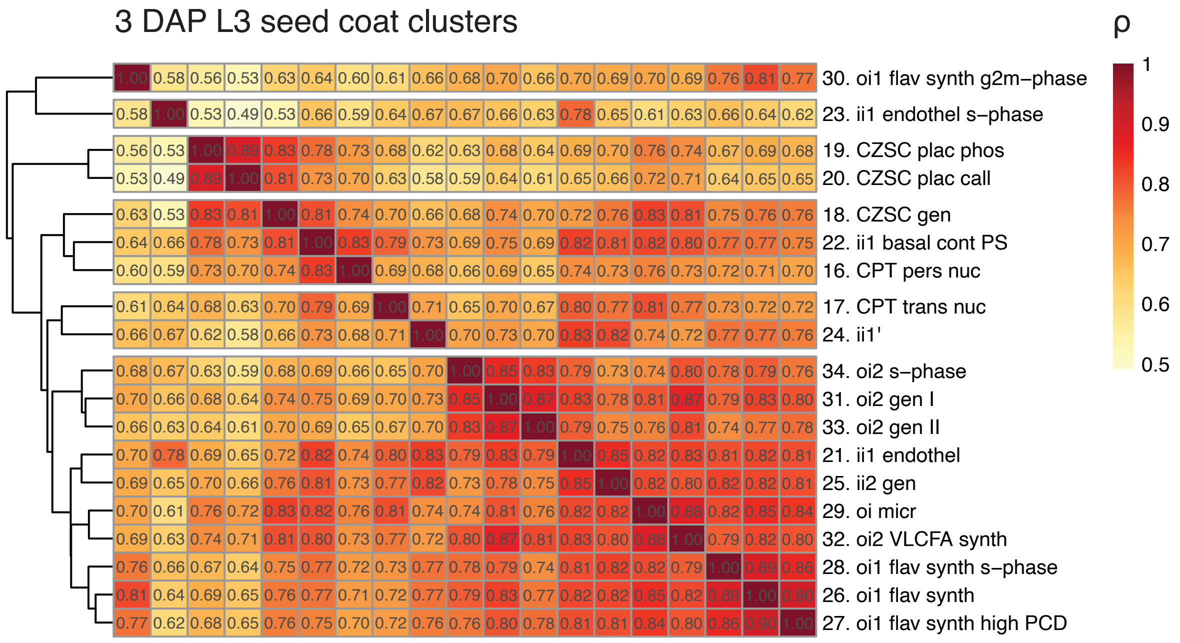


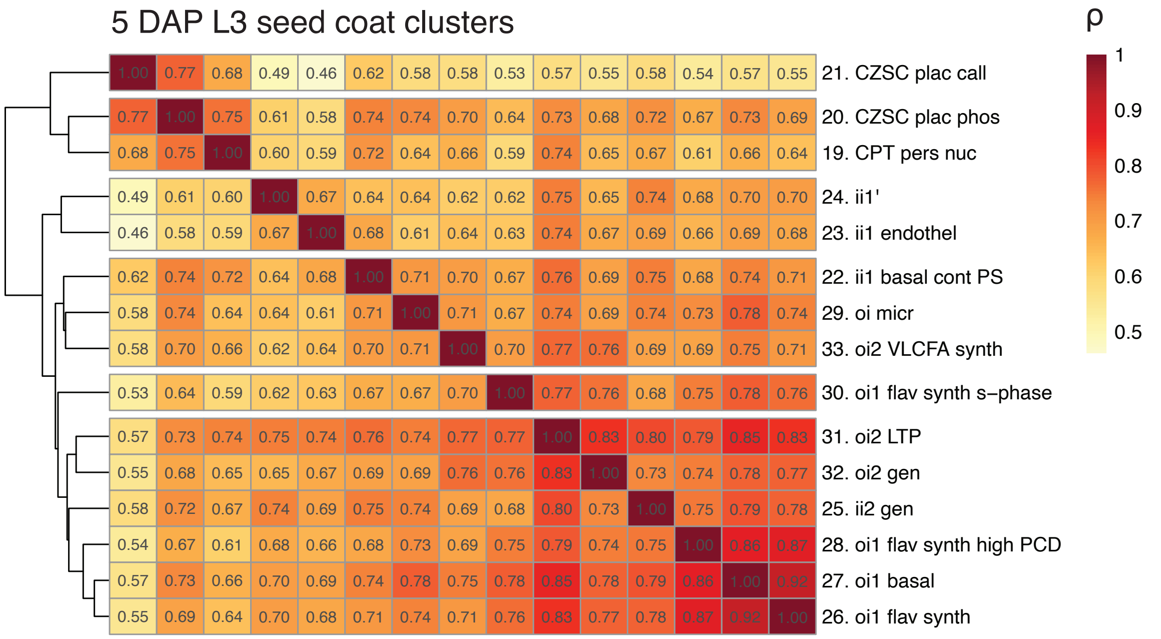


**Supplementary Figure 6. Level 3 seed coat cluster similarity at 3 and 5 DAP.** Clustered heatmaps of the Spearman correlation coefficients from pairwise comparisons of aggregated expression of highly variable genes within L3 clusters. See Supplementary Table 3 to match abbreviated L3 names to their full descriptions.

**A**

secretory signal sequence (SignalP6)

```
TPD1 1    MNRRRLLVSATLLSYLLYGMAL---VSVEASGGEKLRDNLDLTKTTTSSPSISHRKM-----LLLSPGTGKTERSVEPER  72
TDL1 1    MRMEHIYKFQHWLFFIGLGVLLSLSLSVKANDFEGSNDRRSIALKARSFVSINENRTALSRKLLLSPDIGDGTN-----R  75

TPD1 73   IGEKCKSTDIVVNQAVTEPMPNGIPGYMVEITNQCMSGCIISRIHINCGWFSSAKLINPRVFKRIHYDDCLVNNGKPLPF  152
TDL1 76   IGQDCSKDDIVLFQGSTNPLPSGVPSYTVEIFNSCVSDCNIAEIHVSCGWFSSVRLVNPRVFRRLDYDDCLVNDGQPLGP  155

TPD1 153  GSTLSFHYANTFPYHLSVAFVTCA  176
TDL1 156  GQSLSFQYANSFSYPLSVASVSCF  179
```

46% identity overall

59.6% identity excluding secretory signal sequence

**B**

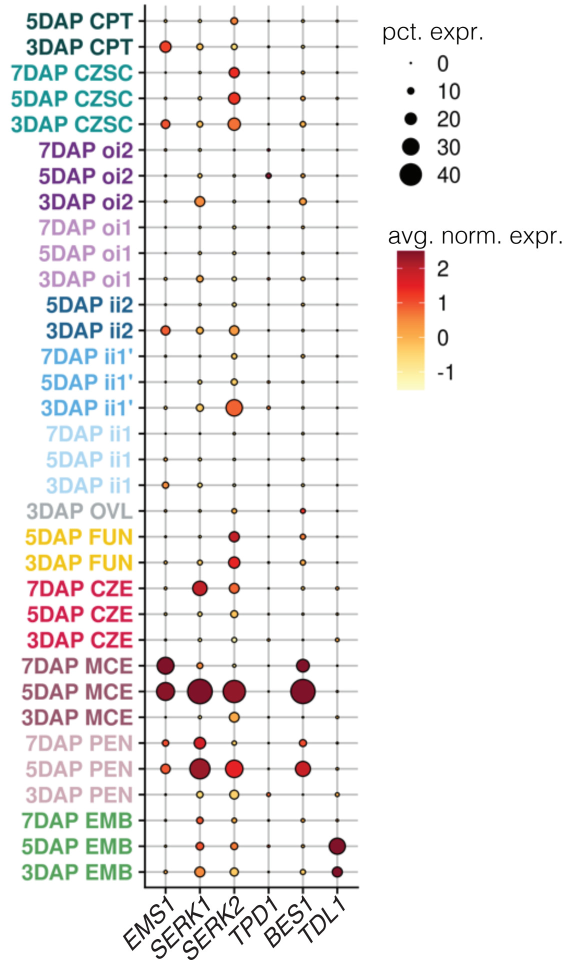


**Supplementary Figure 7. TDL1 is a TPD1-like SSP expressed in the early embryo. A,** Protein sequence alignment for TPD1 and TDL1, the most similar SSP to TPD1 in the Arabidopsis genome by BLAST. The N-terminal regions, which contain secretory signal sequences, are less conserved than the C-terminal regions. **B,** the expression patterns for genes underlying BR-independent acivation of *BES1* through *TPD1,* contrasted with that of *TDL1*.

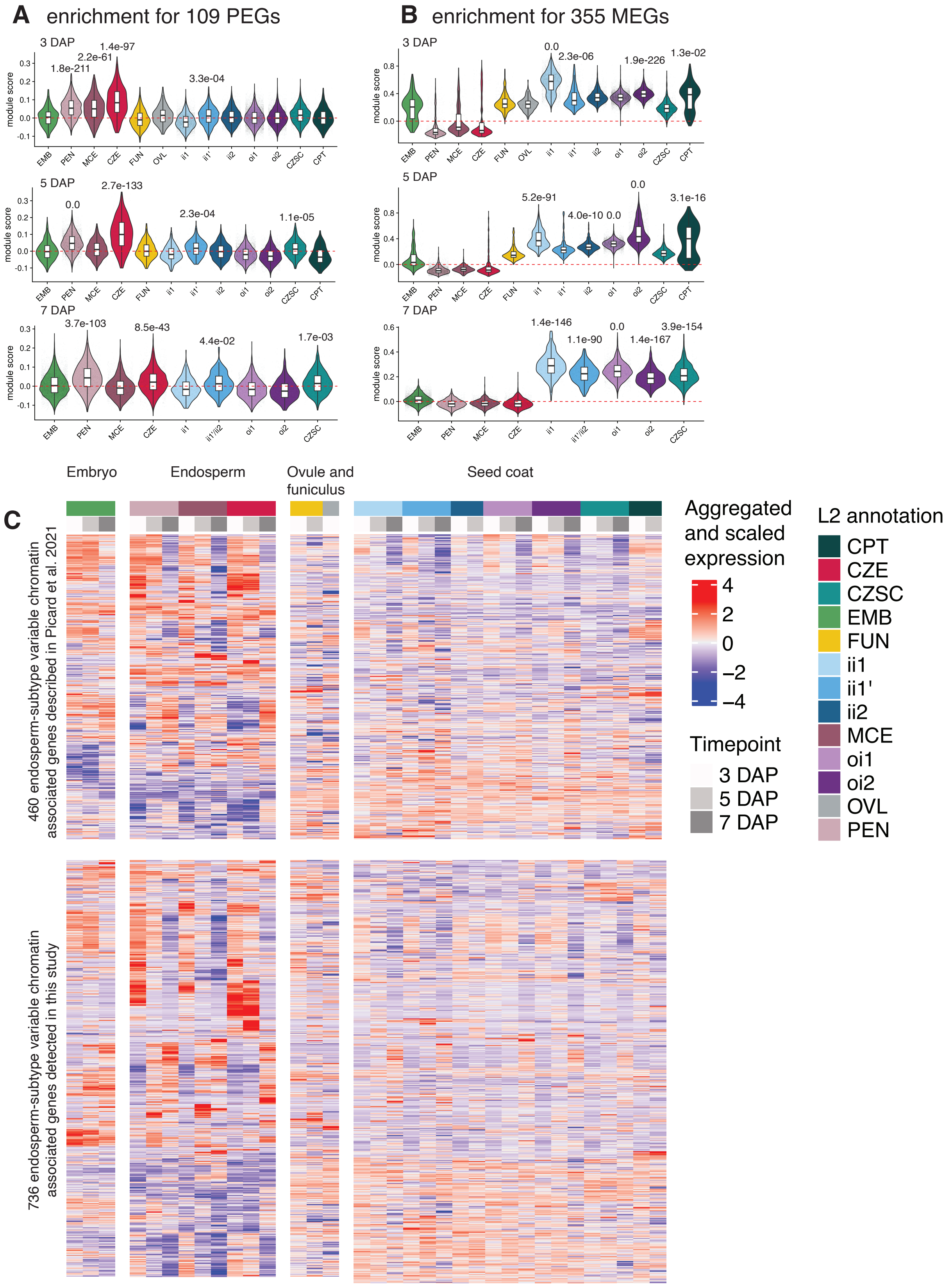


**Supplementary Figure 8. Gene expression enrichment for imprinted genes as well as epigenetic and transcriptional regulators in endosperm subtypes. A-B,** Module score analysis of paternally-expressed genes (PEGs) and maternally-expressed genes (MEGs) defined in Picard et al. 2021 across all timepoints and L2 annotations, respectively. p-values in A and B are centered above clusters with significantly high positive module scores in a cluster-vs-all nuclei comparison. p-values are derived from a two-sided Wilcoxon Rank-Sum test with Bonferroni correction. See Supplementary Table 7 for the module scores and p-values for all clusters. **C**, Top: cluster-aggregated expression patterns for chromatin-assoicated genes that were variably expressed across endosperm subtypes described in Picard et al. 2021. Bottom: cluster-aggregated expression for genes associated with GO terms including chromatin (GO:0000785), epigenetic regulation of gene expression (GO:0040029), and transcriptional regulation of gene expression (GO:0010468) that are differentially expressed across endosperm subtypes in this study (log2FC > 1, adjusted p-value ≤ 0.05). Rows are clustered, columns are not. See Supplementary Table 5 for the differential expression analysis for the genes that underlie this heatmap.

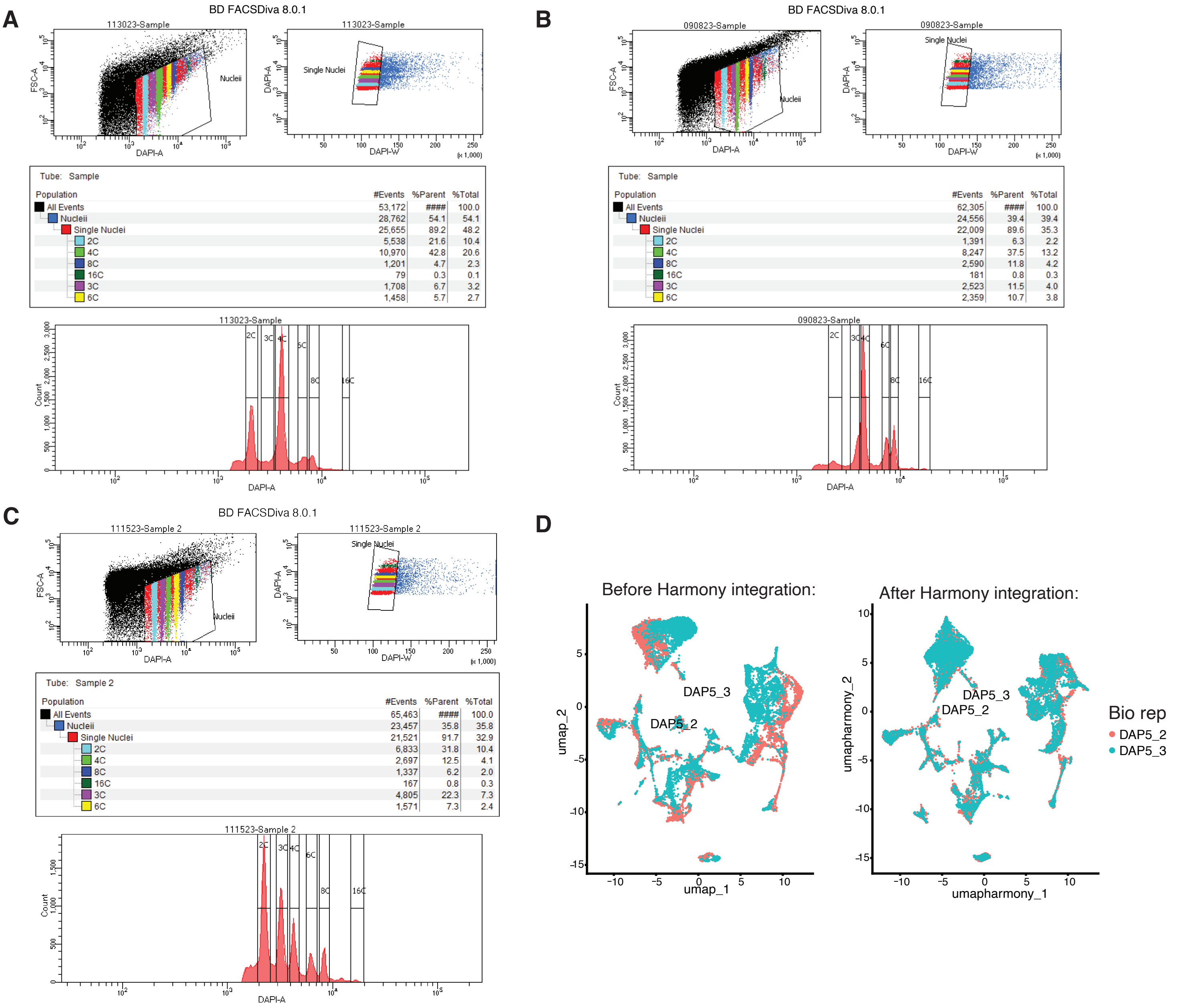


**Supplementary Figure 9. FANS gating scheme for purifying seed nuclei before snRNA-seq and replicate integration. A-C**, Example gating parameters used for one of the 3-7 DAP replicates, respectively, all gates shown were collected. **D**, 5 DAP biological replicates before and after Harmony integration. Biological replicates showed improved mixing after Harmony integration.
